# Precision cooling assists electrical stimulation: Breaking selectivity barriers in nerve control

**DOI:** 10.64898/2026.09.14.751475

**Authors:** Shu Yang, Xiao Yu, Shihao Yang, Qianqian Wang, Guo-Qiang Bi, Yun Liu, Zhanhong Du

## Abstract

Peripheral neuromodulation can relieve pain through sensory activation or conduction block. However, electrical stimulation of mixed nerves couples these intended effects to unwanted neural activity. Here we demonstrate local cooling can separate targeted from off-target neural activity. In the rat sciatic nerve, precooling reduced the onset response to kilohertz-frequency stimulation by about 90% and maintained block in the target pathway during rewarming. During low-frequency stimulation, spatially graded cooling suppressed Aδ-enriched activity and blocked nontarget propagation. This produced a directional, Aβ-biased afferent output. A cooling–electrical neural digital twin predicted both responses and linked them to field-edge transition dynamics and spatially separated control of recruitment and conduction. These findings provide a mechanistically grounded route beyond the targeting limits of electrical stimulation alone.

## Introduction

The continuing burden of opioid-related harms has intensified the search for nonopioid approaches to pain management (*1–3*). Peripheral neuromodulation can relieve pain through sensory activation or conduction block, providing a reversible alternative to systemic pharmacological treatment (*4–11*). In mixed nerves, however, electrical stimulation couples these intended effects to off-target neural activity. Fibers with different physiological functions can be recruited by the same stimulus (*12–14*), while electrically evoked action potentials propagate both proximally and distally (*15–17*), linking sensory activation to motor output and other off-target responses (*7–11, 18–22*). Selective control therefore depends not only on whether neural activity is evoked or blocked, but also on which fibers are recruited and where their activity propagates (*18, 20–22*).

This coupling is difficult to avoid because fibers of different diameters and physiological functions lie close together and experience overlapping electrical fields (*12–14*). Changing current amplitude, waveform, or electrode configuration can shift recruitment and conduction thresholds, but these electrical parameters do not independently control fiber recruitment and action-potential propagation (*15, 17, 18, 20–23*). Two common neuromodulation modes illustrate this limitation (Fig. 1A). Kilohertz-frequency stimulation (KHFS) can rapidly and reversibly block conduction (*4, 5, 9, 24, 25*), yet its initiation often produces an onset response before stable block is established (*15, 16, 26*). Low-frequency stimulation can engage sensory pathways associated with analgesia and plasticity (*7, 8, 10, 11*), but it can also recruit off-target fiber populations and launch activity in both directions (*12–14, 18*). Antidromic recruitment of sensory fibers may further engage peripheral neurogenic inflammatory pathways (*27–30*). Although waveform shaping and multielectrode strategies can improve particular aspects of selectivity (*15, 20, 22, 23, 26*), electrical stimulation alone leaves fiber recruitment and propagation tightly coupled.

**Fig. 1.**
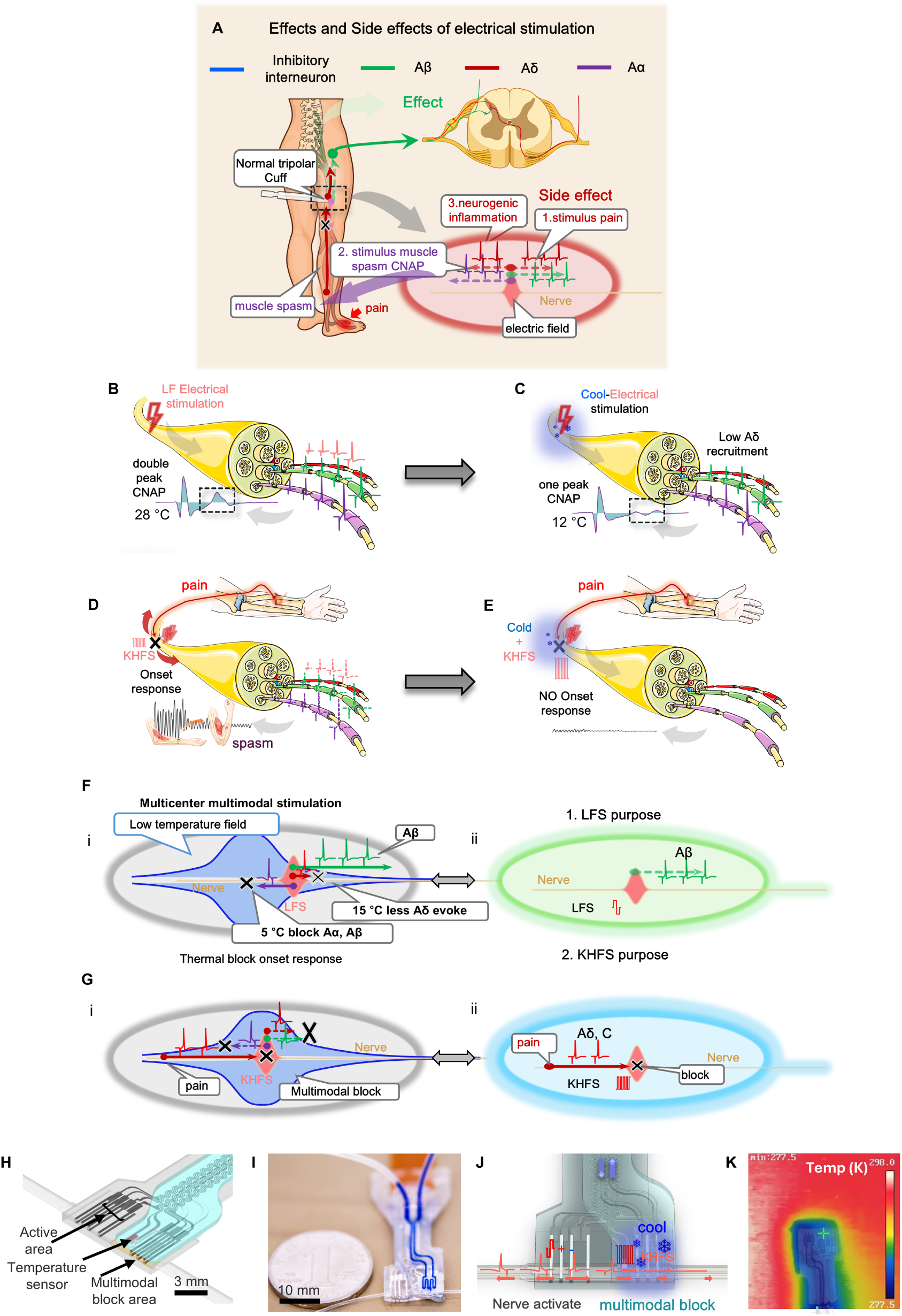
Cooling-assisted electrical control of mixed peripheral nerves with the NeuroSwitch cuff. (A) Schematic of off-target effects caused by nonselective fiber recruitment during KHFS block for acute pain and low-frequency feedback stimulation for chronic pain: muscle spasm through the Aα motor pathway, neurogenic inflammation through antidromic Aδ activity, and pain through orthodromic Aδ activity. (B) Low-frequency electrical stimulation recruits multiple fiber classes in a mixed nerve, producing a double-peaked compound nerve action potential (CNAP) in the rat sciatic nerve at 28 °C. (C) Cooling-assisted low-frequency stimulation at 12 °C preferentially retains the Aα/Aβ-enriched component while reducing Aδ recruitment, consistent with the single-peaked CNAP recorded in vivo. (D) Under the uncooled experimental condition (28 °C), kilohertz-frequency high-frequency stimulation (KHFS) intended for analgesic conduction block elicits an onset response, muscle spasm, and ankle locking; the schematic is accompanied by representative rat ankle images before and after KHFS onset. (E) Under cooling, KHFS produces no marked onset-related muscle spasm or ankle locking. (F) Spatially separated multicenter cooling and low-frequency stimulation generate a cleaner Aβ-biased afferent output. (i) The cooling core (5 °C) blocks non-target-side Aα/Aβ propagation, whereas residual cooling near the stimulation site (15 °C) suppresses Aδ recruitment while retaining part of the Aβ response. (ii) Target output state for low-frequency stimulation. (G) Colocalized cooling and KHFS suppress the KHFS onset response. (i) Because the cooling field extends beyond the electrical block field, precooling suppresses activation in the edge zone. (ii) Target state in which multimodal conduction block interrupts Aδ/C-fiber nociceptive input. (H) Rendering of the final test-device architecture. (I) Photograph of the fabricated test device. (J) Functional layout showing the low-frequency nerve-activation site on the left and the cooling-assisted electrical block site on the right. (K) Infrared image of the device during active cooling.

Separating these processes requires a control variable that changes the physiological state of the nerve and thereby reshapes its response to the applied electric field. Temperature provides such a variable. Cooling alters ion-channel kinetics and membrane excitability (*31–35*), slows action-potential conduction (*6, 36, 37*), and, at sufficiently low temperatures, can reversibly block propagation (*6, 38, 39*). Temperature also shifts the operating windows of both cooling block and kilohertz-frequency stimulation (*19, 39, 40*). These effects could expand the control space of electrical stimulation. Cooling near the stimulation site could reshape fiber recruitment, whereas a cooler region along a propagation pathway could block conduction after activity has been initiated. Spatially structured cooling could therefore separate recruitment from propagation and reshape neural outputs that remain coupled under electrical stimulation alone (Fig. 1F and G).

Using temperature in this way requires more than adding a cooling element to a conventional stimulator. The response of an axon to electrical stimulation depends on local temperature: cooling alters membrane excitability, conduction velocity, and block threshold, so the same extracellular potential can produce different outcomes at different temperatures (*6, 19, 31–37, 39*). Neural activation, propagation, and block consequently depend on how the electric and temperature fields overlap in space and evolve in time. Existing models reproduce many features of electrically evoked neural activity (*22, 41–46*), but mixed-nerve control under local cooling additionally requires temperature-dependent membrane dynamics and spatially resolved predictions of fiber recruitment and conduction. A predictive framework must therefore link the spatial and temporal structure of the cooling and electrical fields to fiber-specific activation, propagation, and block (*47*).

Here, we used local temperature to shift the operating state of mixed nerves and thereby reshape their responses to electrical stimulation. We implemented this principle in NeuroSwitch, a microfluidic cooling–electrical nerve interface (Fig. 1H to K), and paired it with an experimentally informed, multiscale cooling–electrical neural digital twin. Precooling strongly reduced the KHFS onset response and maintained block in the target pathway during rewarming. During low-frequency stimulation, spatially graded cooling preferentially suppressed Aδ-enriched activity near the drive site while the colder core blocked nontarget propagation.

Together, these effects converted bidirectional mixed activity into a directional, Aβ-biased afferent output. The digital twin predicted both response modes and linked KHFS onset suppression to field-edge transition dynamics and low-frequency selectivity to spatially separated control of recruitment and conduction. These results show that local cooling expands electrical control into a joint cooling–electrical space that allows fiber recruitment, propagation, and conduction block to be manipulated with greater independence.

## Results

### NeuroSwitch implements spatially patterned cooling–electrical control

Device development began with a first-generation cuff in which the centers of cooling and electrical stimulation were separated by only 4 mm. During low-frequency stimulation, cooling abolished the evoked afferent compound nerve action potential (CNAP) when the measured core nerve temperature reached approximately 19 °C (fig. S2A to H). This temperature was substantially higher than the sub-10 °C temperatures typically required for direct cold conduction block (*6, 38*), indicating that overlap of the cooling and electrical fields suppressed neural recruitment at the stimulation site before blocking action-potential propagation. This unexpected observation motivated us to spatially separate recruitment from conduction block and to map interference between the drive and block regions (fig. S2J to L).

NeuroSwitch was configured in two spatial arrangements for in vivo cooling–electrical control. In the block–drive configuration, the low-frequency drive was positioned within the warmer portion of the cooling gradient, whereas the colder core lay along the nontarget propagation pathway (Fig. 1F). In the cooling–KHFS configuration, the cooling region was centered on the high-frequency block site, allowing the same nerve segment to be precooled before KHFS (Fig. 1G).

Representative in vivo responses illustrated the effects of cooling in both configurations. At 28 °C, low-frequency stimulation produced a double-peaked CNAP, whereas cooling to 12 °C shifted the response toward a predominantly single-peaked waveform (Fig. 1B and C). Under the uncooled condition, KHFS elicited a pronounced onset-associated motor response, which was markedly attenuated after precooling (Fig. 1D and E).

The flexible NeuroSwitch cuff integrates a microfluidic cooling channel, platinum cuff electrodes, and an embedded temperature sensor for simultaneous local cooling, electrical stimulation, and temperature monitoring (Fig. 1H to J). Infrared thermography confirmed spatially localized cooling around the block region (Fig. 1K). The center-to-center distance between the activation and KHFS block regions exceeded 8 mm, greater than the 7.5-mm no-interference threshold established by multispacing tests (fig. S2J to L). Device fabrication is detailed in fig. S1 and Table S2.

### A multiscale cooling–electrical neural digital twin captures responses to cooling and electrical stimulation

To predict how local temperature reshapes neural responses to electrical stimulation, we established a multiscale cooling–electrical neural digital twin. The model was developed to reproduce experimental observations and quantify the activation, propagation, and conduction-block boundaries of different fibers within a mixed nerve across stimulation currents, temperatures, and spatial configurations.

The digital twin spans three scales: the macroscale nerve–device system, mesoscale nerve fibers, and microscale membrane dynamics. At the macroscale, we reconstructed the implanted device geometry around the rat sciatic nerve (Fig. 2A) and calculated the temperature and electric-field distributions under different stimulation modes (Fig. 2B and C; fig. S3G and H; fig. S4). At the mesoscale, we implemented two complementary fiber models. A fiber reconstructed directly within the finite-element model (Fig. 2J) resolved action-potential propagation in detail, whereas compartmental models of fiber populations (Fig. 2I), with dimensions derived from histological cross-sections (Fig. 2G), simulated recruitment and block across fiber classes and reconstructed CNAPs (Fig. 2H and I; figs. S8 and S11; see Materials and Methods). At the microscale, we extended established ion-channel models to incorporate temperature dependence in membrane potential and the open probabilities of Na_V_1.8, Na_V_1.6, Na_V_1.1, and K_V_1 channels (*34, 35, 48*). We also replaced the passive leak conductance with the voltage- and temperature-sensitive K2P channels TREK-1 and TRAAK (*32, 33, 49, 50*) (Fig. 2K; figs. S9 and S10). Detailed methods for cooling-field simulation, single-fiber modeling, and MRG model construction are provided in the supplementary modeling materials (fig. S3A to E; see Materials and Methods).

**Fig. 2.**
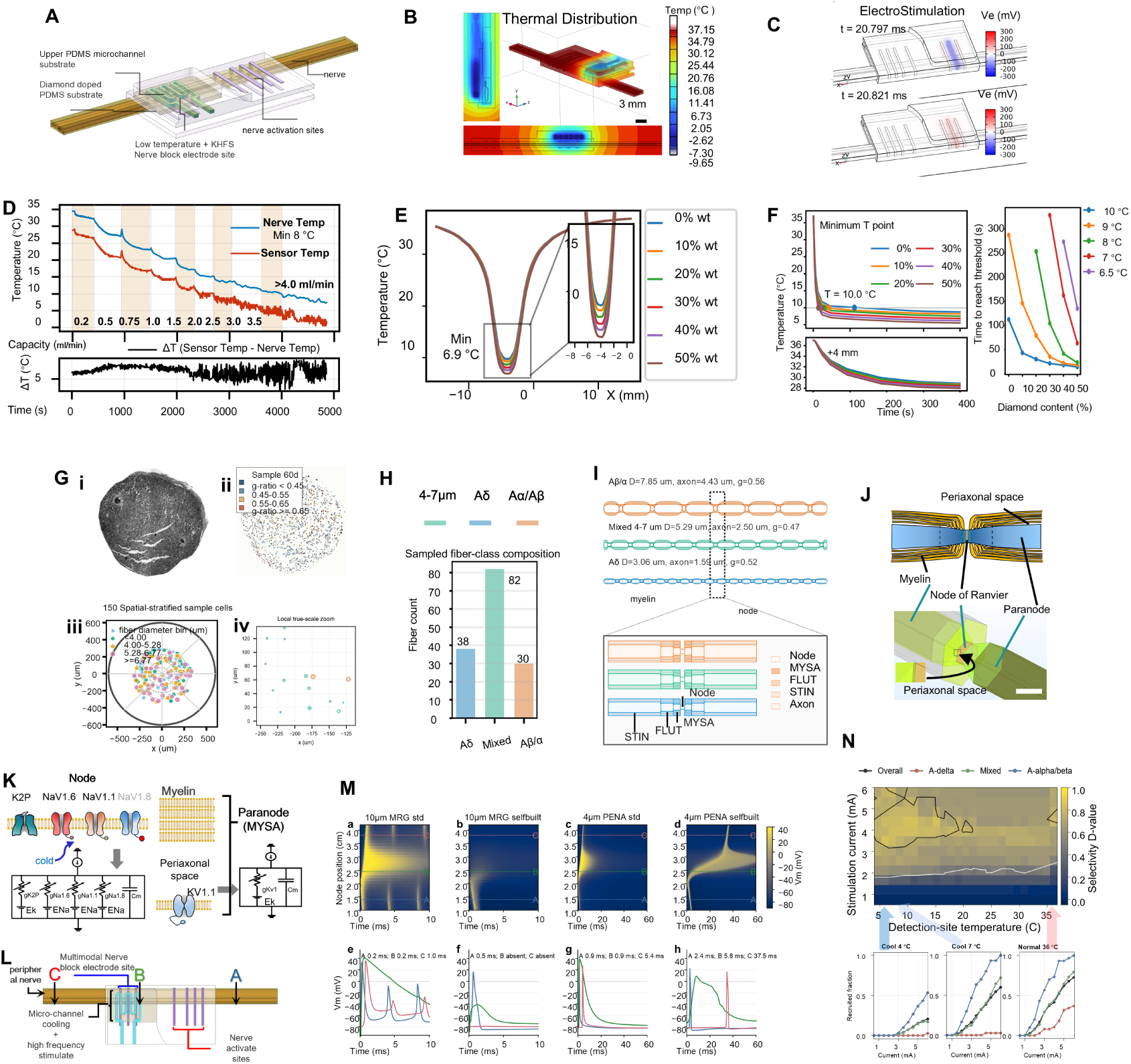
A multiscale cooling-assisted electrical digital twin defines fiber-selective control of mixed nerves. (A) Three-dimensional implanted device-nerve geometry used for coupled simulations. (B) Simulated temperature field and transverse and longitudinal sections under the standard finite-element thermal-simulation condition (flow rate, 4 mL min⁻¹; inlet temperature, −10 °C). (C) Simulated extreme-value extracellular-potential distributions during tripolar stimulation. (D) Intraneural temperature at site 1, sensor temperature at site 3, and their difference (ΔT) during stepwise changes in flow rate; measurement sites are defined in figs. S5A and S6A. (E) Simulated axial temperature profiles from the transient finite-element comparison along the lower nerve surface for devices containing 0 to 50 wt.% diamond; the approximately 7 °C and 10 °C values are transient field-comparison results. (F) Cooling transients at the longitudinal midpoint beneath the microchannel and 4 mm lateral to the channel center, together with the diamond-content dependence of the time required to reach each block temperature. (G) Reconstruction of the mixed-fiber population from histology. (i) Enhanced histological cross-section. (ii) Segmentation of nonadherent fibers and extraction of axon diameter and g-ratio. (iii) Spatially and diameter-stratified sampling of 150 fibers. (iv) Enlarged view of local fiber geometry. (H) Composition of the sampled population: 38 Aδ, 82 mixed-diameter, and 30 Aα/Aβ fibers. (I) Compartmental representations of the modeled fiber classes. (J) Three-dimensional finite-element fiber model and enlarged nodal region. (K) Temperature-coupled ion-channel distributions at nodes, paranodes, and juxtaparanodal myelin. (L) Stimulation, block, and recording sites in the mixed-nerve model. (M) Action-potential initiation and cold-block behavior of standard and temperature-responsive models with matched structures (10-μm MRG and 4-μm PENA fibers). (N) Fiber-class recruitment across the 0.5- to 6.0-mA mixed-population sweep and cooling-center temperature; the representative 6.0-mA point uses the 8.44 °C steady-state minimum-temperature input, distinct from the transient values in (E). The upper map shows selectivity and the lower plots show class-specific recruitment fractions at representative temperatures.

We first characterized the cooling performance and electrochemical properties of the NeuroSwitch cuff. A benchtop platform reproducing the implantation environment was used to measure the spatial extent of cooling and assess the ability of the integrated sensor to track temperature changes at the neural interface (Fig. 2D; figs. S5 and S6A; Table S3). The diamond-filled PDMS interface reduced nerve temperature to 8 °C, below the cooling-block threshold (fig. S3F and G). The embedded thermistor maintained an approximately 5 °C offset from intraneural temperature (Fig. 2D), with negligible delay (fig. S6B). At a fixed cold-trap temperature of −35 °C, nerve temperature decreased exponentially with increasing flow rate (figs. S5H and S6C and E), while heat conduction broadened the cooling profile along the nerve (fig. S6D). Electrochemical testing yielded a cathodal charge-storage capacity of 20.9 mC/cm², an impedance below 100 Ω at 1000 Hz, and a charge-injection capacity of 112 µC/cm², supporting operation within the stimulation conditions used here (fig. S7; Table S4).

Using these benchtop measurements, we constructed a digital twin of nerve cooling by the NeuroSwitch cuff. Parameters for thermally conductive layers containing different diamond fractions (fig. S3D and F) were incorporated into a finite-element model that included metabolic heat generation and blood perfusion (see Materials and Methods) to simulate steady-state and transient cooling. The temperature-field simulations confirmed the importance of the highly conductive diamond-filled silicone layer. Increasing diamond content enhanced cooling depth (Fig. 2E) and markedly shortened the time required to reach block temperature (Fig. 2F). Under otherwise identical conditions (inlet temperature, −10 °C; flow rate, 5 mL min⁻¹), the nerve temperature beneath the microchannel center reached 7 °C with 50 wt.% diamond but 10 °C with pure PDMS (Fig. 2E; fig. S3G), whereas further increases in flow rate did not continuously improve cooling performance (fig. S3H). Diamond loading had an even greater effect on cooling kinetics: reaching a minimum nerve temperature of 9 °C required approximately 20 s with the 50 wt.% device but approximately 290 s with the 0 wt.% device (Fig. 2F).

We next examined why conventional fiber models fail to reproduce cooling-induced action-potential conduction block. The simulated nerve-temperature distribution was imported into the fiber model, and a temperature-dependent coefficient, CP, was used to represent different levels of cooling (fig. S11A and B); action potentials were sampled at the locations shown in Fig. 2L. Conventional models treat temperature only as a scaling factor for ion-channel transition rates. Cooling therefore prolonged channel-gating cycles without reducing their peak amplitudes, paradoxically increasing charge transfer (fig. S10). Action-potential conduction slowed, but physiologically consistent cooling block remained difficult to reproduce (Fig. 2L; Fig. 2M, a and c). We addressed this limitation by introducing temperature dependence into maximal channel conductances and ionic reversal potentials (see Materials and Methods). These modifications reduced charge transfer during each ion-channel cycle (fig. S9) and enabled the models to reproduce cooling-induced conduction block. The revised models captured slowed conduction under mild cooling and reversible block under deeper cooling (Fig. 2M; fig. S13F to H; Tables S7 to S9), providing a basis for mixed-nerve population simulations.

To extend the analysis from individual fibers to the mixed-fiber population, we reconstructed fiber diameter, g-ratio, and spatial distributions from stained rat sciatic-nerve cross-sections and selected 150 representative fibers by stratified sampling across space and diameter (Fig. 2G; fig. S8). Fibers were grouped by diameter into Aδ (<4 μm), mixed-diameter (4 to <7 μm), and Aα/Aβ (≥7 μm) classes (Fig. 2H and I). A compartmental model was constructed for each fiber using its measured position, diameter, g-ratio, and temperature-responsive ion channels (Fig. 2G and I). The resulting population model quantified activation thresholds, propagation states, and block fractions (Fig. 2G, H, and N; figs. S8 and S11; Tables S5 and S6). An initial simulation of temperature-dependent electrical recruitment is shown in Fig. 2N. Although not yet calibrated against experimental data, the model resolved how temperature shifted the recruitment profiles of different fiber classes. The simulations suggested that cooling could enhance preferential Aα/Aβ recruitment by differentially suppressing recruitment across fiber populations. Temperature could therefore reshape fiber recruitment across the current–temperature plane to support additional control objectives. This initial analysis informed the subsequent in vivo experiments, after which ion-channel densities were calibrated to align simulated responses with physiological measurements.

### Cooling suppresses the onset response by reshaping the transition pathway into KHFS block

At physiological temperature, KHFS typically produces a brief onset response before stable conduction block is established. Previous studies have shown that cooling can attenuate this response, but the underlying mechanism and effective spatial boundaries remain unclear. We therefore combined in vivo experiments with the digital twin to examine cooling-mediated onset suppression, test the transition from cooling block to KHFS block, and define the membrane dynamics and spatial relationships that govern combined cooling–KHFS control.

We first tested whether local cooling could suppress the KHFS onset response. Tripolar 10-kHz KHFS was applied intermittently at different temperatures on the rat sciatic nerve, and onset severity was quantified by the ankle-angle deflection at stimulation onset (see Materials and Methods). Near physiological temperature, KHFS evoked pronounced ankle movement and toe contraction. As local temperature decreased, the onset-related ankle deflection declined steadily from approximately 50° to approximately 5° at 10 °C, corresponding to about 90% suppression. The mechanical response recovered after rewarming, indicating that the reduction primarily reflected a reversible, temperature-dependent change in nerve excitability rather than fatigue from repeated stimulation (Fig. 3A and B).

**Fig. 3.**
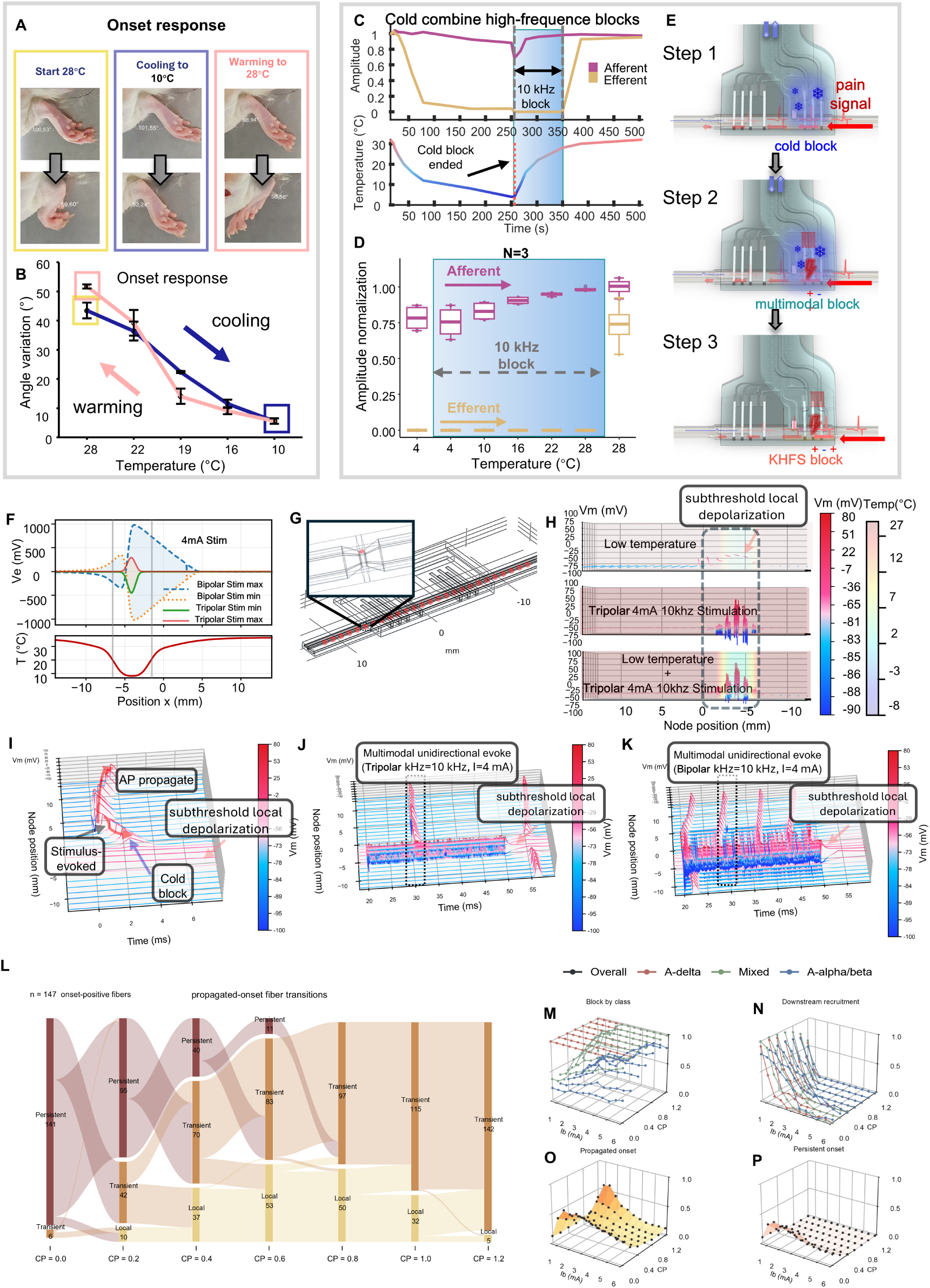
Cooling suppresses the KHFS onset response and enables stable cooling-assisted conduction block. (A) Representative ankle positions at baseline, during cooling, and after rewarming during tripolar KHFS onset testing. (B) KHFS-evoked ankle-angle deflection decreases with cooling and recovers on rewarming. Values are mean ± SD across three rats. (C) Representative afferent and efferent fast-component CNAP amplitudes during cold block, transition to 10-kHz KHFS, and recovery. (D) Across-animal quantification of normalized maximum afferent and efferent CNAP amplitudes during cooling and KHFS maintenance (n = 3 rats). Magenta and gold denote afferent and efferent recordings, respectively, as in (C); box-plot summaries indicate mean ± SEM. The blue shaded region denotes the interval during which KHFS was applied. (E) Three-step onset-suppressed block sequence: cooling first lowers excitability in the block region, cooling and KHFS overlap during handoff, and KHFS then maintains block during rewarming. (F) Axial envelopes of bipolar and tripolar KHFS potentials compared with the cooling-induced temperature profile. (G) Device-to-axon spatial mapping and membrane-potential (*V_m_*) sampling nodes used to generate spatiotemporal maps. (H) Axial *V_m_* and temperature profiles under cooling alone, tripolar KHFS alone (10 kHz, 4 mA), and combined cooling plus tripolar KHFS. (I) Action-potential initiation, propagation, subthreshold depolarization, and cold block under cooling alone. (J) Combined cooling and tripolar KHFS suppress onset while establishing directional block. (K) Combined cooling and bipolar KHFS permits onset activity on both sides of the broader bipolar field. (L) State transitions of the 147 onset-positive fibers as cooling strength increases. Onset positivity was assessed across all KHFS amplitudes, with a fiber classified as positive if onset occurred at any stimulation amplitude; responses were classified as persistent propagated, transient propagated, or local/nonpropagating onset. (M) Fiber-class block fraction across cooling strength and KHFS amplitude. (N) Residual downstream recruitment after combined block. (O) Fraction of fibers with transient propagated onset. (P) Fraction of fibers with persistent propagated onset.

We next examined whether cooling block could transition into KHFS block without transient recovery of conduction. As shown in Fig. 3E, cooling was first applied until the efferent CNAP disappeared. KHFS was then initiated at low temperature, and the nerve was gradually rewarmed while KHFS continued. The efferent CNAP remained suppressed throughout rewarming, whereas the afferent CNAP gradually recovered as temperature increased. Both CNAP directions recovered after KHFS was discontinued (Fig. 3C to E; see Materials and Methods). Thus, conduction block was maintained continuously during the transition from cooling block to KHFS block and throughout subsequent rewarming.

We then used the digital twin to examine why cooling suppresses KHFS onset and how this effect depends on the spatial relationship between the cooling and electric fields. A 10-μm axon was embedded in the full finite-element model, with membrane potentials sampled at successive nodes of Ranvier, to resolve the spatiotemporal extracellular potential, membrane-state changes, and action-potential propagation during tripolar KHFS (Fig. 3G; fig. S12A and B). The simulations localized onset initiation to the edges of the KHFS electric field rather than the center of the steady-state block region (fig. S12E). At these field edges, the high-frequency stimulus was insufficient to establish stable block immediately after stimulation began, yet sufficiently strong to perturb membrane potential and trigger propagating action potentials during the transition from normal conduction to block.

Device-scale field simulations and single-fiber modeling further supported this mechanism. The NeuroSwitch cooling field fully covered both the principal tripolar KHFS electric field and its edge regions, whereas the more diffuse bipolar KHFS field extended partly beyond the cooled region (Fig. 3F; fig. S4). Steady-state membrane-potential analysis further showed that both cooling and KHFS locally depolarized the block region, consistent with previous patch-clamp recordings (*51, 52*) and suggesting that both contribute to conduction block (Fig. 3H; fig. S13A to E). Dynamic simulations showed that KHFS alone produced onset activity before establishing block (fig. S12D to F, H, J, and K), whereas cooling alone established block without onset (Fig. 3I; fig. S12C). Combined cooling and tripolar KHFS likewise blocked conduction without onset (Fig. 3J; fig. S12G, I, and M). Combined cooling and bipolar KHFS still established block (Fig. 3K; fig. S12L), but permitted residual onset activity, primarily at electric-field edges that extended beyond adequate cooling (Fig. 3K; fig. S12E and J). Effective suppression of KHFS onset therefore depended on cooling the electric-field edge regions from which onset activity was most likely to emerge.

Finally, we used the mixed-fiber population model to quantify block and onset during combined cooling and KHFS (see Materials and Methods). At physiological temperature, 147 fibers were onset-positive across the tested stimulation amplitudes. Onset probability progressively decreased with cooling and reached a minimum at CP = 0.8, corresponding to a minimum temperature of approximately 14 °C (Fig. 3L, O, and P; fig. S14). Thus, effective onset suppression did not require cooling to the cooling-block threshold. Population analyses after fiber activation showed that cooling and KHFS acted synergistically to produce conduction block and that both preferentially blocked larger-diameter fibers (Fig. 3M and N; figs. S14B and S15). Cooling also shifted persistent propagated onset responses toward transient or nonpropagating responses, suggesting that electrically evoked activation is more sensitive to cooling than subsequent action-potential propagation.

Together, these findings show that precooling reshapes the transition into KHFS block by reducing local neural excitability, slowing membrane dynamics, and establishing a depolarized local state before KHFS begins. The simulations identify field-edge transition dynamics as a basis for cooling-mediated onset suppression: the effect depends on spatial overlap between the cooling field and the KHFS field edges, where onset activity preferentially emerges. Adequate cooling of these edge regions therefore provides a key spatial criterion for suppressing propagated onset during combined cooling and high-frequency conduction block.

### CNAP reconstruction links the quick and slow components to distinct fiber populations

Before analyzing block–drive output, we determined the fiber populations contributing to the two experimentally observed CNAP components. Electrically evoked CNAPs consistently contained a short-latency N1–P1 component and a delayed N2–P2 component, referred to here as the quick and slow components, respectively (Fig. 4B, C, and G; see Materials and Methods). Based on conduction-velocity differences among fiber classes, we hypothesized that the quick component was enriched in Aα/Aβ activity, whereas the slow component was enriched in Aδ activity. We therefore reconstructed the experimental CNAP with the digital twin and decomposed the modeled waveform according to fiber diameter.

**Fig. 4.**
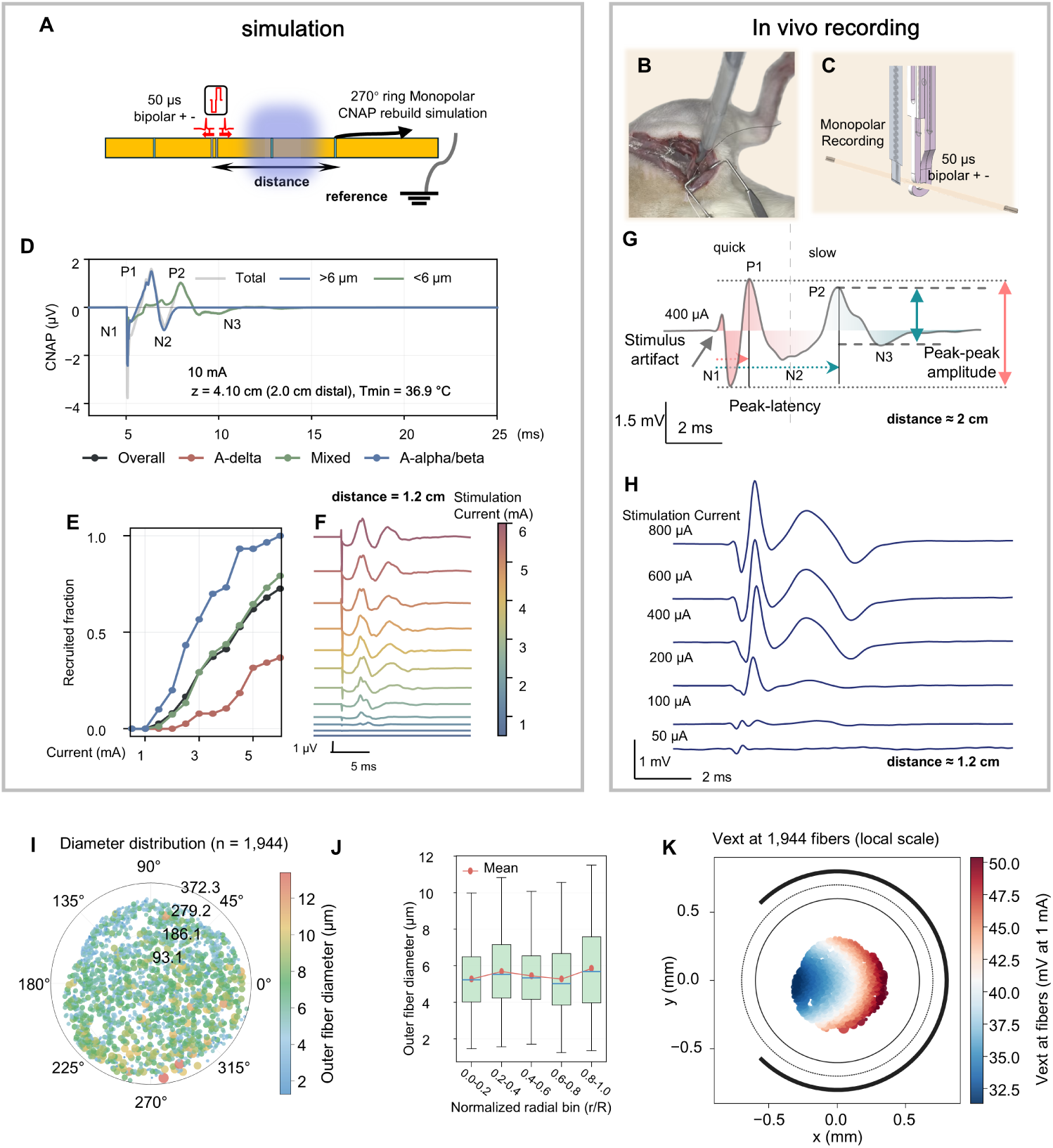
Experimental–computational CNAP reconstruction links quick and slow compound components to fiber-diameter contributions. (A) Simulation and recording scheme using charge-balanced bipolar stimulation and a 270° ring monopolar recording electrode. (B) Intraoperative photograph of the nerve–electrode interface. (C) In vivo bipolar-stimulation and monopolar-recording configuration. (D) Simulated total CNAP and contributions from fibers >6 μm and <6 μm at physiological temperature (10 mA; recording site 2.0 cm distal to stimulation). (E) Simulated recruited fractions of the overall population and the Aδ, mixed-diameter, and Aα/Aβ classes as a function of stimulation current at a 1.2-cm recording distance. (F) Current-dependent simulated CNAPs at the same distance. (G) Representative in vivo CNAP defining the quick and slow components, peak-to-peak amplitude, and peak latency. (H) In vivo CNAPs evoked by 50- to 800-μA stimulation at an approximately 1.2-cm recording distance (n = 3 rats). (I) Outer-fiber-diameter distribution of the reconstructed 1,944-fiber population. (J) Fiber diameter summarized across normalized radial bins. (K) Extracellular potentials sampled at the 1,944 fiber positions beneath the 270° ring electrode.

The model incorporated the in vivo bipolar stimulation geometry, 270° cuff monopolar recording configuration, and sciatic-nerve cross-sectional anatomy to reconstruct CNAPs from the extracellular potentials generated by the modeled fiber population (Fig. 4A to F and I to K; fig. S17). Simulated and experimental waveforms shared key features: both exhibited temporally separated double peaks and similar increases in peak amplitude and waveform evolution with increasing stimulation current (Fig. 4D to H). Agreement in waveform morphology, peak timing, and current dependence indicated that the model captured the fiber recruitment underlying the experimental CNAPs. The model could therefore be used to resolve the fiber contributions to each waveform component. As stimulation current increased from 50 to 800 μA with 50-μs phases, both CNAP components increased in amplitude, but neither the experiments nor the simulations revealed a later peak attributable to C-fiber activity (Fig. 4H; fig. S16).

Decomposition by fiber diameter showed that, using 6 μm as the threshold for waveform decomposition, fibers >6 μm contributed predominantly to the early quick component, whereas fibers <6 μm contributed more strongly to the delayed slow component (Fig. 4D). Together with the fiber classifications used in the population model, these results support interpreting the quick component as Aα/Aβ-enriched activity and the slow component as activity enriched in smaller-diameter Aδ fibers.

Neither the experiments nor the simulations showed complete separation of Aα/Aβ and Aδ recruitment as stimulation current decreased. The model provided an explanation for this overlap. Fiber diameter and g-ratio showed no simple spatial ordering within the reconstructed nerve bundle, whereas the electric field generated by the 270° cuff was spatially nonuniform. Activation thresholds were therefore determined jointly by intrinsic fiber properties and local electric-field strength and did not follow the simple diameter-dependent recruitment order expected for isolated fibers (Fig. 4I to K; fig. S17). The quick and slow CNAP components should therefore be interpreted as readouts enriched in different fiber populations rather than as fiber-specific markers.

Overall, the digital twin reproduced the experimental CNAP in waveform morphology, peak timing, and stimulation-current dependence and resolved the fiber populations contributing to the two components. This correspondence provides the basis for using the quick and slow CNAP components to quantify fiber bias in the subsequent block–drive experiments.

### Cooling-gated block–drive converts bidirectional mixed activity into a directional, Aβ-biased afferent output in vivo

Using the CNAP-component interpretation established above, we next tested whether cooling-gated block–drive could convert the bidirectional mixed activity evoked by low-frequency stimulation into a directional, Aβ-biased afferent output. In NeuroSwitch, the drive site is spatially offset from the cooling center, allowing the temperature gradient to influence both recruitment at the stimulation site and subsequent action-potential propagation. Residual cooling near the drive site can alter fiber recruitment, whereas the colder core along the nontarget pathway can further suppress propagation. To distinguish these effects, we simultaneously recorded afferent and efferent CNAPs on opposite sides of the drive site (Fig. 5A to C; see Materials and Methods).

**Fig. 5.**
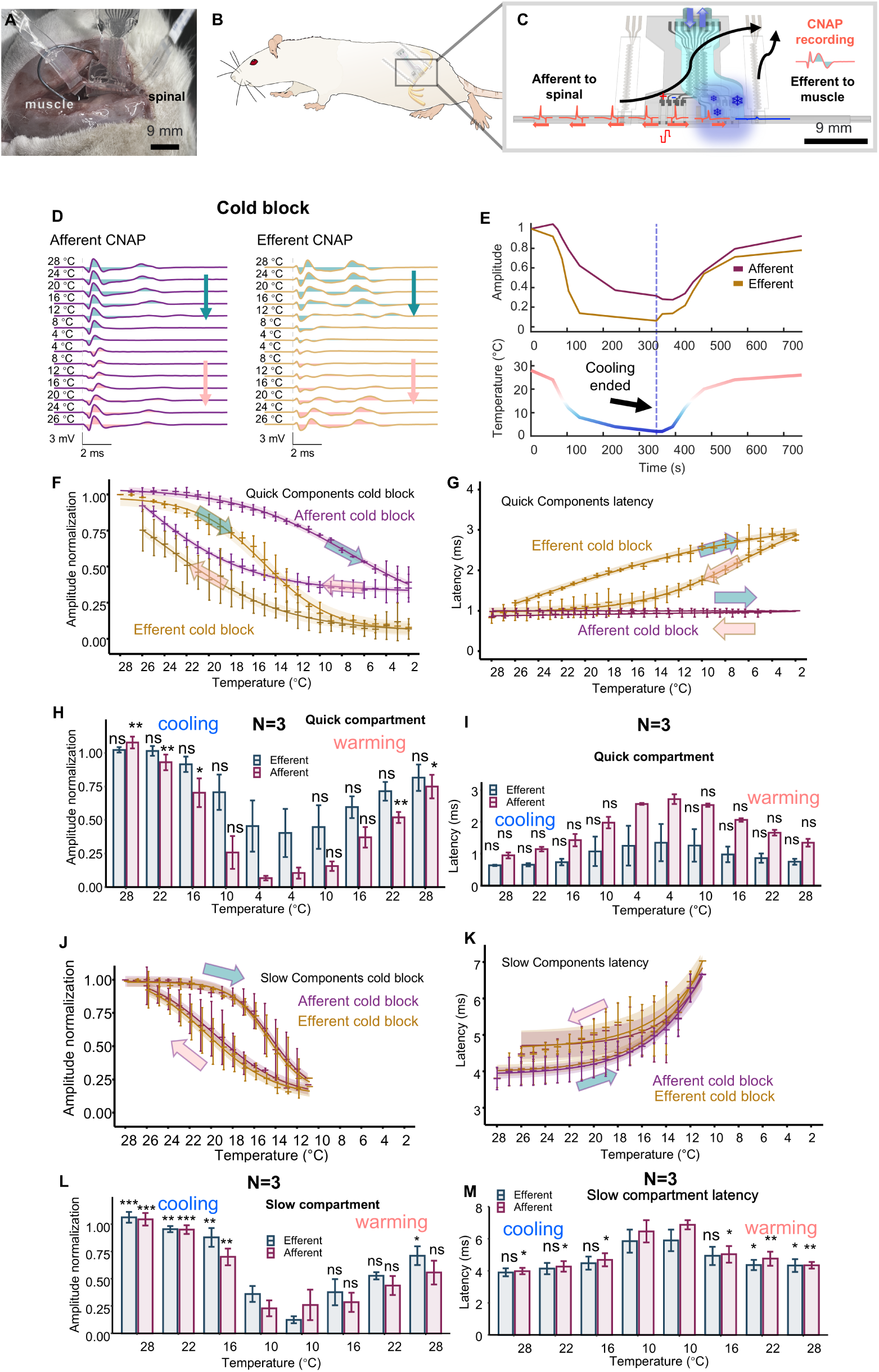
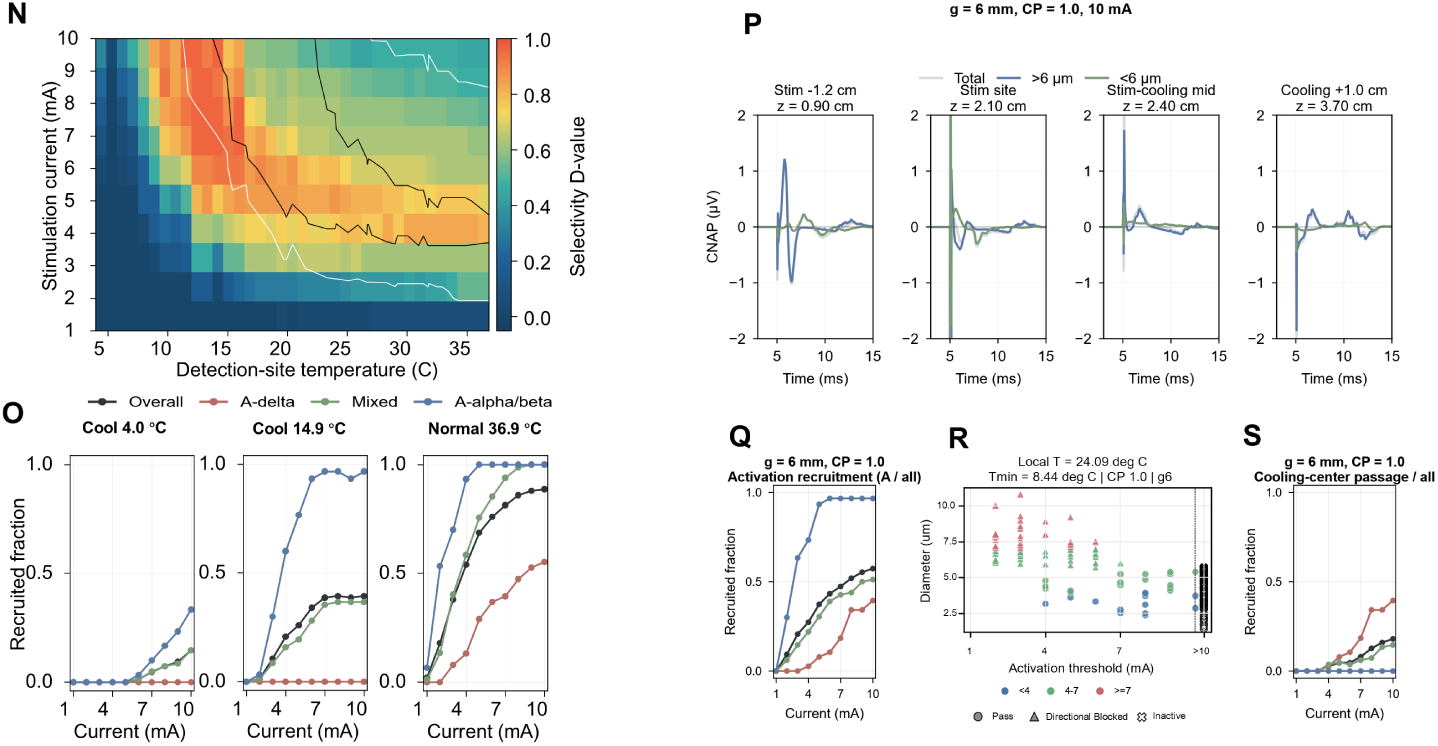
Local cooling reversibly reshapes bidirectional CNAP output and defines a directional, fiber-selective operating window. (A) Intraoperative photograph of the NeuroSwitch cuff implanted on the rat sciatic nerve. (B) Implantation schematic. (C) Experimental configuration showing local cooling within the cuff and upstream and downstream recordings used to distinguish afferent propagation toward the spinal cord from efferent propagation toward muscle. (D) Representative afferent and efferent CNAP waveforms at successive temperature plateaus during cooling and rewarming. (E) Time course of normalized quick-component amplitude and nerve temperature; the dashed line marks the end of cooling. CNAPs were evoked by cathodic-first, charge-balanced biphasic constant-current pulses (400 μA, 50 μs per phase). (F and G) Temperature dependence of normalized amplitude and peak latency for the quick, short-latency component enriched in Aα/Aβ activity in one rat. Points show the means of three repeated cycles, and error bars indicate SD. (H and I) Group-level quick-component amplitude and latency during cooling and rewarming (n = 3 rats); error bars indicate SD. (J and K) Temperature dependence of normalized amplitude and peak latency for the slow, longer-latency component enriched in Aδ activity in one rat. Points show the means of three repeated cycles, and error bars indicate SD. (L and M) Group-level slow-component amplitude and latency during cooling and rewarming (n = 3 rats); error bars indicate SD. At each temperature in (H), (I), (L), and (M), the value within the same recording direction and CNAP component was compared with its lowest-temperature value using a two-sided paired t test. Statistical annotations: ns, not significant; *P < 0.05; **P < 0.01; ***P < 0.001. (N) Selectivity D-value across stimulation current and detection-site temperature. (O) Recruited fractions of the overall population and the Aδ, mixed-diameter, and Aα/Aβ classes at representative normal and cooled temperatures. (P) Position-resolved simulated CNAPs along the stimulation-to-cooling axis, decomposed into total, >6-μm, and <6-μm contributions (10 mA; block–drive spacing, 6 mm; cooling-profile parameter, 1.0). (Q) Activation recruitment as a function of current under the same geometry. (R) Fiber diameter versus activation threshold, with outcomes classified as pass, directionally blocked, or inactive. (S) Fraction of fibers passing the cooling center as a function of stimulation current. Together, these simulations link the in vivo temperature-dependent CNAP changes in (A) to (M) to a propagation-aware control window.

Cooling reduced CNAP amplitudes in both directions, and the responses recovered after rewarming, demonstrating reversibility (Fig. 5D and E). The quick component, however, showed a clear directional difference. As temperature decreased, the afferent N1–P1 component was partially preserved, whereas the efferent N1–P1 component was strongly attenuated and nearly abolished; P1 latency also increased (Fig. 5F to I). Thus, some large-diameter fibers remained recruitable at the drive site, whereas propagation in the efferent direction was delayed or blocked by the cooling core, producing a directional bias. Because the efferent large-diameter response predominantly reflects the muscle-directed Aα motor pathway, its near-complete suppression argues against a substantial propagated Aα contribution to the retained afferent response and supports its interpretation as a directional, Aβ-biased afferent output.

In contrast, the Aδ-enriched slow component decreased substantially on both the afferent and efferent sides and showed increased P2 latency, without the directional difference observed for the quick component (Fig. 5J to M). This pattern indicates that attenuation of the slow component occurred primarily before the propagation pathways diverged and is consistent with suppression of small-diameter fiber recruitment by residual cooling near the stimulation site. The in vivo results therefore distinguish two forms of selectivity within the block–drive architecture: directional bias arises primarily from blocking nontarget propagation, whereas fiber bias arises primarily from recruitment filtering. Together, these effects convert the bidirectional mixed output of electrical stimulation into a directional, Aβ-biased afferent output that preferentially preserves the quick Aα/Aβ-enriched component while suppressing the slow Aδ-enriched component.

The digital twin further examined this mechanism and defined the corresponding operating window. The selectivity D value across the current–temperature plane identified a region in which Aα/Aβ recruitment was preferentially retained while Aδ recruitment was suppressed (Fig. 5N and O). When CNAPs were reconstructed along the axis from the drive site to the cooling center, fibers >6 μm and <6 μm showed distinct patterns of propagation attenuation (Fig. 5P; fig. S19). By integrating fiber activation, the relationship between fiber diameter and activation threshold, and propagation states across the cooling core, the model further distinguished fibers that were activated and passed, directionally blocked, or not activated (Fig. 5Q to S; fig. S19). These simulations linked the experimentally observed recruitment filtering and block of nontarget propagation to specific fiber-level states.

These findings define the principal operating constraints for cooling-gated block–drive. The drive current must reliably recruit the target Aβ-enriched component, while the temperature at the stimulation site must suppress Aδ recruitment without excessively suppressing the target Aβ-enriched component. At the same time, the cooling core along the nontarget propagation pathway must reach the conduction-block threshold, and the block–drive spacing must keep this colder region away from the target propagation pathway while maintaining an appropriate residual temperature gradient at the drive site. Together, these constraints connect the current–temperature operating window with the in vivo CNAP measurements, propagation-aware simulations, and block–drive spacing tests (Fig. 2N; Fig. 5F to S; fig. S2J to N).

Finally, hematoxylin and eosin (H&E) and Luxol fast blue staining revealed no apparent abnormalities in nerve architecture, myelinated-area fraction, or immune-cell density after acute cooling–rewarming and KHFS cycles or 60 days after implantation of the device head (fig. S18A to D; see Materials and Methods). These findings provide preliminary histological support for the tissue compatibility of the control strategy and device materials.

## Discussion

Together, these results show that temperature provides a simple physiological axis for reshaping how mixed peripheral nerves respond to electrical stimulation. Cooling shifts local neural state, and different fiber populations respond differently to this shift, allowing the same electrical input to produce distinct patterns of activation, propagation, and block. In this study, this principle enabled onset-suppressed KHFS block and a directional, Aβ-biased afferent output. Selectivity therefore emerges from the interaction between electrical input and temperature-dependent neural state.

These implementations are examples rather than endpoints of this control principle. The cooling–electrical neural digital twin links temperature-dependent neural responses to nerve geometry and field distributions, providing a basis for extending the same principle to other selective-control objectives and peripheral nerves. Thus, a simple physiological variable can support diverse neural control states beyond those accessible through electrical parameter tuning alone.

## Acknowledgments

Unless otherwise noted, schematic illustrations were drawn by the authors using Procreate. Illustration elements in Fig. 1B–E were adapted from Servier Medical Art (https://smart.servier.com/), and the rat illustration in Fig. 5B was adapted from Antonis Asiminas, SciDraw (doi:10.5281/zenodo.3926277); both are licensed under CC BY 4.0 (https://creativecommons.org/licenses/by/4.0/). During the preparation of this manuscript, the authors used ChatGPT (OpenAI) to edit the English text and improve readability. These tools did not generate scientific content, figures, references, or interpretations.

## Author contributions

S.Y. conceived the study and completed all major stages of the work, including conceptual development; device and experimental design; device fabrication and benchtop characterization; animal experiments; electrophysiological testing; modeling and simulation; data processing; figure preparation; and manuscript drafting. X.Y. primarily participated in animal experiments, neural electrophysiological data processing, and histological analyses. Shihao Yang, Qianqian Wang, Y.L., and G.-Q.B. contributed to scientific discussion and manuscript revision. Z.D. supervised the study, guided the overall scientific direction, and revised the manuscript. All authors reviewed and approved the final manuscript.

## Data, code, and materials availability

Data, analysis code, and simulation resources are being curated in a private GitHub repository at https://github.com/yangshu717/multimodal_NeuroSwitch. Access will be provided to editors and reviewers upon request. The repository will be made public upon publication and archived as a versioned release with a permanent DOI. NeuroSwitch devices and fabrication files are available from the lead contact upon reasonable request.

## References

1. S. S. Casagrande, A. Z. Beccera, K. F. Rust, C. C. Cowie, Opioid prescription and diabetes among Medicare beneficiaries. Diabetes Res Clin Pract 196, 110240 (2023).

2. T. H. Nost et al., Prevalence of substance use disorder diagnoses in patients with chronic pain receiving reimbursed opioids: An epidemiological study of four Norwegian health registries. Scand J Pain 24, 20240059 (2024).

3. N. D. Volkow, D. L. Longo, A. T. McLellan, Opioid Abuse in Chronic Pain — Misconceptions and Mitigation Strategies. New England Journal of Medicine 374, 1253–1263 (2016).

4. Y. J. Kim et al., Wireless and bioresorbable triboelectric nerve block system for postoperative pain control. Nature Biomedical Engineering. 2026 (10.1038/s41551-025-01579-2).

5. G. Lee et al., A bioresorbable peripheral nerve stimulator for electronic pain block. Sci Adv 8, eabp9169 (2022).

6. J. T. Reeder et al., Soft, bioresorbable coolers for reversible conduction block of peripheral nerves. Science 377, 109–115 (2022).

7. C.-E. Wong et al., Sciatic nerve stimulation alleviates acute neuropathic pain via modulation of neuroinflammation and descending pain inhibition in a rodent model. Journal of Neuroinflammation 19, 153 (2022).

8. C. E. Wong et al., Sciatic nerve stimulation alleviates neuropathic pain and associated neuroinflammation in the dorsal root ganglia in a rodent model. J Transl Med 22, 770 (2024).

9. K. L. Kilgore, N. Bhadra, Reversible nerve conduction block using kilohertz frequency alternating current. Neuromodulation 17, 242–254; discussion 254–255 (2014).

10. S. Liu et al., Neural basis of transcutaneous electrical nerve stimulation for neuropathic pain relief. Neuron 113, 3616–3631.e6 (2025).

11. D. D. Price, Dorsal horn neuronal responses and quantitative sensory testing help explain normal and abnormal pain. Pain 154, 1161–1162 (2013).

12. X. Navarro et al., A critical review of interfaces with the peripheral nervous system for the control of neuroprostheses and hybrid bionic systems. Journal of the Peripheral Nervous System 10, 229–258 (2005).

13. N. Jayaprakash et al., Organ- and function-specific anatomical organization of vagal fibers supports fascicular vagus nerve stimulation. Brain Stimul 16, 484–506 (2023).

14. U. Latif et al., Consensus Guidelines for the Use of Peripheral Nerve Stimulation in the Treatment of Chronic Pain and Neurological Diseases: A Neuron Project from the American Society of Pain and Neuroscience. J Pain Res 18, 5949–5990 (2025).

15. D. M. Ackermann, Jr., N. Bhadra, E. L. Foldes, X. F. Wang, K. L. Kilgore, Effect of nerve cuff electrode geometry on onset response firing in high-frequency nerve conduction block. IEEE Trans Neural Syst Rehabil Eng 18, 658–665 (2010).

16. D. M. Ackermann, N. Bhadra, M. Gerges, P. J. Thomas, Dynamics and sensitivity analysis of high-frequency conduction block. J Neural Eng 8, 065007 (2011).

17. C. A. Miller, P. J. Abbas, K. V. Nourski, N. Hu, B. K. Robinson, Electrode configuration influences action potential initiation site and ensemble stochastic response properties. Hearing Research 175, 200–214 (2003).

18. K. P. Cheng et al., Application of kilohertz-frequency block to mitigate off-target motor effects of vagus nerve stimulation in swine. Nat Commun 17, 1066 (2025).

19. C. Tai, J. Wang, M. B. Chancellor, J. R. Roppolo, W. C. de Groat, Influence of temperature on pudendal nerve block induced by high frequency biphasic electrical current. J Urol 180, 1173–8 (2008).

20. U. Ahmed et al., Anodal block permits directional vagus nerve stimulation. Sci Rep 10, 9221 (2020).

21. C. van den Honert, J. T. Mortimer, Generation of unidirectionally propagated action potentials in a peripheral nerve by brief stimuli. Science 206, 1311–2 (1979).

22. F. Ciotti et al., Towards enhanced functionality of vagus neuroprostheses through in silico optimized stimulation. Nat Commun 15, 6119 (2024).

23. D. M. Ackermann, Jr., E. L. Foldes, N. Bhadra, K. L. Kilgore, Effect of bipolar cuff electrode design on block thresholds in high-frequency electrical neural conduction block. IEEE Trans Neural Syst Rehabil Eng 17, 469–77 (2009).

24. Y. A. Patel, R. J. Butera, Differential fiber-specific block of nerve conduction in mammalian peripheral nerves using kilohertz electrical stimulation. J Neurophysiol 113, 3923–9 (2015).

25. N. Bhadra, K. L. Kilgore, High-frequency electrical conduction block of mammalian peripheral motor nerve. Muscle Nerve 32, 782–90 (2005).

26. M. Franke et al., Combined KHFAC + DC nerve block without onset or reduced nerve conductivity after block. J Neural Eng 11, 056012 (2014).

27. F. Lembeck, P. Holzer, Substance P as neurogenic mediator of antidromic vasodilation and neurogenic plasma extravasation. Naunyn Schmiedebergs Arch Pharmacol 310, 175–83 (1979).

28. I. M. Chiu, C. A. von Hehn, C. J. Woolf, Neurogenic inflammation and the peripheral nervous system in host defense and immunopathology. Nature Neuroscience 15, 1063–1067 (2012).

29. S. D. Brain, T. J. Williams, J. R. Tippins, H. R. Morris, I. MacIntyre, Calcitonin gene-related peptide is a potent vasodilator. Nature 313, 54–6 (1985).

30. W. Janig, S. J. Lisney, Small diameter myelinated afferents produce vasodilatation but not plasma extravasation in rat skin. J Physiol 415, 477–86 (1989).

31. R. A. Rietmeijer, B. Sorum, B. Li, S. G. Brohawn, Physical basis for distinct basal and mechanically gated activity of the human K(+) channel TRAAK. Neuron 109, 2902–2913.e4 (2021).

32. S. G. Brohawn et al., The mechanosensitive ion channel TRAAK is localized to the mammalian node of Ranvier. Elife 8, e50403 (2019).

33. H. Kanda, S. Tonomura, J. G. Gu, Effects of Cooling Temperatures via Thermal K2P Channels on Regeneration of High-Frequency Action Potentials at Nodes of Ranvier of Rat Abeta-Afferent Nerves. eNeuro 8, ENEURO.0308–21.2021 (2021).

34. K. Zimmermann et al., Sensory neuron sodium channel Nav1.8 is essential for pain at low temperatures. Nature 447, 855–8 (2007).

35. J. R. Schwarz, The effect of temperature on Na currents in rat myelinated nerve fibres. Pflugers Arch 406, 397–404 (1986).

36. S. G. Lomber, B. R. Payne, J. A. Horel, The cryoloop: an adaptable reversible cooling deactivation method for behavioral or electrophysiological assessment of neural function. J Neurosci Methods 86, 179–94 (1999).

37. P. C. Petersen, G. Buzsaki, Cooling of Medial Septum Reveals Theta Phase Lag Coordination of Hippocampal Cell Assemblies. Neuron 107, 731–744.e3 (2020).

38. P. Borgdorff, P. G. Versteeg, An implantable nerve cooler for the exercising dog. Eur J Appl Physiol Occup Physiol 53, 175–9 (1984).

39. T. Morgan et al., Thermal block of mammalian unmyelinated C fibers by local cooling to 15-25 degrees C after a brief heating at 45 degrees C. J Neurophysiol 123, 2173–2179 (2020).

40. D. M. Ackermann, E. L. Foldes, N. Bhadra, K. L. Kilgore, Nerve conduction block using combined thermoelectric cooling and high frequency electrical stimulation. J Neurosci Methods 193, 72–6 (2010).

41. C. C. H. Cohen et al., Saltatory Conduction along Myelinated Axons Involves a Periaxonal Nanocircuit. Cell 180, 311–322.e15 (2020).

42. C. C. McIntyre, A. G. Richardson, W. M. Grill, Modeling the excitability of mammalian nerve fibers: influence of afterpotentials on the recovery cycle. J Neurophysiol 87, 995–1006 (2002).

43. S. F. Lempka, C. C. McIntyre, K. L. Kilgore, A. G. Machado, Computational analysis of kilohertz frequency spinal cord stimulation for chronic pain management. Anesthesiology 122, 1362–76 (2015).

44. E. D. Musselman, N. A. Pelot, W. M. Grill, Validated computational models predict vagus nerve stimulation thresholds in preclinical animals and humans. J Neural Eng 20, 036032 (2023).

45. M. A. Hussain, W. M. Grill, N. A. Pelot, Highly efficient modeling and optimization of neural fiber responses to electrical stimulation. Nat Commun 15, 7597 (2024).

46. D. P. Marshall, E. S. Farah, E. D. Musselman, N. A. Pelot, W. M. Grill, PyFibers: An open-source NEURON-Python package to simulate responses of model nerve fibers to electrical stimulation. PLoS Comput Biol 21, e1013764 (2025).

47. C. Boehler, S. Carli, L. Fadiga, T. Stieglitz, M. Asplund, Tutorial: guidelines for standardized performance tests for electrodes intended for neural interfaces and bioelectronics. Nat Protoc 15, 3557–3578 (2020).

48. K. G. Beam, P. L. Donaldson, A quantitative study of potassium channel kinetics in rat skeletal muscle from 1 to 37 degrees C. J Gen Physiol 81, 485–512 (1983).

49. H. Kanda et al., TREK-1 and TRAAK Are Principal K(+) Channels at the Nodes of Ranvier for Rapid Action Potential Conduction on Mammalian Myelinated Afferent Nerves. Neuron 104, 960–971.e7 (2019).

50. M. Schewe et al., A Non-canonical Voltage-Sensing Mechanism Controls Gating in K2P K(+) Channels. Cell 164, 937–49 (2016).

51. M. Volgushev, T. R. Vidyasagar, M. Chistiakova, T. Yousef, U. T. Eysel, Membrane properties and spike generation in rat visual cortical cells during reversible cooling. J Physiol 522 **Pt** **1**, 59–76 (2000).

52. B. Bromm, Spike frequency of the nodal membrane generated by high-frequency alternating current. Pflugers Arch 353, 1–19 (1975).

53. E. D. Musselman, J. E. Cariello, W. M. Grill, N. A. Pelot, ASCENT (Automated Simulations to Characterize Electrical Nerve Thresholds): A pipeline for sample-specific computational modeling of electrical stimulation of peripheral nerves. PLOS Computational Biology 17, e1009285 (2021).

54. M. L. Hines, N. T. Carnevale, The NEURON simulation environment. Neural Comput 9, 1179–209 (1997).

55. C. A. Bossetti, M. J. Birdno, W. M. Grill, Analysis of the quasi-static approximation for calculating potentials generated by neural stimulation. J Neural Eng 5, 44–53 (2008).

56. A. L. Hodgkin, A. F. Huxley, A quantitative description of membrane current and its application to conduction and excitation in nerve. The Journal of Physiology 117, 500–544 (1952).

57. E. Peña, N. A. Pelot, W. M. Grill, Computational models of compound nerve action potentials: Efficient filter-based methods to quantify effects of tissue conductivities, conduction distance, and nerve fiber parameters. PLOS Computational Biology 20, e1011833 (2024).

58. R. V. Shannon, A model of safe levels for electrical stimulation. IEEE Transactions on Biomedical Engineering 39, 424–426 (1992).

59. S. F. Cogan, Neural stimulation and recording electrodes. Annu Rev Biomed Eng 10, 275–309 (2008).

60. G. Yi, W. M. Grill, Kilohertz waveforms optimized to produce closed-state Na+ channel inactivation eliminate onset response in nerve conduction block. PLOS Computational Biology 16, e1007766 (2020).

61. E. Peña, N. A. Pelot, W. M. Grill, Non-monotonic kilohertz frequency neural block thresholds arise from amplitude- and frequency-dependent charge imbalance. Scientific Reports 11, 5077 (2021).

62. E. Peña, N. A. Pelot, W. M. Grill, Spatiotemporal parameters for energy efficient kilohertz-frequency nerve block with low onset response. Journal of NeuroEngineering and Rehabilitation 20, 72 (2023).

63. M. C. Kiernan, Effects of temperature on the excitability properties of human motor axons. Brain 124, 816–825 (2001).

64. L. Rueda-Ruzafa, S. Herrera-Pérez, A. Campos-Ríos, J. A. Lamas, Are TREK Channels Temperature Sensors? Frontiers in Cellular Neuroscience 15, 744702 (2021).

65. F. Maingret, M. Fosset, F. Lesage, M. Lazdunski, E. Honoré, TRAAK Is a Mammalian Neuronal Mechano-gated K+ Channel. Journal of Biological Chemistry 274, 1381–1387 (1999).

66. F. Maingret et al., TREK-1 is a heat-activated background K+ channel. The EMBO Journal 19, 2483–2491 (2000).

67. J. Noël et al., The mechano-activated K+ channels TRAAK and TREK-1 control both warm and cold perception. The EMBO Journal 28, 1308–1318 (2009).

68. S. G. Brohawn, E. B. Campbell, R. MacKinnon, Physical mechanism for gating and mechanosensitivity of the human TRAAK K+ channel. Nature 516, 126–130 (2014).

69. C. Baumgartner et al. (IT’IS Foundation, 2025).

70. N. A. Pelot, C. E. Behrend, W. M. Grill, On the parameters used in finite element modeling of compound peripheral nerves. J Neural Eng 16, 016007 (2019).

71. S. Gabriel, R. W. Lau, C. Gabriel, The dielectric properties of biological tissues: III. Parametric models for the dielectric spectrum of tissues. Phys Med Biol 41, 2271–93 (1996).

72. H. Ye, J. Ng, Shielding effects of myelin sheath on axolemma depolarization under transverse electric field stimulation. PeerJ 6, e6020 (2018).

73. C. Johnson, W. R. Holmes, A. Brown, P. Jung, Minimizing the caliber of myelinated axons by means of nodal constrictions. J Neurophysiol 114, 1874–84 (2015).

74. C. H. Berthold, M. Rydmark, Electrophysiology and morphology of myelinated nerve fibers. VI. Anatomy of the paranode-node-paranode region in the cat. Experientia 39, 964–76 (1983).

75. S. Y. Chiu, J. M. Ritchie, R. B. Rogart, D. Stagg, A quantitative description of membrane currents in rabbit myelinated nerve. J Physiol 292, 149–66 (1979).

76. H. H. Pennes, Analysis of tissue and arterial blood temperatures in the resting human forearm. J Appl Physiol 1, 93–122 (1948).

