## Supplementary Materials for "Precision cooling assists electrical stimulation: Breaking selectivity barriers in nerve control"

**Contents**

- Materials and Methods
- Supplementary Text
- Figs. S1 to S19
- Tables S1 to S11
- Data and code availability

**Equation numbering.** Because no equations are numbered in the main text, equations in this Supplementary Materials file are numbered consecutively as Eq. 1, Eq. 2, and so on.

### Materials and Methods

Study design and reporting overview

This study combines device engineering, benchtop temperature-field and electrochemical characterization, acute and chronic in vivo experiments, and a digital twin coupling cooling, electrical stimulation, and neural dynamics. The main experimental logic is to test whether local cooling adds an operational control dimension to peripheral nerve stimulation. Main-text figures report the conceptual framework, fiber-control boundary, KHFS onset suppression, CNAP decomposition, and in vivo block–drive output. Supplementary figures and tables provide the detailed engineering, temperature-field, electrochemical, histological, and modeling support (Figs. 1 to 5; figs. S1 to S19; Tables S1 to S10).

Experimental, device, thermal, electrochemical, in vivo, and histology methods

Animals and ethics

All in vivo experiments were performed in adult male Sprague-Dawley rats weighing 250–300 g. Animals were housed under standard barrier conditions with ad libitum access to food and water. Surgery and in vivo electrophysiology were conducted under inhaled isoflurane anesthesia, and body temperature was maintained throughout the procedure. All procedures were approved by the Animal Care and Use Committee of the Shenzhen Institutes of Advanced Technology, Chinese Academy of Sciences (protocol IACUC-2023-009) and were performed in accordance with AAALAC International standards. Exact $n$ values, the biological unit represented by each $n$, and the corresponding statistical comparisons are provided in the figure legends, main text, and quantification section (fig. S18A to D; see Methods and figure legends).

Device design and fabrication

To characterize the physical performance of the cooling and stimulation modules before finalizing the implantable device, we first fabricated a test device containing a central block region and symmetric drive regions. The test structure used the same thermally conductive block-region architecture, electrode material, electrode line width, and electrode spacing as the final NeuroSwitch cuff, allowing the thermal-management and electrochemical results obtained from the test platform to constrain the final device directly. The final NeuroSwitch cuff had a flattened geometry with a 5-mm chamber width, a 600-μm internal height, and a 9-mm center-to-center spacing between the block and drive regions.

The electrode layer was fabricated using a high-precision laser-assisted transfer and patterning workflow (fig. S1E and F). Metal interconnects and insulating layers were built sequentially on an elastic substrate, after which the block- and drive-region traces and stimulation sites were patterned using an LPKF S4 laser-processing platform. In the final device, Pt sites were optimized for the rat sciatic nerve, with a 300-μm line width and a 1-mm spacing between adjacent sites. The block region integrated a ring-like tripolar stimulation configuration beneath the nerve, a reserved cooling region above the nerve, and a patch thermistor at the center of the block zone for real-time temperature readout.

A microfluidic mold was produced by high-resolution 3D printing, and PDMS replica molding was used to create the microchannel body. After surface activation, the channel body was bonded to a thin sealing membrane to form a closed fluidic conduit. The final channel cross-section was 300 μm × 300 μm (width × height), and the channel was aligned coaxially above the block-region nerve interface to create a stable local cooling window. Flexible tubing connected the distal end of the channel to the external cooling circuit.

The microfluidic and electrode layers were plasma-activated, aligned, and bonded so that the cooling channel was positioned directly above the block region. To reduce local thermal resistance while maintaining mechanical compliance, a 50 wt.% diamond-filled PDMS layer (mean particle diameter, 2.5 μm) was introduced between the microchannel and the nerve-facing electrode interface. The device head was then shaped into a flat cuff form that conformed to the surface of the rat sciatic nerve (Fig. 1I to K; fig. S1D and H to J; Table S2).

Cooling system and thermal characterization

For the bench cooling experiments in the tissue-mimicking phantom, ethanol was driven through the microfluidic channel by an external pump after precooling in a dry-ice trap. The trap consisted of coiled metal tubing immersed in the cooling medium, allowing the ethanol to remain within a stable low-temperature window before entering the device (fig. S5A to G; Table S3). Under the operating conditions used here, the inlet temperature was approximately −10 °C to −16 °Cover a measured flow-rate range of 0–5.5 mL $min^{-1}$. Inlet and outlet coolant temperatures were monitored by thermocouples, and block-zone temperature was read simultaneously from the integrated thermistor using a custom Arduino Uno-based readout system.

To evaluate localized cooling performance, the device head was embedded in uncured 1 wt.% agar and allowed to solidify, thereby forming a tissue-mimicking phantom nerve and surrounding soft-tissue environment. The phantom was held in a 37 °C water bath. Six temperature-measurement locations were defined as in figs. S5A and S6A: point 1 at the nerve center beneath the microchannel; point 2 at a lateral position 4 mm from the channel center; point 3 at the embedded block-zone thermistor; points 4 and 5 at the coolant outlet and inlet, respectively; and point 6 in the surrounding agar. With the dry-ice trap held at about −35 °C, flow was varied stepwise while all temperature channels were recorded simultaneously.

To visualize the spatial distribution of cooling, the assembled implant was cooled in air using the same external cooling system and imaged with a Hikvision H11 Pro infrared camera fitted with a macro lens. To measure thermal properties as a function of material composition, cylindrical samples (2-cm diameter, 1-cm thickness; two replicates per condition) were prepared and tested using a TPS 2500S thermal-constants analyzer according to the standard instrument protocol. Thermal conductivity, specific heat, and thermal diffusivity were compared across diamond-loading conditions.

Cooling-test data were analyzed offline in Python. The thermistor trace from the block zone was converted to elapsed time and resampled onto a uniform time grid using the median sampling interval; when the estimated interval was close to 2 s, a 2-s resampling step was used. Steady-state plateaus were detected using a rolling local linear-regression slope and a rolling standard deviation computed over a 120-s odd-length window. Samples were classified as steady state only when both the absolute local slope and the rolling standard deviation were below data-driven thresholds, and only plateau segments lasting at least 80 s were retained. The representative steady-state temperature of each plateau was defined as the median value of the plateau core after trimming the first and last 10% of the segment, or at least 20 s from each end. The SD of this trimmed core window was used as the within-plateau variability. The point-1 versus point-3 traces were compared by normalized full cross-correlation after mean-centering to determine whether a measurable delay existed between the intraneural and embedded-sensor readouts. For the flow-temperature relation, plateau temperatures were paired with coolant flow rates and fitted with a constrained single-exponential model (fig. S5H; fig. S6B to E; Table S3).

Electrochemical characterization

Electrochemical measurements were performed in PBS (pH 7.4) with an Ag/AgCl reference electrode. For a single Pt site, cyclic voltammetry, electrochemical impedance spectroscopy, and voltage-transient measurements under biphasic pulsing were recorded. The exposed Pt area used for charge-injection-capacity estimation was 0.3 mm × 10 mm. Cyclic voltammetry was performed at 50 mV/s. Voltage transients were recorded under 500-μs-per-phase charge-balanced biphasic pulses, and the safe single-phase current and charge-injection capacity were estimated using a water-window constraint of −0.8 to +1.0 V. Pulse responses were recorded with a DScope U3P100 oscilloscope (fig. S7; Table S4).

Chronic implantation, histology, and image analysis

For 60-d chronic cuff implantation around the sciatic nerve, animals were placed on a thermostatically controlled heating pad after surgery until fully recovered. Animals were returned to their home cages only after the righting reflex and spontaneous activity had stabilized. During the first 72 h after surgery, meloxicam (1.5 mg/kg, subcutaneous, once daily) was administered for analgesia. During this acute recovery period, body weight, spontaneous activity, posture, grooming, food and water intake, and incision condition were assessed at least daily. Particular attention was paid to self-biting or self-injury of the operated hindlimb and to guarding or reduced limb use. Thereafter, animals were examined two to three times per week throughout the 60-d implantation period for wound healing, local inflammation, infection, gait abnormalities, and device-related complications such as exposed leads. Readily accessible food and hydrogel were provided as supportive care when needed.

Prespecified humane endpoints included severe infection, wound dehiscence, persistent self-injury, body-weight loss of at least 20% relative to the preoperative baseline, or persistent distress. Animals reaching an endpoint were euthanized promptly according to the approved protocol. All animal procedures, postoperative monitoring, and interventions were performed by trained personnel under protocol IACUC-2023-009 and in accordance with AAALAC International standards.

For histological assessment, sciatic-nerve samples collected after acute experiments and at longer implantation time points were fixed, paraffin embedded, and sectioned serially at 4 μm. Adjacent sections were stained with H&E to assess fascicular architecture and granulocytic infiltration or with Luxol fast blue (LFB) to assess myelin integrity. Sections were first examined on an Olympus FV3000 microscope and then scanned on a VS200-BU slide scanner using consistent bright-field settings across groups (fig. S18A to D; Table S1 for histology resources).

Granulocytes in H&E-stained sections were quantified using a semiautomated ImageJ workflow with manual correction. Three randomly selected, nonoverlapping fields (164.5 × 109.3 μm per field) were counted in each section plane and averaged. Three section planes were analyzed per nerve per rat, and their mean was treated as the final value for that nerve. When required, counts were normalized to field area and reported as cell density.

LFB bright-field images were processed using a local Python workflow. Images were converted to grayscale and enhanced using fixed parameters, after which axons and myelin were segmented with the AxonDeepSeg bright-field model model_seg_generalist_BF_light. A closed polygonal region of interest (ROI) was drawn manually on the enhanced image, and subsequent analysis was restricted to the corresponding binary ROI mask. Axon and myelin areas were defined by the numbers of nonzero pixels in their respective masks, and myelin-area fraction was calculated as myelin/(myelin + axon). Because pixel dimensions were not physically calibrated in this workflow, these measurements represent relative pixel-domain area fractions for comparisons among samples and groups rather than absolute morphometric areas. Multiple ROIs or section planes from the same animal were averaged within animal before statistical analysis.

In vivo electrophysiology and CNAP recording

For in vivo acute electrophysiology, animals were maintained on a thermostatically controlled heating pad under inhaled isoflurane anesthesia. The surgical site was shaved and disinfected, erythromycin ophthalmic ointment was applied to prevent corneal drying, and exposed tissues were kept moist with sterile saline throughout the procedure. Sterile sutures were used for device fixation or incision management as required by the experimental workflow. The rat sciatic nerve was surgically exposed and the NeuroSwitch cuff was placed around the nerve. Recording cuffs were positioned upstream and downstream to resolve afferent (toward the spinal cord) and efferent (toward the muscle) CNAP propagation. Charge-balanced cathodic-first biphasic constant-current pulses (400 μA, 50 μs per phase unless otherwise noted) were used for low-frequency activation. CNAPs were quantified by peak-to-peak amplitude and peak latency, with the fast component corresponding mainly to Aα/Aβ-enriched signaling and the slow component corresponding mainly to Aδ-enriched signaling.

To identify the minimum block–drive spacing compatible with directional control, a multi-spacing test cuff was implanted so that different block–drive separations could be evaluated within the same preparation. During low-frequency drive combined with tripolar KHFS block, CNAPs were recorded on both sides of the cuff to determine when the block field began to interfere directly with the activated region. This mapping was used to define a low-interference spacing window and guide the final 8- to 9-mm design choice.

To characterize cooling block in vivo, the cuff was precooled while afferent and efferent CNAPs were recorded continuously during cooling and rewarming. Temperature plateaus were used to quantify amplitude and latency changes in fast and slow CNAP components and to identify the temperature windows in which conduction was reduced, abolished, or recovered.

To quantify onset-related side effects during kilohertz block, tripolar 10-kHz stimulation was initiated intermittently at different nerve temperatures, and the resulting ankle-angle deflection was recorded as a measure of onset-induced muscle twitch. The peak angular deflection after KHFS onset was used as the onset metric for the analysis shown in Fig. 3A and B.

To test the cooling-to-KHFS handoff strategy, cooling was first used to establish block, after which kilohertz stimulation was initiated during rewarming to maintain suppression. In separate experiments, low-frequency stimulation was delivered at the drive site while the cooled block site acted as a directional valve. Afferent and efferent CNAPs were monitored continuously to determine whether propagation was preserved in the desired direction while suppressed in the undesired direction.

CNAP signals were processed offline using peak-to-peak amplitude and peak latency as the primary features (Figs. 4 and 5; fig. S16). Opposite stimulus polarities were compared to separate propagating CNAP components from stimulus artifacts. Fast and slow components were identified on the basis of latency and waveform morphology and tracked across temperature, block, and activation conditions.

**Device-level cooling-assisted electrical field and neural digital-twin methods**

The following sections provide the formula-based device-level and neural digital-twin methods (Figs. 2 to 4; figs. S3, S4, and S8 to S15; Tables S4 to S10). Overlapping descriptions from the original STAR-methods-derived support and the standardized modeling workflow were merged so that each governing equation, model assumption, and implementation step appears once in the main support body.

Digital twin and simulations

We constructed a coupled device-nerve finite-element model that integrated cooling-induced temperature fields, stimulation-induced extracellular potentials, and axonal membrane dynamics within a common computational framework. The model used the deformed nerve geometry and device layout described in the supplementary figures and incorporated material thermal and electrical properties from the parameter tables. The simulation framework was designed to define transferable relationships among block–drive spacing, cooling windows, stimulation conditions, and membrane-state transitions.

The thermal-fluid module solved incompressible laminar flow and conjugate heat transfer in the microchannel and surrounding solid/tissue domains. The electrical module solved the stimulus-induced extracellular field using a quasi-static potential formulation. The electrophysiology module was defined on region-specific axonal membrane boundaries and was driven by sampled local temperature and extracellular potential at each membrane compartment.

Thermal-fluid and heat-transfer equations

The thermal module used the following governing equations for microchannel flow, conjugate heat transfer, tissue bioheat, and heat-source terms (fig. S3C to H; figs. S5A and S6A; Tables S3 and S10). Definitions of the five principal field variables are summarized in Table S11; remaining symbols and material constants are defined locally and in the parameter tables.

Flow incompressibility constraint:

$$\nabla \cdot \mathbf{u} = 0,$$

(Eq. 1)

Navier-Stokes momentum balance:

$$\rho_f \left( \frac{\partial \mathbf{u}}{\partial t} + (\mathbf{u} \cdot \nabla)\mathbf{u} \right) = -\nabla p + \mu_f \nabla^2 \mathbf{u} + \mathbf{f},$$

(Eq. 2)

Fluid-domain advection-diffusion heat equation:

$$\rho_f C_{p,f} \left( \frac{\partial T}{\partial t} + \mathbf{u} \cdot \nabla T \right) = \nabla \cdot \left( k_f \nabla T \right) + Q_f,$$

(Eq. 3)

Temperature and heat-flux continuity at solid-fluid interfaces:

$$T_{solid} = T_{fluid}, \qquad \mathbf{n} \cdot k_{solid} \nabla T_{solid} = \mathbf{n} \cdot k_f \nabla T_{fluid}.$$

(Eq. 4)

Solid-domain transient heat equation:

$$\rho C_p \frac{\partial T}{\partial t} = \nabla \cdot (k \nabla T) + Q,$$

(Eq. 5)

Perfusion and metabolic heat-source terms:

$$Q = Q_{met} + Q_{perf}, \qquad Q_{perf} = \rho_b C_b \omega_b (T_b - T),$$

(Eq. 6)

Pennes-type tissue bioheat equation:

$$\rho C_p \frac{\partial T}{\partial t} = \nabla \cdot (k \nabla T) + Q_{met} + \rho_b C_b \omega_b (T_b - T).$$

(Eq. 7)

Joule-heating source term:

$$Q_J = \mathbf{J} \cdot \mathbf{E},$$

(Eq. 8)

Electrical-field and transmembrane-coupling equations

Electrical stimulation was represented with a quasi-static potential formulation (fig. S4C to E; Tables S4 and S5). The field solution was coupled to the membrane model through extracellular and intracellular normal-current boundary conditions so that capacitive and ionic membrane currents entered the field problem with consistent sign conventions.

Electric field from scalar potential:

$$\mathbf{E} = -\nabla V,$$

(Eq. 9)

Conductive and displacement current densities:

$$\mathbf{J} = \sigma \mathbf{E} + \frac{\partial \mathbf{D}}{\partial t}, \qquad \mathbf{D} = \varepsilon_0 \varepsilon_r \mathbf{E}.$$

(Eq. 10)

Quasi-static conduction-displacement potential equation:

$$\nabla \cdot (\sigma \nabla V) + \nabla \cdot \left( \varepsilon_0 \varepsilon_r \frac{\partial}{\partial t} \nabla V \right) = 0.$$

(Eq. 11)

Low-frequency terminal current:

$$I_{LF}(t) = I_0 \, s_{LF}(t),$$

(Eq. 12)

Kilohertz terminal currents for the bipolar pair:

$$I_{HF+}(t) = \frac{I_{pp}}{2} \, s_{HF}(t), \qquad I_{HF-}(t) = -\frac{I_{pp}}{2} \, s_{HF}(t),$$

(Eq. 13)

Dimensionless square-wave carrier:

$$s_{HF}(t) = sign\big(\sin(2\pi f_{HF} t)\big) \in \{-1, +1\}.$$

(Eq. 14)

Extracellular-side membrane current boundary condition:

$$\mathbf{n} \cdot \mathbf{J}_e = -i_m,$$

(Eq. 15)

Intracellular-side membrane current boundary condition:

$$\mathbf{n} \cdot \mathbf{J}_i = +i_m,$$

(Eq. 16)

Capacitive and ionic transmembrane current density:

$$i_m = i_C + \sum_k i_k\,, \qquad i_C = C_m \frac{\partial V_m}{\partial t},$$

(Eq. 17)

Effective transmembrane voltage used by the coupling model:

$$V_m = V_i \cdot TkVm - V_e.$$

(Eq. 18)

Compact ionic-current expression:

$$i_k = g_k^{(r)}\, F_k(\mathbf{s}_k)\, drv\big(V_m - E_{k,T}\big)\, A_k(T),$$

(Eq. 19)

Temperature scaling and GHK interpretation

In the mixed-fiber model, temperature entered the membrane model through three coordinated pathways (Fig. 2K and M; figs. S9 and S10; fig. S11A to H; fig. S13F to H; Tables S7 to S9): rescaling of effective transmembrane voltage, temperature-aware adjustment of sodium and potassium reversal potentials, and $Q_{10}$-type or analytic scaling of channel kinetics and selected amplitude terms. The GHK relationships below were used as an interpretation layer linking the empirically constrained cooling-associated depolarization to permeability-weighted ionic contributions.

The effective membrane voltage used by channel kinetics was defined in Eq. 18.

Temperature mapping factor for effective membrane voltage:

$$TkVm = 1 + \frac{(T - T_{ref}) \cdot 0.45\,\mathrm{mV\,K^{-1}}}{|V_{rest}|}.$$

(Eq. 20)

Temperature-scaled potassium reversal term:

$$TkEk = 1 - \frac{(T - T_{ref}) \cdot 0.31\,\mathrm{mV\,K^{-1}}}{E_K}, \qquad E_{K,T} = E_K \cdot TkEk.$$

(Eq. 21)

Temperature-scaled K2P leak-like reversal potential:

$$E_{L,T} = E_L \cdot TkEk.$$

(Eq. 22)

Temperature-scaled sodium reversal potential:

$$E_{Na,T} = E_{Na} \cdot TkENa, \qquad TkENa = 1 + \frac{(T - T_{ref}) \cdot 0.2\,\mathrm{mV\,K^{-1}}}{E_{Na}}.$$

(Eq. 23)

Nernst interpretation of temperature-dependent reversal potentials:

$$E_x(T) = \frac{RT}{z_x F} \ln\left(\frac{[x]_o}{[x]_i}\right),$$

(Eq. 24)

Generic $Q_{10}$ scaling for time constants:

$$\tau(T) = \frac{\tau(T_{ref})}{Q_{10}^{\frac{T-T_{ref}}{10\,K}}}.$$

(Eq. 25)

Numerically stable exponential operator:

$$\mathrm{Exp}_{safe}(x) = \begin{cases} 0, & x < -100, \\ e^x, & \textbf{otherwise}. \end{cases}$$

(Eq. 26)

Saturating driving-force operator:

$$\mathrm{drv}_{0.2\,\mathrm{mV}}(x) = \mathrm{sgn}(x) \cdot 0.2\,\mathrm{mV} \cdot \left(1 - e^{-|x|/(0.2\,\mathrm{mV})}\right).$$

(Eq. 27)

Empirical cooling-associated membrane-potential slope:

$$\frac{dV_m}{dT} \approx -0.45\,\mathrm{mV\,K^{-1}}.$$

(Eq. 28)

Temperature-dependent permeability definition:

$$P_i(T) = P_i(T_{ref})\,S_i(T), \qquad i \in \{K, Na\},$$

(Eq. 29)

Eyring-like permeability scaling:

$$S_i(T) = \frac{T}{T_{ref}} \exp\left[-\frac{\Delta H_i^{\ddagger}}{R}\left(\frac{1}{T} - \frac{1}{T_{ref}}\right)\right], \qquad i \in \{K, Na\}.$$

(Eq. 30)

Reference and temperature-dependent Na/K permeability ratio:

$$r_{ref} = \frac{P_{Na}(T_{ref})}{P_K(T_{ref})}, \qquad r(T) = \frac{P_{Na}(T)}{P_K(T)} = r_{ref}\,\frac{S_{Na}(T)}{S_K(T)}.$$

(Eq. 31)

K/Na Goldman-Hodgkin-Katz membrane potential:

$$V_{GHK}(T) = \frac{RT}{F} \ln\left(\frac{[K]_o + r(T)[Na]_o}{[K]_i + r(T)[Na]_i}\right).$$

(Eq. 32)

Extracellular GHK permeability weights:

$$w_K^{(o)}(T) = \frac{[K]_o}{[K]_o + r(T)[Na]_o}, \qquad w_{Na}^{(o)}(T) = 1 - w_K^{(o)}(T),$$

(Eq. 33)

Intracellular GHK permeability weights:

$$w_K^{(i)}(T) = \frac{[K]_i}{[K]_i + r(T)[Na]_i}, \qquad w_{Na}^{(i)}(T) = 1 - w_K^{(i)}(T).$$

(Eq. 34)

Reference permeability ratio in the K-dominant limit:

$$r_{ref} = \frac{1 - w_K^{(o)}(T_{ref})}{w_K^{(o)}(T_{ref})} \cdot \frac{[K]_o}{[Na]_o}.$$

(Eq. 35)

Logarithmic temperature derivative of permeability:

$$\eta_i(T) = \frac{1}{P_i(T)}\frac{dP_i(T)}{dT} = \frac{d}{dT}\ln P_i(T), \qquad i \in \{K, Na\}.$$

(Eq. 36)

Analytic temperature slope of the GHK potential:

$$\frac{dV_{GHK}}{dT} = \frac{V_{GHK}(T)}{T} + \frac{RT}{F}\Big(w_K^{(o)}(T)\eta_K(T) + w_{Na}^{(o)}(T)\eta_{Na}(T) - w_K^{(i)}(T)\eta_K(T) - w_{Na}^{(i)}(T)\eta_{Na}(T)\Big).$$

(Eq. 37)

Ion-channel current and gating equations

The conductance-based membrane model included fast transient, slow persistent, and temperature-responsive $Na_V1.8$-like sodium components, delayed-rectifier potassium components, and a K2P-dominant nodal stabilization module representing TRAAK- and TREK-1-like behavior (figs. S9 and S10; Tables S7 to S9). Region-dependent conductance densities and kinetic coefficients are defined in the parameter tables.

Total sodium current:

When the temperature-sensitive upgraded $Na_V1.8$ model is incorporated into the active node, the total sodium current is:

$$i_{Na} = i_{Na16} + i_{Na11} + i_{Na18}.$$

(Eq. 38)

Temperature-shifted sodium activation voltage:

$$V_{m,Na_m} = V_m - \min(0,\ 0.000353\,\mathrm{V\,K^{-1}} \cdot (T - 310.15\,\mathrm{K})),$$

(Eq. 39)

Temperature-shifted sodium inactivation voltage:

$$V_{m,Na_h} = V_m - 0.735\,\mathrm{mV\,K^{-1}} \cdot (T - 310.15\,\mathrm{K}).$$

(Eq. 40)

$Q_{10}$ accelerations for sodium kinetics:

$$q_{10,1}(T) = 2.2^{(T-309.15\,\mathrm{K})/(10\,\mathrm{K})}, \qquad q_{10,2}(T) = 2.9^{(T-309.15\,\mathrm{K})/(10\,\mathrm{K})},$$

(Eq. 41)

Temperature-dependent sodium amplitude factor:

$$peak_{Na}(T) = \begin{cases} \dfrac{1.12^{(T-283.15)/10}}{1.12^{(310.15-293.15)/10}}, & T < 283.15, \\ \dfrac{1.12^{(T-283.15)/10}}{1.12^{(310.15-293.15)/10}} \cdot \dfrac{2.0^{(T-283.15)/10}}{1.12^{(T-283.15)/10}}, & T \geq 283.15. \end{cases}$$

(Eq. 42)

In Eqs. 43 and 49, the unqualified symbols gNa16 and gNa11 denote the maximum conductances used by the applicable implementation layer rather than one universal parameter pair. The EFM neuron model implementation uses the values in Table S8, whereas the compartmental active-node implementation uses the pre- or post-low-temperature-response-calibration values in Table S9.

Fast transient sodium current:

$$i_{Na16} = gNa16 \cdot m^3 \cdot h \cdot drv(V_m - E_{Na,T}) \cdot peak_{Na}(T).$$

(Eq. 43)

Activation-gate rate definitions:

$$\alpha_m = vtrap6(V_{m,Na_m}), \quad \beta_m = vtrap7(V_{m,Na_m}),$$

(Eq. 44)

Inactivation-gate rate definitions:

$$\alpha_h = vtrap8(V_m), \quad \beta_h = vtrap9(V_{m,Na_h}).$$

(Eq. 45)

Sodium-gate steady states and time constants:

$$m_\infty = \frac{\alpha_m}{\alpha_m + \beta_m}, \quad \tau_m = \frac{1}{\alpha_m + \beta_m}, \qquad h_\infty = \frac{\alpha_h}{\alpha_h + \beta_h}, \quad \tau_h = \frac{1}{\alpha_h + \beta_h}.$$

(Eq. 46)

Activation-gate ODE with temperature acceleration:

$$\frac{dm}{dt} = \frac{m_\infty(V_{m,Na_m}) - m}{\tau_m(V_m)/q_{10,1}(T)},$$

(Eq. 47)

Inactivation-gate ODE with temperature acceleration:

$$\frac{dh}{dt} = q_{10,2}(T)\big(\alpha_h(V_m)(1-h) - \beta_h(V_{m,Na_h})h\big).$$

(Eq. 48)

Slow/persistent sodium current:

$$i_{Na11} = gNa11 \cdot mp^3 \cdot drv(V_m - E_{Na,T}) \cdot peak_{Na}(T).$$

(Eq. 49)

Persistent sodium gate rates:

$$\alpha_{mp} = vtrap1(V_{m,Na_m}), \quad \beta_{mp} = vtrap2(V_{m,Na_m}).$$

(Eq. 50)

Persistent sodium steady state and time constant:

$$mp_{\infty} = \frac{\alpha_{mp}}{\alpha_{mp} + \beta_{mp}}, \qquad \tau_{mp} = \frac{1}{\alpha_{mp} + \beta_{mp}}.$$

(Eq. 51)

Persistent sodium gate ODE:

$$\frac{dmp}{dt} = \frac{mp_{\infty}(V_{m,Na_m}) - mp}{\tau_{mp}(V_m)/q_{10,1}(T)}.$$

(Eq. 52)

Numerically stable vtrap1 function:

$$vtrap1(x) = \begin{cases} ampA \cdot ampC\left(1 - \frac{x + ampB}{2ampC}\right), & \left|\frac{x + ampB}{ampC}\right| < \epsilon, \\ ampA \cdot \frac{x + ampB}{1 - Exp\left(-\frac{x + ampB}{ampC}\right)}, & \textbf{otherwise}. \end{cases}$$

(Eq. 53)

Numerically stable vtrap2 function:

$$vtrap2(x) = \begin{cases} bmpA \cdot bmpC\left(1 - \frac{x + bmpB}{2bmpC}\right), & \left|\frac{x + bmpB}{bmpC}\right| < \epsilon, \\ bmpA \cdot \frac{x + bmpB}{1 - Exp\left(-\frac{x + bmpB}{bmpC}\right)}, & \textbf{otherwise}. \end{cases}$$

(Eq. 54)

Numerically stable vtrap6 function:

$$vtrap6(x) = \begin{cases} amA \cdot amC\left(1 - \frac{x + amB}{2amC}\right), & \left|\frac{x + amB}{amC}\right| < \epsilon, \\ amA \cdot \frac{x + amB}{1 - Exp\left(-\frac{x + amB}{amC}\right)}, & \textbf{otherwise}. \end{cases}$$

(Eq. 55)

Numerically stable vtrap7 function:

$$vtrap7(x) = \begin{cases} bmA \cdot bmC\left(1 - \frac{-x - bmB}{2bmC}\right), & \left|\frac{-x - bmB}{bmC}\right| < \epsilon, \\ bmA \cdot \frac{-x - bmB}{1 - Exp\left(-\frac{-x - bmB}{bmC}\right)}, & \textbf{otherwise}. \end{cases}$$

(Eq. 56)

Numerically stable vtrap8 function:

$$vtrap8(x) = \begin{cases} ahA \cdot ahC\left(1 - \frac{-x - ahB}{2ahC}\right), & \left|\frac{-x - ahB}{ahC}\right| < \epsilon, \\ ahA \cdot \frac{-x - ahB}{1 - Exp\left(-\frac{-x - ahB}{ahC}\right)}, & \textbf{otherwise}. \end{cases}$$

(Eq. 57)

Numerically stable vtrap9 function:

$$vtrap9(x) = \begin{cases} bhA \cdot bhC\left(1 - \frac{x + bhB}{2bhC}\right), & \left|\frac{x + bhB}{bhC}\right| < \epsilon, \\ bhA \cdot \frac{x + bhB}{1 - Exp\left(-\frac{x + bhB}{bhC}\right)}, & \textbf{otherwise}. \end{cases}$$

(Eq. 58)

Initial sodium-gate states:

$$m(0) = m_\infty(V_{rest} \cdot TkVm), \quad h(0) = h_\infty(V_{rest} \cdot TkVm), \quad mp(0) = mp_\infty(V_{rest} \cdot TkVm), \quad s(0) = s_\infty(V_{rest} \cdot TkVm), \quad u(0) = u_\infty(V_{rest} \cdot TkVm).$$

(Eq. 59)

Temperature-sensitive upgraded Na$_V$1.8 model

The temperature-sensitive upgraded Na$_V$1.8 model retained the four-state conductance structure of the standard Na$_V$1.8 model, with activation, fast inactivation, and two slower availability gates:

$$i_{Na18} = g_{Na18}\, g_{\text{scale,Na18}}(T)\, m^3 \cdot h \cdot s \cdot u\, drv(V_m - E_{Na,T}).$$

(Eq. 60)

Here, $g_{\text{Na18}}$ is the maximum Na$_V$1.8 conductance density in S cm$^{-2}$; $g_{\text{scale,Na18}}(T)$ is dimensionless; and $m$, $h$, $s$, and $u$ are dimensionless state variables. Thus, $i_{\text{Na18}}$ is expressed in mA cm$^{-2}$ when voltage is expressed in mV.

The model receives the numerical temperature $T_c$ in °C. For the equations below, temperature is converted once to the absolute scale:

$$T = (T_c + 273.15)\, K, \qquad TkVm(T) = 1 - \frac{(T - T_{ref})k_{TkVm}}{|V_{rest}|}, \qquad V_m = v\, TkVm(T),$$

(Eq. 61)

Here, $T_c$ is the numerical temperature in °C and $T$ and $T_{\text{ref}}$ are in K; $v$, $V_m$, and $V_{\text{rest}}$ are in mV; and $k_{\text{TkVm}}$ is in mV K$^{-1}$. The factors TkVm and TkENa are dimensionless. The temperature-sensitive upgraded Na$_V$1.8 model uses the minus-sign convention in this channel-local TkVm expression; this convention differs from the general mapping in Eq. 20 and the two expressions should therefore be treated separately.

$$TkENa(T) = 1 + \frac{(T - T_{ref})k_{TkENa}}{E_{Na}}, \qquad E_{Na,T} = E_{Na}\, TkENa(T).$$

(Eq. 62)

Here, $E_{\text{Na}}$ and $E_{\text{Na},T}$ are in mV, and $k_{\text{TkENa}}$ is in mV K$^{-1}$. The mapping is a model-defined temperature correction and is not a dynamic Nernst calculation from sodium concentrations.

For Na$_V$1.8 activation, a small low-temperature facilitation was applied below the channel-specific Kelvin reference temperature:

$$V_{m,m} = V_m - \min\left(0, k^{cold}_{Vshift,m}(T - T_{ref,vshift})\right) + shift_{act}.$$

(Eq. 63)

The original NaV1.8 rate functions were evaluated at $V_{m,m}$ for activation and at $V_m$ for the remaining gates. Here, $\text{shift}_{\text{act}}$ and $k^{\text{cold}}_{\text{Vshift},m}(T - T_{\text{ref,vshift}})$ are in mV. Gate kinetics were then accelerated by channel-specific $Q_{10}$ factors:

$$Q_{10,x}(T) = q_{10,x}^{(T-T_{ref,gate})/(10\,K)}, \qquad \tau_x(T) = \frac{\tau_{x,0}}{Q_{10,x}(T)}, \qquad x \in \{m, h, s, u\}.$$

(Eq. 64)

The temperature difference in the exponent is in K and the denominator is 10 K; $Q_{10,x}$ is dimensionless and $\tau_x$ is in ms.

The NaV1.8-specific conductance scaling was defined from dimensionless empirical relative-current anchors of 0.62 at 283.15 K, 0.82 at 293.15 K, and 1.00 at $T_{\text{ref,amp}} = 303.15$ K:

$$g_{\text{raw,Na18}}(T) = \begin{cases} 0.62\,Q_{10,g,\text{low}}^{(T-283.15\,K)/(10\,K)}, & T < 283.15\,K, \\ 0.62 + (0.020\,K^{-1})(T - 283.15\,K), & 283.15\,K \le T < 293.15\,K, \\ 0.82 + (0.018\,K^{-1})(T - 293.15\,K), & 293.15\,K \le T < 303.15\,K, \\ Q_{10,g,\text{high}}^{(T-303.15\,K)/(10\,K)}, & T \ge 303.15\,K, \end{cases}$$

(Eq. 65)

$$g_{\text{scale,Na18}}(T) = clip\left(\frac{g_{\text{raw,Na18}}(T)}{g_{\text{raw,Na18}}(T_{ref,amp})}, 0.15, 2.50\right), \qquad clip(x, a, b) = \min(\max(x, a), b).$$

(Eq. 66)

Here, $g_{\text{raw,Na18}}$ and $g_{\text{scale,Na18}}$ are dimensionless relative-amplitude factors. The two intervals between the three empirical anchors are linearly interpolated; the low- and high-temperature branches are extrapolated with dimensionless $Q_{10,g,\text{low}}$ and $Q_{10,g,\text{high}}$ factors. The model therefore treats this scaling as a phenomenological NaV1.8 cold-resistance term rather than as a direct measurement of channel abundance. The numerical Kelvin values correspond to the 10 °C, 20 °C, and 30 °C model anchors.

The bounded driving-force function used by the temperature-sensitive upgraded model was:

$$drv(x) = \text{sgn}(x) V_{drv}\left(1 - \exp\left(-\frac{|x|}{V_{drv}}\right)\right).$$

(Eq. 67)

In Eq. 67, $x = V_m - E_{\text{Na},T}$ and $V_{\text{drv}}$ are both in mV; therefore, $\text{drv}(x)$ is in mV. The sign function is dimensionless. Together with $g_{\text{Na18}}$ in S cm$^{-2}$, this gives $i_{\text{Na18}}$ in mA cm$^{-2}$.

Default NaV1.8 parameter values and units are provided in Table S8. All reference temperatures are expressed in K in the equations; their °C equivalents are reported to clarify the corresponding experimental temperature scale.

Total voltage-gated potassium current:

$$i_K = i_{Kf} + i_{Ks}.$$

(Eq. 68)

Fast delayed-rectifier potassium current:

$$i_{Kf} = g_{Kf}\, n_f^4\, (V_m - E_{K,T}), \qquad \frac{dn_f}{dt} = \frac{n_{f,\infty}(V_m) - n_f}{\tau_{n_f}(V_m, T)}.$$

(Eq. 69)

Slow delayed-rectifier potassium current:

$$i_{Ks} = g_{Ks}\, n_s\, (V_m - E_{K,T}), \qquad \frac{dn_s}{dt} = \frac{n_{s,\infty}(V_m) - n_s}{\tau_{n_s}(V_m, T)}.$$

(Eq. 70)

Total K2P current:

$$i_{K2P} = i_{TRAAK} + i_{TREK1},$$

(Eq. 71)

TRAAK-like and TREK-1-like K2P currents:

$$i_{TRAAK} = g_{TRAAK}\, O_{TRAAK}(T, V_m)\, \mathrm{drv}(V_m - E_{L,T})\, Q_{10,L}(T),$$

$$i_{TREK1} = g_{TREK1}\, O_{TREK1}(T, V_m)\, drv(V_m - E_{L,T})\, Q_{10,L}(T).$$

(Eq. 72)

TRAAK first-order relaxation:

$$\frac{dO_{TRAAK}}{dt} = \frac{O_{\infty,TRAAK}(T, V_m) - O_{TRAAK}}{\tau_{TRAAK}(T)}.$$

(Eq. 73)

TRAAK effective gating-charge steady state:

$$O_{\infty,TRAAK}(T, V_m) = \frac{1}{1 + exp\left(\frac{\Delta G_{TRAAK}(T, V_m)}{RT}\right)}, \qquad \Delta G_{TRAAK}(T, V_m)$$
$$= \Delta G_{0,TRAAK}(T) - z_{eff} F (V_m - V_{1/2}).$$

(Eq. 74)

TREK-1 hysteresis-aware relaxation:

$$\frac{dO_{TREK1}}{dt} = \frac{w \cdot O_{\infty,act}(T, V_m) + (1 - w) \cdot O_{\infty,deact}(T, V_m) - O_{TREK1}}{w \cdot \tau_{on}(T) + (1 - w) \cdot \tau_{off}(T)}.$$

(Eq. 75)

Numerical integration and compartmental mapping

For each axonal compartment, local temperature and extracellular potential were sampled from the finite-element fields and supplied to the membrane ODE system (Fig. 2K to M; fig. S4C to E; fig. S11A and B; fig. S12A and B; fig. S13A to E; Tables S5 to S10). Depending on the simulation mode, the ODEs were integrated using fixed-step or adaptive-step solvers, and gating variables were initialized at their steady states under baseline temperature and voltage.

Compartment-level sampling of temperature and extracellular potential:

$$T_c(t) = T(\mathbf{x}_c, t), \qquad V_{e,c}(t) = V_e(\mathbf{x}_c, t).$$

(Eq. 76)

Compartmental membrane equation:

$$C_m \frac{dV_i}{dt} = -\left(i_{Na16} + i_{Na11} + i_{Na18} + i_{Kf} + i_{Ks} + i_{K2P} + i_L\right) + i_{axial},$$

(Eq. 77)

Compartment-specific effective transmembrane voltage:

$$V_m = V_i \cdot TkVm(T_c) - V_{e,c}.$$

(Eq. 78)

Initial gating states for the coupled ODE system:

$$m(0) = m_\infty(V_{rest}),\ h(0) = h_\infty(V_{rest}),\ n_f(0) = n_{f,\infty}(V_{rest}),\ n_s(0) = n_{s,\infty}(V_{rest}),$$

(Eq. 79)

Membrane-potential trajectories were sampled at every node to construct spatiotemporal maps of conduction, block establishment, and onset dynamics. Node-resolved mean membrane-potential maps were computed over predefined analysis windows to classify stable subthreshold depolarization, fluctuating transition states, and propagated action potentials.

### Standardized 150-fiber temperature-responsive digital-twin workflow

Computational overview

We developed a multiscale simulation workflow to test whether local temperature modulation increases the selectivity of electrical control in a heterogeneous myelinated nerve (*53*). The workflow linked four levels: temperature-dependent membrane mechanisms at individual axonal sections, longitudinal cable simulations of single myelinated fibers, a histology-informed transverse nerve cross-section, and compound nerve action potential (CNAP) reconstruction by extracellular summation. Single-fiber dynamics were solved in NEURON, and extracellular stimulation and recording were represented with quasi-static volume-conductor assumptions (*54, 55*).

The model was designed to represent a local thermal control field superimposed on a local electrical stimulation field. Therefore, temperature was assigned locally along each fiber rather than as a single global value. Each axonal section had its own axial coordinate, local temperature, membrane mechanisms, axial resistance, and membrane capacitance. This allowed fibers to be classified not only by electrical recruitment near the stimulation site, but also by whether recruited spikes propagated through the cooled region, were blocked by the cooled region, or changed their onset response during kilohertz-frequency stimulation.

The main population model consisted of 150 myelinated fibers reconstructed from histology-derived morphometric measurements (Tables S5 and S6). The same reconstructed population was reused across activation, local cooling, kilohertz-frequency block, onset-response, rewarming, and CNAP analyses to avoid confounding selectivity comparisons by changes in fiber geometry.

Temperature-dependent membrane model within the standardized fiber workflow

The standardized 150-fiber workflow used the temperature-dependent membrane mechanisms defined in the formula-rich modeling section above, including the effective voltage and reversal-potential temperature mappings, GHK interpretation layer, fast and slow sodium-current modules, K2P-like temperature-sensitive conductances, vtrap-based gating rates, $Q_{10}$-type kinetic scaling, and compartment-level numerical integration. To avoid duplicating the same equations, this section records how those mechanisms were applied to the reconstructed fiber population rather than restating the full formula set (*56*).

Active nodes contained fast transient $Na_V1.6$-like, slow/persistent $Na_V1.1$-like, the temperature-sensitive upgraded $Na_V1.8$ model, and TRAAK-like and TREK-1-like functional components. These terms denote model-defined functional components rather than direct molecular-expression measurements in the rat sciatic nerve. The upgraded $Na_V1.8$ model retained the standard four-state gating structure but added shared voltage/reversal-potential mappings, $Na_V1.8$-specific cold-resistance amplitude scaling, gate-specific $Q_{10}$ kinetics, and bounded driving-force evaluation as defined in Eqs. 60 to 67. Myelinated sections contained temperature-dependent fast and slow potassium mechanisms. The key modeling change relative to fixed-temperature or globally temperature-scaled axon models was that local temperature was assigned to each axonal section, so sodium gating, sodium availability, $Na_V1.8$ current amplitude, K2P-like conductance, axial resistance, and membrane capacitance could vary along a single fiber according to the local cooling field.

Local cooling field

The axial cooling profile was controlled by a dimensionless cooling coefficient, $\mathrm{CP}$. Let $T_{\mathrm{refprof}}(z)$ be the analytic reference temperature profile in Kelvin and $T_0 = 310$ K the baseline reference temperature. The CP-scaled local temperature was:

$$T(z; CP, s_w) = T_0 + CP\left[T_{\text{refprof}}\left(z_c + \frac{z - z_c}{s_w}\right) - T_0\right],$$

(Eq. 80)

where $z$ is the section-center coordinate, $z_c$ is the cooling-center coordinate, and $s_w$ is the axial width-scale factor. $\text{CP} = 0$ gives the no-cooling baseline, whereas larger CP values impose stronger cooling. The width-scale factor changes the axial spread of the cooling profile.

For each section, the local temperature $T(z; \text{CP}, s_w)$ was assigned to every temperature-dependent mechanism in that section. Axial resistance and membrane capacitance were also rescaled from their baseline values:

$$R_a(T) = R_{a,0}\frac{f_R(T)}{f_R(T_{\text{base}})}, \qquad C_m(T) = C_{m,0}\frac{f_C(T)}{f_C(T_{\text{base}})}.$$

(Eq. 81)

The implemented scaling functions were:

$$f_R(T) = \max\left(0.05, 1 + \frac{T - 298.15}{50}\right), \qquad f_C(T) = \max\left(0.05, 1 + \frac{T - 310.15}{80}\right),$$

(Eq. 82)

with $T$ in Kelvin. In the representative 150-fiber mixed-population simulation, the cooling center was at 2.8 cm, $\text{CP} = 1.0$, and $s_w = 2.0$, giving a minimum local temperature of 8.44 °C. This is the steady-state temperature exported from the thermal field used as the representative input for the 150-fiber population model. It should not be conflated with the 7 °C and 10 °C values in Fig. 2E, which are results of the transient finite-element thermal-field comparison.

Single-fiber axon models

Each reconstructed fiber was assigned to one of two myelinated-fiber model families based on equivalent outer fiber diameter (Table S6). Fibers with diameter <5.7 µm used a self-built Peña-style small-diameter branch, whereas fibers with diameter ≥5.7 µm used a self-built MRG-style branch. This hybrid strategy was used because small myelinated fibers require ultrastructural scaling outside the standard large-fiber MRG parameter range, whereas MRG-style models remain the standard framework for larger mammalian myelinated fibers (*5, 46, 57*).

The Peña-style branch used diameter-dependent equations to derive axon diameter, node diameter, paranodal length, internodal distance, and number of myelin lamellae. The MRG-style branch retained the canonical node-paranode-juxtaparanode-internode compartmental sequence, with geometry interpolated across tabulated MRG diameters and optionally constrained by histology-derived axon diameter, g-ratio, or myelin thickness. In both branches, the repeated longitudinal unit consisted of an active node, paranodal myelin attachment segments, juxtaparanodal segments, and internodal segments. The number of nodes was selected to span the 5.1-cm modeled nerve length, with passive end nodes included to reduce boundary artifacts.

Initial-state replay was used to reduce artifacts from starting cooled simulations from a naive resting state. Pre-stimulus states from 40-ms reference simulations were mapped to each fiber by model family, nearest diameter, section kind, and normalized axial position before the stimulus was applied.

Histology-informed nerve cross-section

The transverse nerve population was reconstructed from histology-derived segmentation metrics. The source morphometric table contained equivalent outer fiber diameter, equivalent axon diameter, myelin thickness, g-ratio, axon and fiber centroids, and source labels for myelinated fibers (Tables S5 and S6). The source table contained 1,944 eligible fibers, all of which passed the reconstruction filters of positive myelin thickness and equivalent outer diameter ≤16 μm.

A compact representative population of 150 fibers was selected by spatial-diameter stratified sampling. Source fibers were binned by radial position, angular sector, and diameter quantile. Sampling was performed to preserve the joint distribution of fiber diameter and transverse position rather than sampling diameters independently of location. This was necessary because stimulation, cooling, propagation, and recording all depend on both fiber size and transverse distance from the electrode.

The sampled fibers were placed at their recentered empirical centroids with native scale and no artificial redistribution. The intraneural radius was 600 μm. The reconstructed sample remained within this boundary, with a maximum center radial position of 350.02 μm and a maximum outer-fiber edge radial position of 355.03 μm. No equivalent-circle overlaps were present in the reconstructed representation.

Fibers were grouped for analysis into Aδ fibers (<4 μm), mixed-diameter fibers (4 to <7 μm), and Aα/β fibers (≥7 μm). The reconstructed population contained 38 Aδ fibers, 82 mixed-diameter fibers, and 30 Aα/β fibers.

Extracellular stimulation and recruitment detection

Electrical activation used a bipolar point-source pair in a homogeneous isotropic extracellular medium with conductivity 1.0 S/m. The intraneural radius was 600 μm, the epineurium thickness was 100 μm, and stimulation contacts were positioned at a radial distance of 800 μm. The stimulation center was at 2.0 cm, with 1.0-mm inter-contact spacing. The waveform was a symmetric biphasic pulse with 0.05-ms phase width, zero interphase gap, and onset at 10 ms. Current amplitude was swept from 0.5 to 6.0 mA in 0.5-mA increments (Table S2).

The reported current amplitudes correspond to distinct modeling and experimental layers and were not pooled or treated as interchangeable conditions. The 0.5- to 6.0-mA range was the mixed-population recruitment sweep, and 6.0 mA was the representative mixed-population point in Fig. 2N. The 10-mA conditions in fig. S19 were separate fixed-current CNAP reconstructions at the labeled geometries and cooling-profile values rather than part of the 0.5- to 6.0-mA recruitment sweep. The 4-mA conditions in fig. S4D and E and fig. S13B and D were panel-specific neural finite-element simulation settings. In vivo low-frequency CNAP recordings used 400 μA with 50-μs phases unless otherwise stated, whereas fig. S16 used a 50- to 800-μA amplitude series to test waveform and polarity robustness (Table S2).

The activation threshold of a fiber was defined as the lowest current amplitude that produced at least one action potential at a detection point 1 mm distal to the stimulation center. Additional propagation stations were used to separate local recruitment from propagation across the cooled region: a proximal reference station at 1.4 cm, an intermediate station between the stimulation and cooling centers at 2.45 cm, and a distal recovery station defined as the first point distal to the cooling center where the temperature recovered to 35 °C. For the representative $\mathrm{CP} = 1.0$, $s_w = 2.0$ condition, this distal station was 4.706 cm.

Kilohertz-frequency block and onset-response classification

Kilohertz-frequency stimulation simulations used the same fiber geometries and temperature-assignment procedure. Block/pass status and onset pattern were treated as orthogonal classifications. A fiber could be blocked or pass the block region, and independently could show no propagating onset, transient propagating onset, or persistent propagating onset.

KHFS frequencies were retained as panel-specific conditions. The in vivo onset experiment and the primary baseline simulation comparisons used 10 kHz, whereas selected cooling-assisted simulation panels used 20 kHz to test whether onset suppression and block behavior generalized beyond the primary frequency. The 10- and 20-kHz results therefore represent separate conditions and were not pooled as a single stimulation setting; exact frequencies follow the corresponding panel labels (Fig. 3A to E; fig. S12D, H to J, and M; fig. S13B and D).

Propagating onset events were not defined solely by a membrane-potential crossing near the KHFS electrode, because local high-frequency stimulation can produce field-induced depolarization near the stimulation center without launching a true propagating action potential. Instead, onset classification required action-potential-like amplitude and propagation trajectory at remote detection positions away from the KHFS center. This criterion reduces false-positive persistent-onset classifications caused by local subthreshold depolarization.

CNAP reconstruction

CNAPs were reconstructed by linear superposition of single-fiber action potentials (SFAPs). For each fiber $i$, probe $p$, current amplitude $I$, and time $t$, let $V_{\mathrm{SFAP},i,p}(I,t)$ be the simulated single-fiber contribution and $a_i(I)$ be the activation indicator for that fiber and current amplitude. The compound waveform was:

$$V_{\mathrm{CNAP},p}(I,t) = \sum_{i=1}^{N} a_i\,(I) V_{\mathrm{SFAP},i,p}(I,t).$$

(Eq. 83)

Class-specific CNAP components were computed by restricting the summation to each fiber class. The total CNAP therefore equals the direct sum of the component CNAPs when the same activation indicators, recording probe, and current amplitude are used. This summation does not assume that axonal dynamics are linear. The nonlinear membrane dynamics were solved separately for each fiber; only the extracellular recording potentials were summed linearly under the quasi-static volume-conductor approximation.

The standard production recording geometry used monopolar point-source reciprocity at probes located at 800-μm radial distance on both sides of the nerve. Additional ring-electrode visualizations were produced from saved voltage traces using a 270° ring projection with 100-μm axial width. These ring reconstructions were used for visualization and recording-geometry comparison, whereas the primary CNAP calculation was the SFAP summation in Eq. 83.

Standardized modeling parameter summaries incorporated into final Tables S2-S10

The device, cooling-field, fiber-population, and active-node parameter summaries used in the standardized workflow are provided in Tables S2, S3, S5, S6, and S9. The tables preserve the values and source notes used by the model; the equations and implementation assumptions are defined in the sections above.

Data, code, and references for modeling

Modeling data, code, and reference entries are merged into the global Data and code availability and References sections below.

Methodological limitations specific to the temperature-responsive fiber workflow

The ion-channel components are phenomenological temperature-sensitive representations. They extend standard MRG/Peña-style myelinated-fiber models by adding local temperature mappings, a temperature-sensitive upgraded $\mathrm{Na_V}$1.8 model, K2P-like conductances, and temperature-dependent cable properties. The $\mathrm{Na_V}$1.6-like, $\mathrm{Na_V}$1.1-like, TRAAK-like, and TREK-1-likecomponents should therefore be interpreted as model-defined functional representations rather than direct molecular reconstructions of a specific experimental preparation.

The extracellular stimulation field used in the primary activation simulations was a homogeneous isotropic point-source approximation. This representation supports controlled comparisons across fiber class, CP, electrode spacing, and temperature profile, but it does not include a full finite-element representation of cuff insulation, anisotropic endoneurium, perineurium, fascicular boundaries, or electrode-tissue interface impedance.

Fibers were simulated independently. The model does not include ephaptic coupling, activity-dependent extracellular ion accumulation, vascular feedback, or thermal feedback from stimulation. CNAP reconstruction assumes linear superposition of extracellular potentials after the nonlinear membrane dynamics of each fiber have been solved.

The 150-fiber population is a representative computational subset rather than an exhaustive enumeration of the original histology-derived nerve section. Its role is to preserve the joint distribution of fiber size and transverse position in a population that can be simulated repeatedly across many stimulation and cooling conditions.

CP is a dimensionless cooling-control coefficient that scales an analytic temperature profile. It should be interpreted as a simulation control parameter linked to local minimum temperature and axial thermal width, not as a direct physical measurement of cooling-device power unless separately calibrated to thermal measurements or a thermal transport model.

### Quantification and statistical analysis

Summary statistics and error bars are specified in the corresponding figure legends. Temperature data, electrochemical readouts, ankle-angle changes, and CNAP amplitude and latency measurements were quantified as defined in the main text and corresponding legends. CNAPs were analyzed using peak-to-peak amplitude and peak latency, and fast and slow components were distinguished by baseline propagation delay and waveform morphology. For each rat, three consecutive cycles were recorded under the same experimental condition and averaged to obtain one value per animal; per-animal values from three rats were used as independent biological replicates for group-level analysis. In Fig. 5F, G, J, and K, the plotted values are the means of three repeated cycles from one rat, with SD error bars. In Fig. 5H, I, L, and M, each temperature was compared with the lowest-temperature value within the same recording direction and CNAP component using a two-sided paired t test. The onset response was quantified as the transient ankle-angle deflection immediately after KHFS onset. Statistical annotations are defined as follows: ns, not significant; *P < 0.05; **P < 0.01; ***P < 0.001. For histological analyses, the animal was the biological replicate. Multiple fields, section planes, or ROIs from the same animal were averaged within animal before group-level comparison. Histological data are reported as mean ± SD, with individual biological replicates shown as points. Overall differences among the untreated control, acute-stimulation, and chronic-implantation groups were tested using a two-sided Kruskal–Wallis test. Pairwise comparisons used two-sided Mann–Whitney U tests with Holm correction for multiple comparisons. $P < 0.05$ was considered statistically significant, and ns denotes no statistically significant difference. Statistical analysis and plotting were performed locally in Python 3.13.5. Unless otherwise stated, the histology analyses used $n = 3$ animals per group.

### Supplementary Text

Supplementary Note 1. Engineering rationale and iterative development of NeuroSwitch

This note explains the design progression represented in figs. S1 to S7 and Tables S1 to S4. The first-generation cuff placed the centers of cooling and low-frequency activation only 4 mm apart. During low-frequency stimulation, the afferent CNAP disappeared when the measured core nerve temperature reached approximately 19 °C (fig. S2A to H). This temperature was substantially higher than the sub-10 °C range generally associated with direct cold conduction block *(6, 38)*, indicating that the overlapping cooling field suppressed neural recruitment at the stimulation site before it blocked action-potential propagation. This unexpected response motivated a design in which recruitment filtering and conduction block were separated spatially.

The final architecture was organized around four functional requirements. The cuff had to generate a low-temperature core along the nontarget propagation pathway, retain a warmer gradient near the drive site, confine KHFS to the block region, and provide a local temperature readout. A microfluidic channel generated the cooling field; tripolar block electrodes localized KHFS near its center; low-frequency drive electrodes were displaced along the nerve axis; and an integrated thermistor monitored the block-zone temperature. General electrode and charge-delivery considerations followed established neural-interface principles *(58, 59)*, whereas the KHFS layout addressed onset and field-spread constraints associated with kilohertz nerve block *(9, 60–62)*.

The diamond-filled PDMS layer addressed the thermal resistance of an otherwise compliant silicone interface. Retaining PDMS provided conformity and microfluidic compatibility, but unfilled PDMS limited conductive heat transfer between the channel and the nerve-facing surface. The 50 wt.% diamond-filled composite increased thermal conductivity and diffusivity while preserving the flexible cuff architecture. In the finite-element comparison, the nerve-facing temperature reached approximately 7 °C with the composite and approximately 10 °C with unfilled PDMS, and the time required to approach the same low-temperature target decreased from approximately 290 s to approximately 20 s (Fig. 2E and F; fig. S3F and G). These measurements support the composite as a thermal-interface component rather than as an isolated materials modification.

Electrical-field simulations and multi-spacing experiments then constrained the spatial layout. Tripolar KHFS produced a more localized blocking field than the broader bipolar configuration, reducing direct interference with the low-frequency drive site (fig. S4D and E). In vivo measurements showed that interference was eliminated only when the center-to-center distance between the tripolar block and bipolar drive sites exceeded 7.5 mm (fig. S2J to N). This result supported the 9-mm spacing of the final experimental cuff and the 8-mm representative spacing used in the 150-fiber population model (Table S2). Thus, the final geometry arose from the initial recruitment-suppression observation and subsequent thermal, electrical-field, and spacing constraints.

Supplementary Note 2. Functional roles of the cooling core and surrounding temperature gradient

Spatially graded cooling served two separable functions in the block–drive configuration (Fig. 1F; Fig. 5D to M; fig. S16). The low-temperature core lay along the nontarget propagation pathway and could block recruited activity that crossed the cooling center. The warmer shoulder of the temperature field overlapped the drive region, where it changed recruitment without requiring the local temperature to reach the direct cold-block range. Directional bias therefore arose primarily from conduction block along the nontarget pathway, whereas fiber bias arose primarily from temperature-dependent recruitment near the drive site.

This distinction is important for interpreting the retained CNAP. Cooling strongly attenuated the efferent quick component, consistent with block of the muscle-directed Aα pathway, while partially preserving the afferent quick component and suppressing the Aδ-enriched slow component in both directions. The resulting signal is therefore described as a directional, Aβ-biased afferent output rather than as ordinary unidirectional stimulation or complete isolation of a single fiber class. Localization of the tripolar KHFS field complements this thermal organization by preventing the electrical block field from directly dominating the spatially separated drive region.

Supplementary Note 3. Interpretation of the temperature-responsive neural model and CNAP readouts

This note defines how the temperature-responsive digital twin should be interpreted in relation to Figs. 2 to 5, figs. S8 to S17, and Tables S5 to S9. Conventional fixed-temperature models, or models that apply only a single global $Q_{10}$ kinetic-scaling factor, cannot represent a warm drive site, a temperature-gradient region, and a cold block core along the same fiber. The present model therefore assigned temperature locally to axonal sections, allowing recruitment, propagation delay, and conduction failure to be evaluated at their respective positions. This treatment is consistent with the established effects of cooling on peripheral-nerve excitability and conduction *(63)*.

The temperature-responsive channel terms are model-defined functional components rather than measurements of molecular expression in the rat sciatic nerve. $Na_V1.6$-like, $Na_V1.1$-like, and the upgraded $Na_V1.8$ components represent distinct $Q_{10}$-scaled sodium-current kinetics, whereas TRAAK-like and TREK-1-like terms represent thermosensitive K2P contributions to excitability *(49, 64–68)*. The hybrid Peña-style and MRG-style fiber families extend the simulated diameter range needed to represent smaller, slower fibers and larger, faster myelinated fibers within the same reconstructed population *(42, 53, 57)*. Their equations, conductance sets, compartment assignments, and calibration states are specified in the Methods and Tables S5 to S9.

Precooling was implemented as a thermal trajectory rather than as an instantaneous change in a temperature parameter. Each fiber entered KHFS from a cooling-conditioned membrane state, enabling the model to distinguish local or nonpropagating responses from transient and persistent propagated onset. This distinction connects the simulated state transition to the functional ankle-angle measurement: a local field-induced depolarization that does not leave the block region is not equivalent to an onset event that propagates to the motor pathway *(9, 60–62)*.

For CNAP reconstruction, nonlinear single-fiber dynamics were solved first and the extracellular contributions were then summed linearly. Decomposition by diameter showed that early, short-latency components were enriched in larger and faster Aα/Aβ fibers, whereas delayed components received stronger contributions from smaller and slower Aδ-range fibers. These peaks remain compound readouts: their composition depends on diameter, conduction velocity, stimulation amplitude, field geometry, and recording position. Accordingly, "Aα/Aβ-enriched" and "Aδ-enriched" indicate relative contributions rather than one-to-one identification of fiber types. This model-to-experiment mapping supports interpretation of recruitment filtering and directional propagation without claiming exact molecular or whole-nerve reconstruction. Detailed numerical assumptions and boundary conditions are retained under "Methodological limitations specific to the temperature-responsive fiber workflow."

**Supplementary Figure Legends**

Each supplementary figure is paired with its current image and panel-level legend.

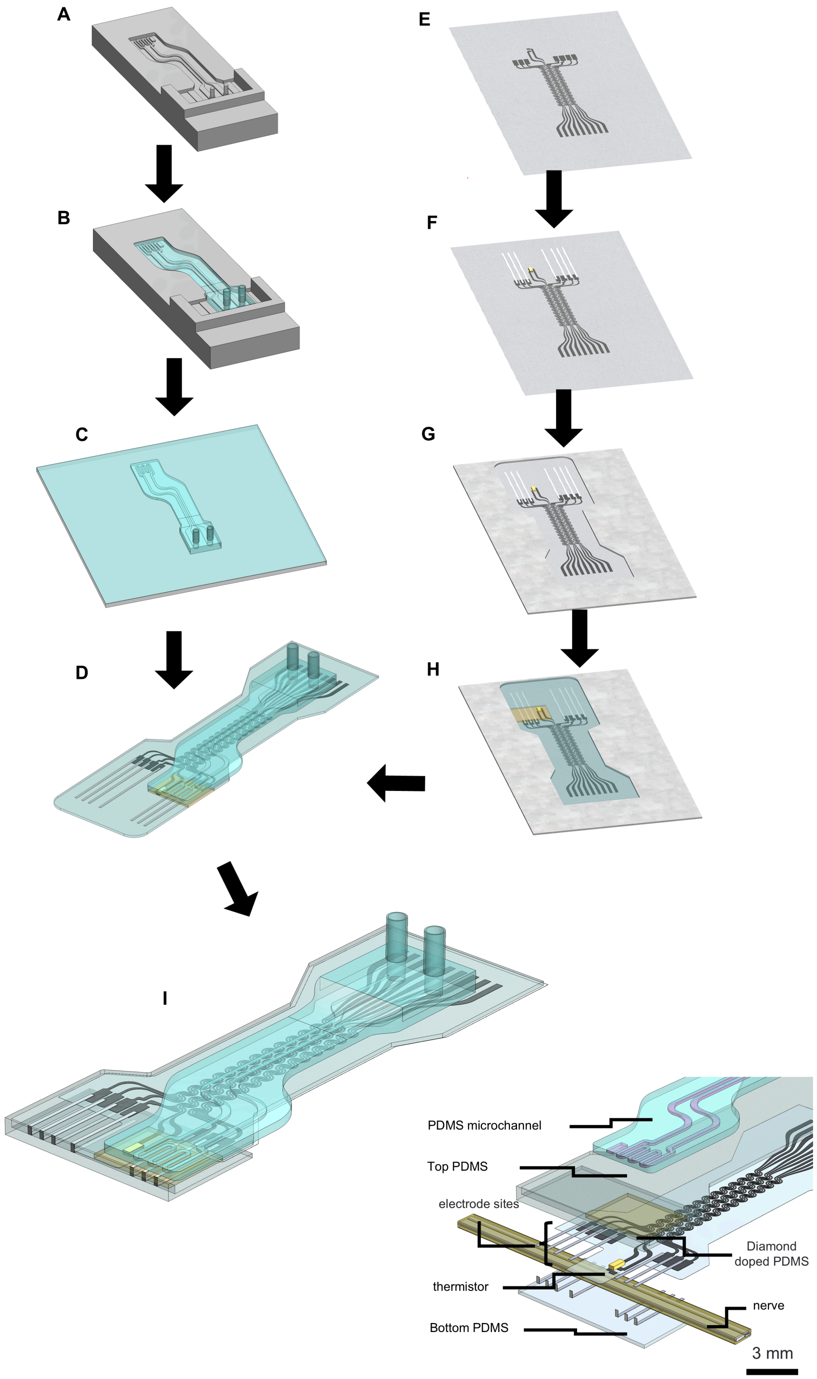


Fig. S1. Fabrication workflow and layer architecture of the NeuroSwitch cuff (A) Three-dimensional printed mold for the microfluidic channel. (B) Polydimethylsiloxane (PDMS)

casting and curing to form the microfluidic layer. (C) Bonding of the microfluidic layer to a 50-μm PDMS sealing membrane. (D) Alignment and bonding of the microfluidic and electrode layers. (E) Laser patterning of the Ti interconnects. (F) Attachment and electrical connection of the Pt electrode sites and thermistor with silver epoxy. (G) Fixture used to define the final device contour. (H) Encapsulation with diamond-filled silicone (yellow) and PDMS. (I) Formation of the leading hook and final device assembly. (J) Exploded view of the device head and constituent layers. Related to Fig. 1.

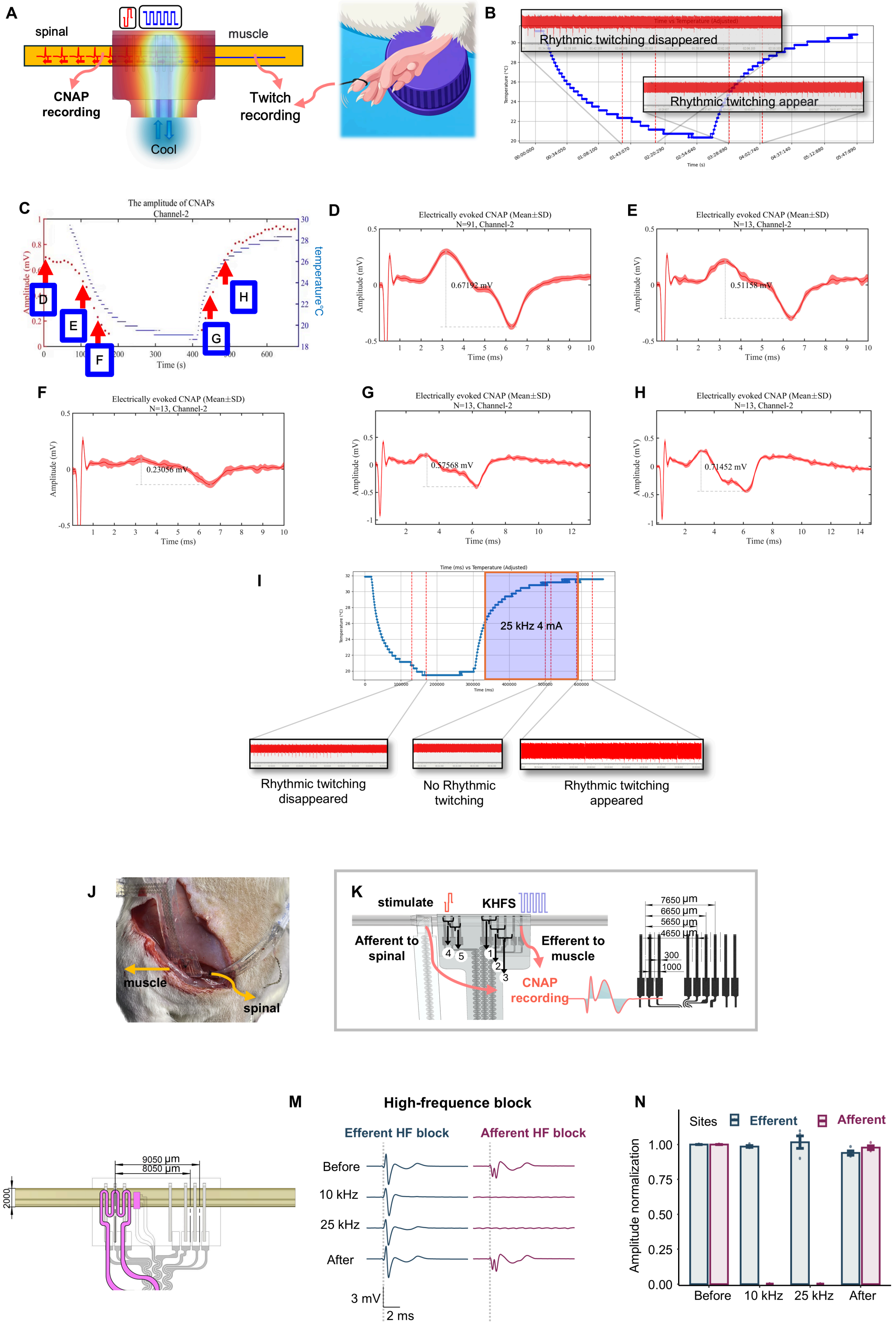


Fig. S2. Low-interference block–drive spacing enables directional neural control (A) Schematic of the testing configuration using the version 1.0 device. Intermittent low-frequency electrical stimulation (50 μs, 400 μA biphasic square pulses) was applied, while afferent CNAPs and toe twitching were recorded sequentially. (B) Toe-twitch amplitude as a function of temperature. Twitching disappeared when the minimum nerve temperature (temperature in the non-activation region) decreased to approximately 21 °C and reappeared at approximately 24 °C during rewarming. (C) Maximum afferent CNAP amplitude recorded with the version 1.0 device during cooling and rewarming. CNAPs disappeared when the core nerve temperature reached approximately 19 °C, indicating that cooling directly suppressed neural recruitment by the stimulation electrode rather than blocking action-potential propagation. (D to H) Representative CNAP waveforms recorded at the indicated time points in panel C; labels N=91 and N=13 denote the numbers of stimulation cycles. (I) Continuous transition of the version 1.0 device from cooling-mediated block to KHFS. At physiological temperature, KHFS similarly directly suppressed neural recruitment within the activation region. (J) Implantation of the multi-spacing test cuff, with afferent and efferent propagation directions indicated. (K) Experimental configuration in which a single cuff provided four block–drive spacings. Efferent CNAPs were recorded during low-frequency drive and KHFS block. Experimental measurements showed that interference was eliminated only when the distance between the center of the tripolar KHFS block and the center of the bipolar activation site exceeded 7.5 mm. (L) Parameterized NeuroSwitch layout derived from experimental interference mapping and simulation. Candidate minimum block–drive spacings exceed the longest experimentally observed interference distance. (M) Representative afferent and efferent CNAP waveforms before, during, and after KHFS. (N) Directional selectivity quantified as the mean normalized afferent and efferent CNAP amplitudes across three rats. Error bars indicate SEM; no inferential statistical comparison was performed. Related to Fig. 2.

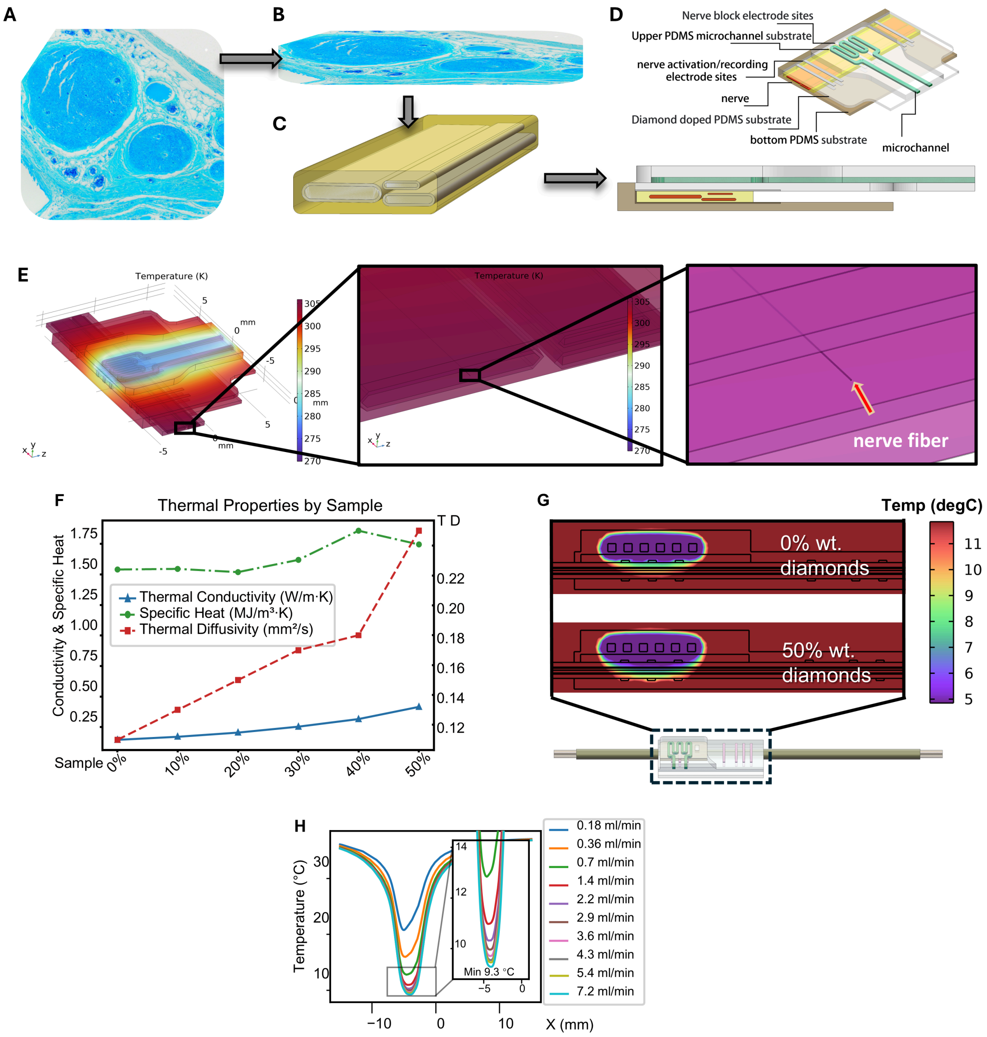


Fig. S3. Device-nerve geometries and material parameters used in thermal simulations (A) Luxol fast blue-stained cross-section of rat sciatic nerve. (B) Virtually compressed nerve cross-section derived from the histological image. (C) Three-dimensional flattened-nerve geometry reconstructed from the compressed section. (D) Three-dimensional implantation geometry of the deformed nerve within the NeuroSwitch cuff, including the electrode, microchannel, and diamond-filled thermal-interface layers. (E) Representative fiber location within the three-dimensional model and local temperature field. (F) Thermal conductivity, specific heat, and thermal diffusivity of PDMS composites containing 0 to 50 wt.% diamond. (G) Finite-element comparison of the device-nerve temperature field with undoped PDMS and 50 wt.% diamond-filled PDMS. Related to Figs. 1 and 2. (H) Simulated axial temperature profiles along the lower nerve surface at different coolant flow rates (inlet temperature, −10 °C).

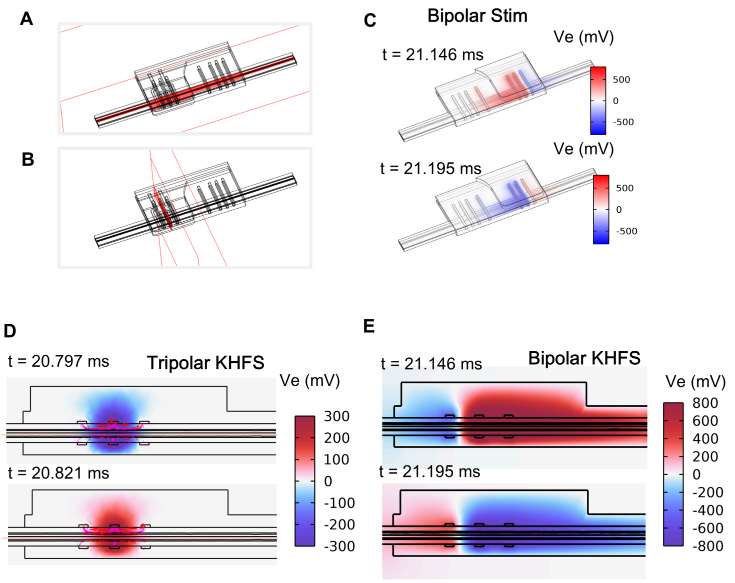


Fig. S4. Section definitions and extracellular-potential fields used in cooling-assisted electrical simulations (A) Transverse section selected through the NeuroSwitch cuff. (B) Longitudinal section selected along the cuff and nerve. (C) Simulated extreme-value extracellular-potential distribution during bipolar stimulation. (D) Extracellular-potential distribution during tripolar KHFS (10 kHz, 4 mA) at a representative peak-amplitude time point; streamlines and arrows indicate current paths and direction. (E) Corresponding distribution during bipolar KHFS (10 kHz, 4 mA). Related to Figs. 1 and 2.

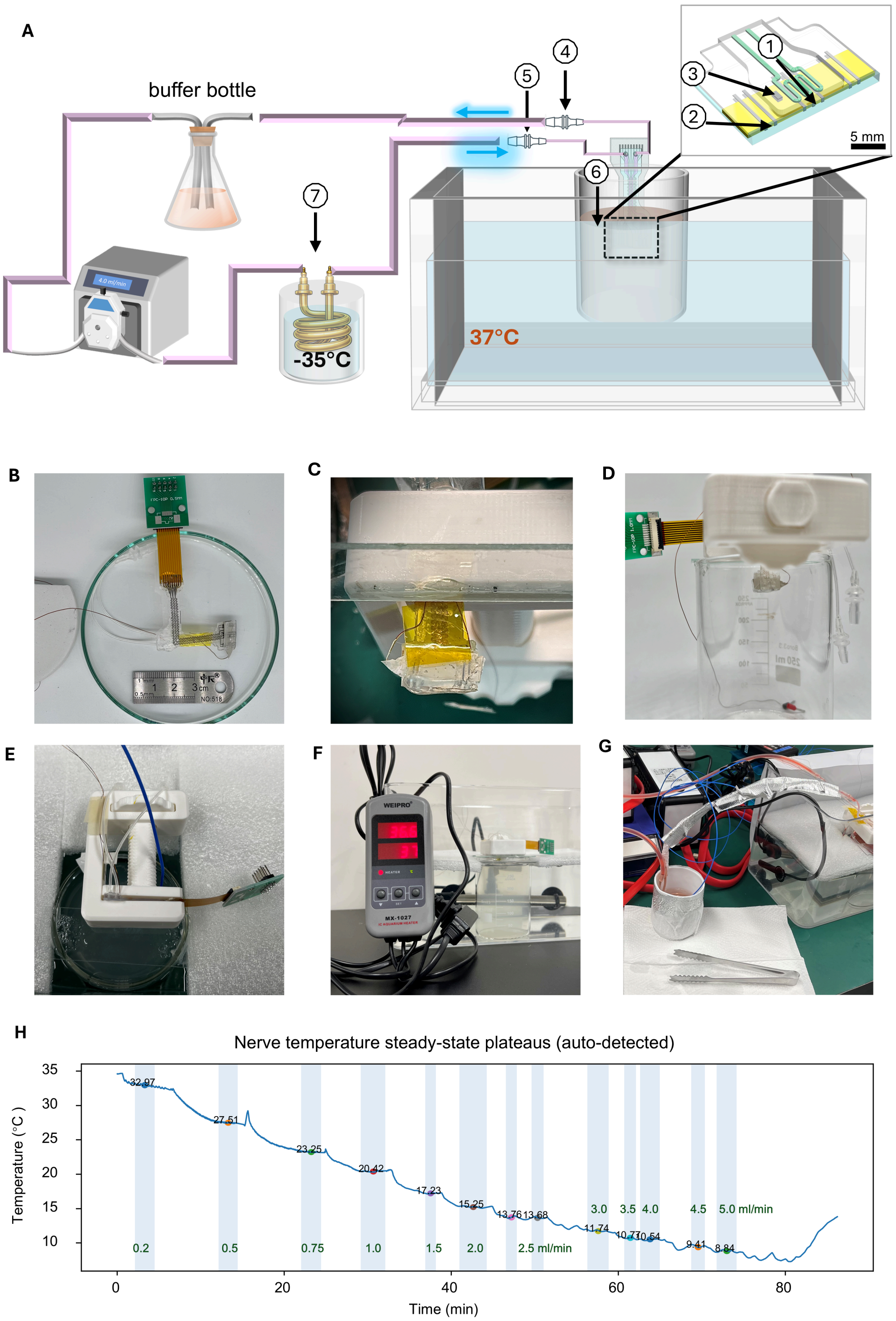


Fig. S5. Experimental platform for thermal characterization of the cooling module (A) Schematic of the cooling-test system, including the pump, precooling bath, tissue-mimicking bath, device, and temperature-measurement sites. (B to G) Photographs of the device, phantom-nerve platform, coolant circuit, temperature readout, and complete thermal-characterization setup. (H) Temperature windows used to calculate the steady-state nerve temperature at each flow-rate plateau. Related to Fig. 2.

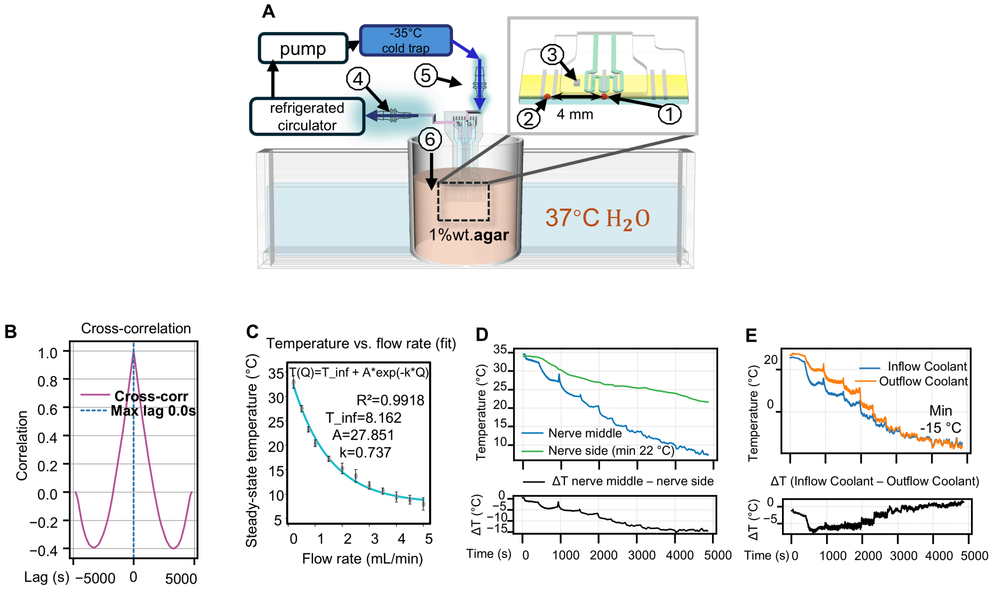


Fig. S6. Thermal regulation and temperature sensing by the NeuroSwitch cuff (A) Cooling-test platform and six temperature-measurement sites. (B) Cross-correlation between intraneural and integrated-sensor temperature signals, showing their temporal alignment and stable offset. (C) Steady-state intraneural temperature as a function of coolant flow rate, with exponential fitting. (D) Spatial confinement of cooling, comparing temperature trajectories at the nerve center (site 1) and 4 mm lateral to the center (site 2). (E) Inlet (site 5) and outlet (site 4) coolant temperatures during stepwise changes in flow rate, together with the inlet-outlet temperature difference ($\Delta$T).

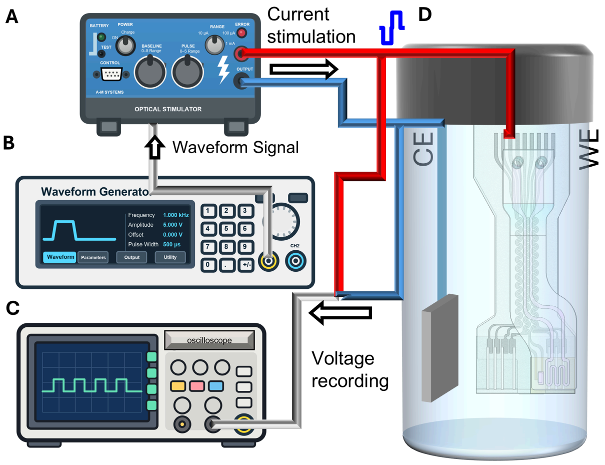


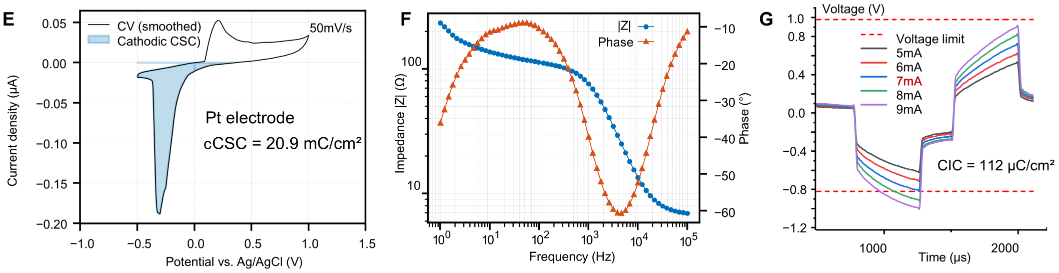


Fig. S7. Electrochemical characterization and charge-injection-capacity measurement of the cuff electrodes (A) Circuit used to measure charge injection capacity (CIC). (B) Waveform-generator and isolated-stimulator connections used to deliver charge-balanced biphasic pulses. (C) Oscilloscope configuration used to record electrode polarization. (D) Electrochemical test cell used for the measurements, showing the counter electrode (CE) and working electrode (WE). (E) Cyclic-voltammetry (CV) curves and cathodic charge-storage-capacity calculation in phosphate-buffered saline (PBS; pH 7.4) with an Ag/AgCl reference electrode at 50 mV/s. (F) Electrochemical-impedance-spectroscopy (EIS) Bode plots measured under the same electrolyte and reference-electrode conditions. (G) Biphasic voltage transients and polarization limits used to estimate CIC.

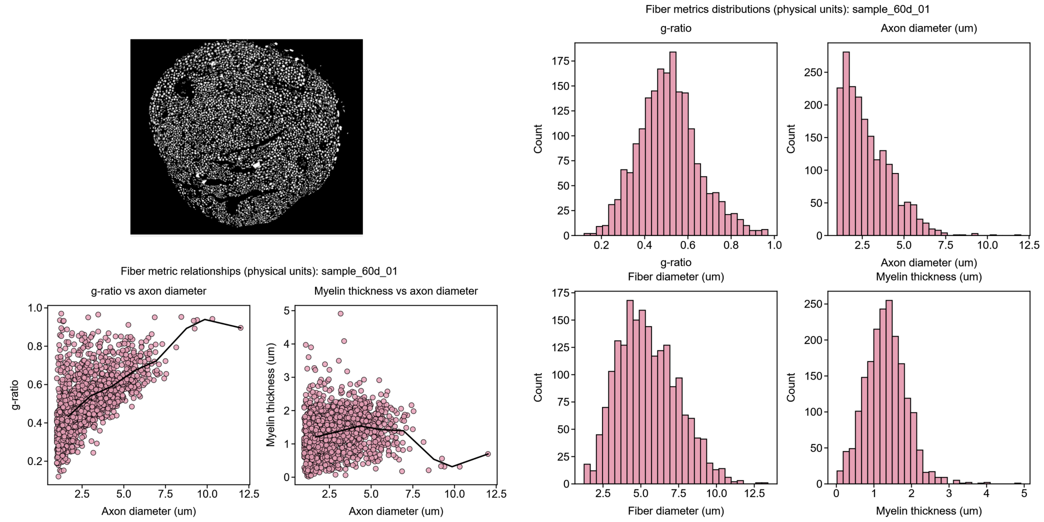


Fig. S8. Morphometric distributions of myelinated fibers used to construct the 150-fiber population A representative segmented sciatic-nerve cross-section is shown together with fiberlevel relationships between g-ratio and axon diameter and between myelin thickness and axon diameter. Histograms summarize the distributions of g-ratio, axon diameter, total fiber diameter, and myelin thickness in the source nerve bundle. These measurements define the anatomical sampling space for the 150-fiber mixed-nerve model.

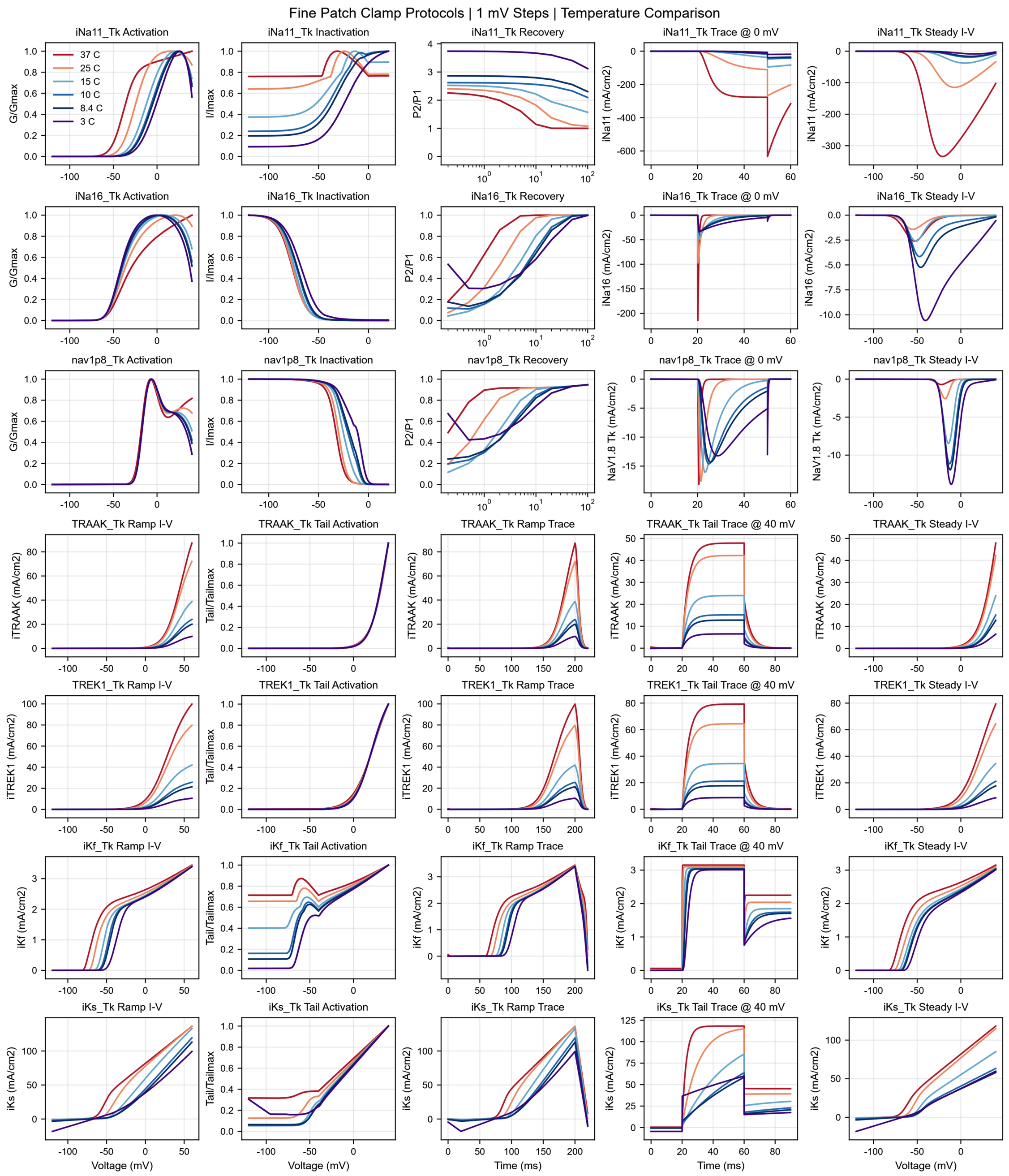


Fig. S9. Temperature-dependent responses of the temperature-sensitive upgraded ion-channel models Fine-patch-clamp simulations with 1-mV voltage steps compare channel behavior from

37 °C to 2 °C. Rows show the temperature-coupled persistent $Na^+$, transient $Na^+$, NaV1.8, TRAAK, TREK-1, fast $K^+$, and slow $K^+$ components. Columns summarize activation, inactivation or recovery, representative voltage-clamp traces, and steady-state current-voltage relations. The coordinated temperature dependence of gating and current amplitude provides the membrane-level basis for the cooling-assisted electrical fiber model. Related to Fig. 2.

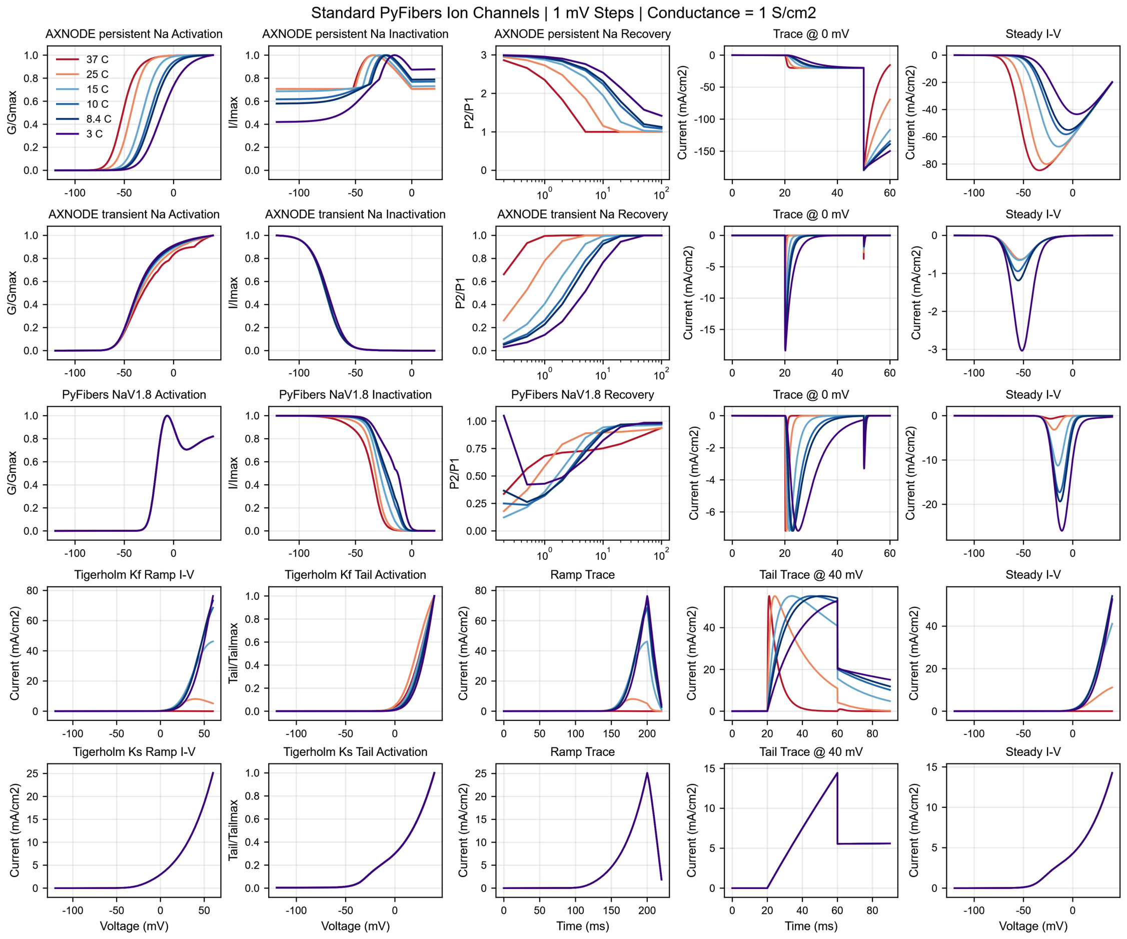


Fig. S10. Temperature responses of standard ion-channel formulations without calibrated thermal sensitivity Standard PyFibers formulations were evaluated with 1-mV voltage steps and a nominal conductance of 1 S cm$^{-2}$ across the same temperature range used for the temperature-sensitive upgraded models. Rows show persistent and transient nodal $Na^+$ currents, PyFibers $Na_V1.8$, and the Tigerholm Kr and Ks components; columns show activation, inactivation or recovery, representative clamp traces, and steady-state current-voltage relations. The incomplete or inconsistent thermal responses illustrate why the standard formulations were not sufficient for the temperature-coupled digital twin. Related to Fig. 2.

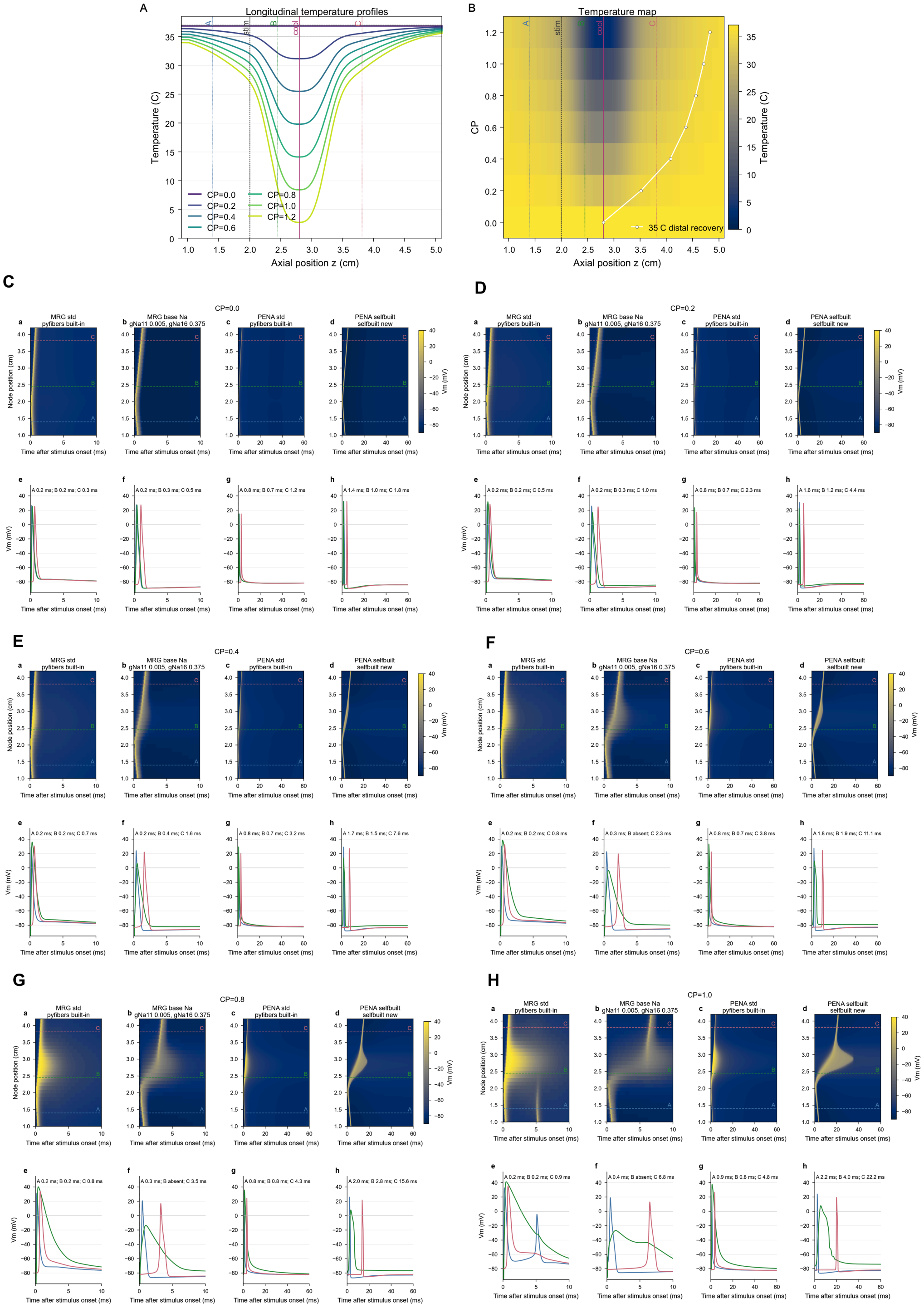


Fig. S11. Spatial cooling profiles reveal temperature-dependent action-potential propagation in standard and temperature-sensitive upgraded fiber models (A) Longitudinal nerve-temperature profiles generated by scaling the finite-element cooling field with the cooling-profile parameter (CP). CP = 1 corresponds to the exported three-dimensional temperature field; other values proportionally scale the temperature reduction relative to 37 °C. (B) Temperature map across CP and axial position, with the distal recovery path to 35 °C indicated. (C to H) Spatiotemporal membrane-potential maps and representative traces for structure-matched standard and temperature-responsive fibers at CP = 0, 0.2, 0.4, 0.6, 0.8, and 1.0. Each condition compares 10-μm MRG fibers and 4-μm PENA fibers and distinguishes action-potential initiation, slowing, propagation, and cold block. Related to Fig. 2M.

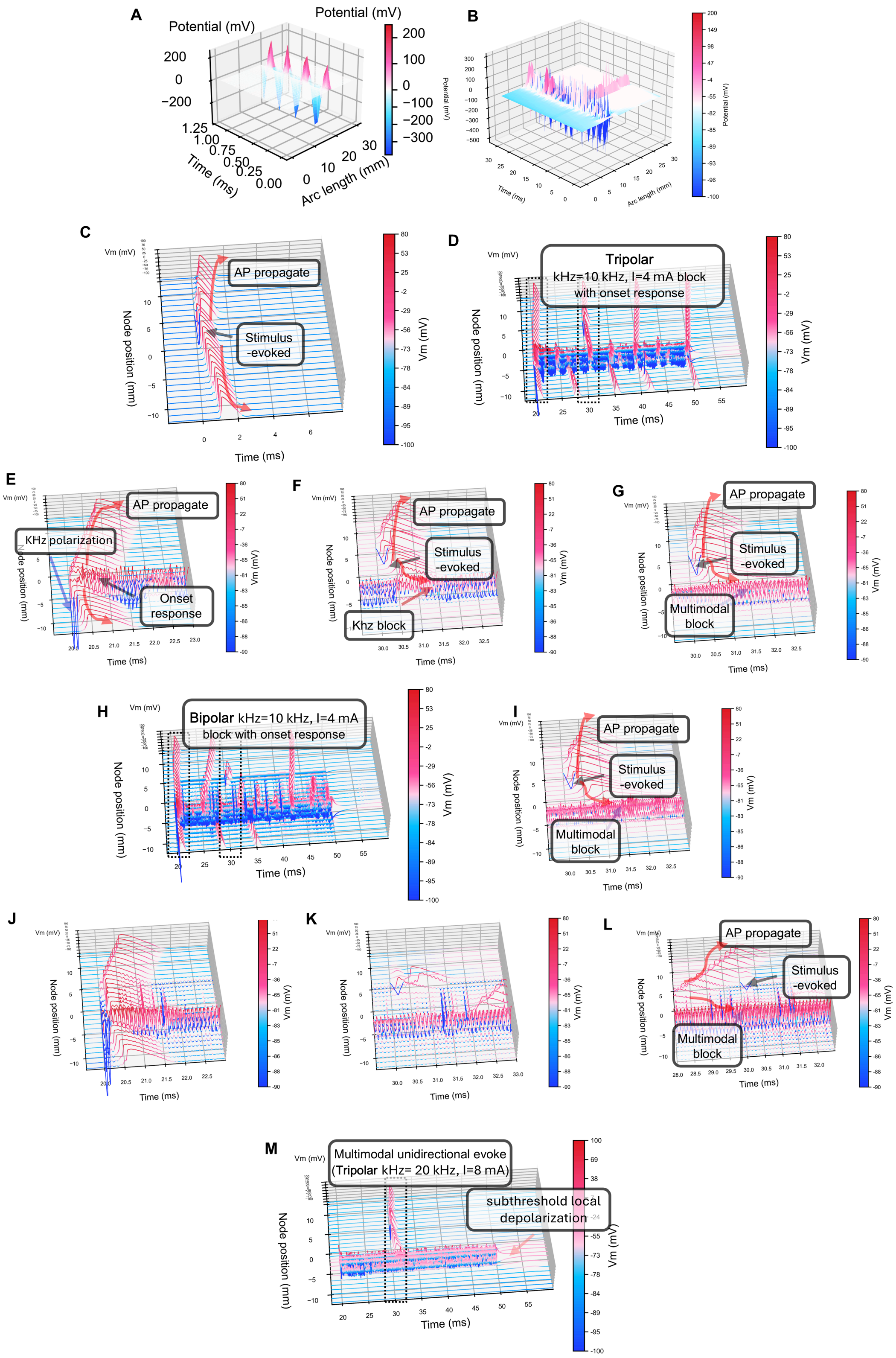


Fig. S12. Multiphysics simulations resolve membrane dynamics during cooling-assisted KHFS block (A) Short-time view of the spatiotemporal extracellular potential generated by tripolar

KHFS. (B) Propagation map showing interruption of an action potential by tripolar KHFS. (C) Action-potential initiation and propagation at baseline temperature. (D) Membrane-potential dynamics during tripolar KHFS (10 kHz) at baseline temperature, showing block preceded by a prominent onset response. (E) Enlarged view of the tripolar-KHFS onset phase. (F) Enlarged view of the tripolar-KHFS steady-state phase. (G) Combined cooling and 10-kHz tripolar KHFS reduces onset activity and produces smoother stabilization. (H) Bipolar KHFS (10 kHz) at baseline temperature produces persistent onset activity. (I) Local membrane dynamics during combined cooling and tripolar KHFS at 20 kHz. (J) Enlarged onset phase during bipolar KHFS (10 kHz). (K) Enlarged steady-state phase during bipolar KHFS, highlighting the spatial separation between stimulus-driven activity and the blocked region. (L) Multimodal block during combined cooling and 10-kHz bipolar KHFS. (M) Combined cooling and tripolar KHFS at 20 kHz suppresses onset and establishes directional valve behavior. The 20-kHz panels test whether the cooling-assisted response generalizes beyond the primary 10-kHz condition.

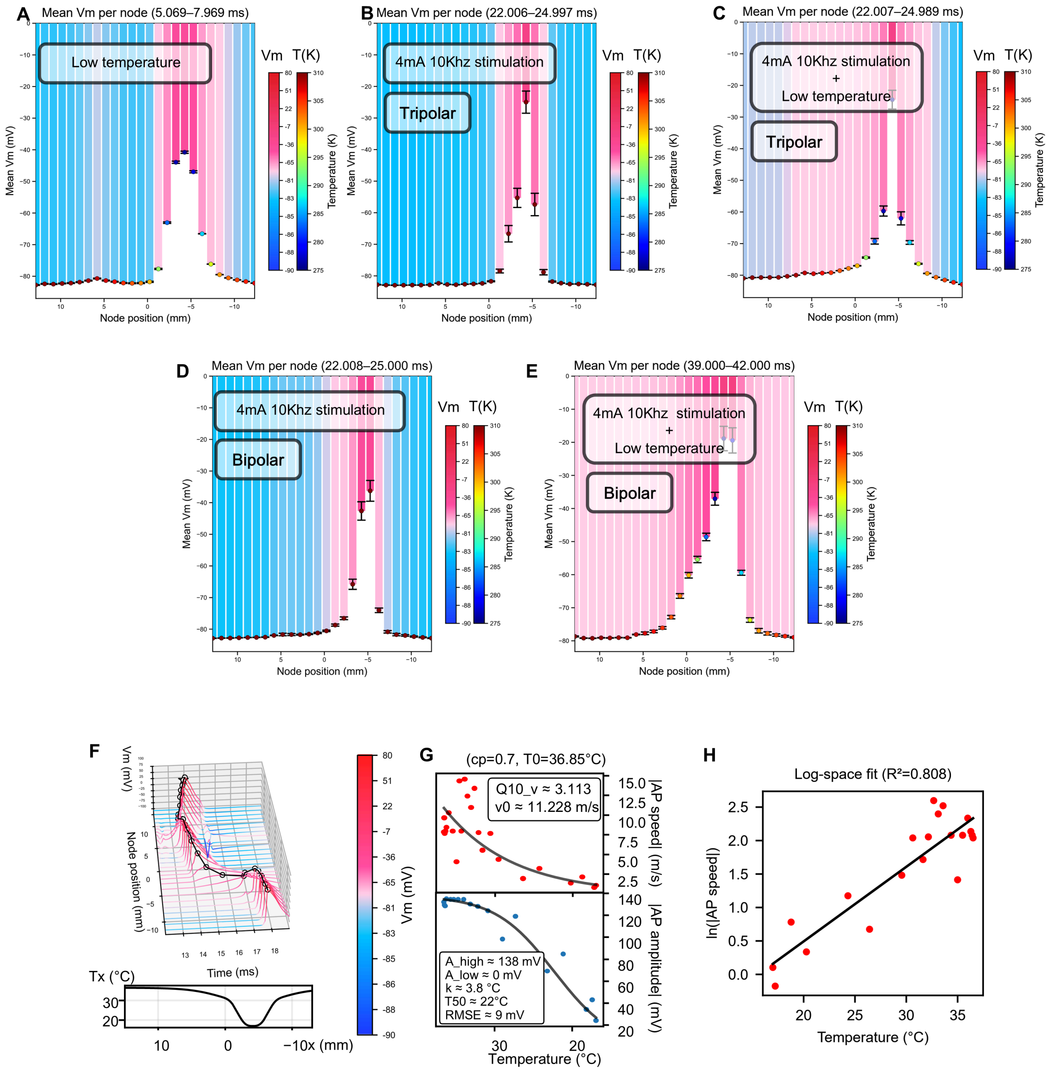


Fig. S13. Node-resolved mean membrane potential quantifies subthreshold states during cooling and KHFS block (A) Mean nodal membrane potential ($V_m$) during cooling alone (5.069 to 7.969

ms), with the temperature field overlaid to localize the cooling-induced subthreshold depolarization zone. (B) Mean nodal $V_m$ during tripolar KHFS (10 kHz, 4 mA; 22.006 to 24.997 ms). (C) Mean nodal $V_m$ during combined cooling and tripolar KHFS (22.003 to 25.000 ms), showing a stabilized subthreshold state. (D) Mean nodal $V_m$ during bipolar KHFS (10 kHz, 4 mA; 22.008 to 25.000 ms). (E) Mean nodal $V_m$ during combined cooling and bipolar KHFS (39 to 42 ms). In (A) to (E), error bars indicate SD within the stated time window. (F) Action-potential waveform under mild cooling (minimum temperature, approximately 15 °C), showing preserved but slowed conduction. (G) Action-potential amplitude and conduction velocity across temperature. (H) Log-linear relation between conduction velocity and temperature. Related to Fig. 3.

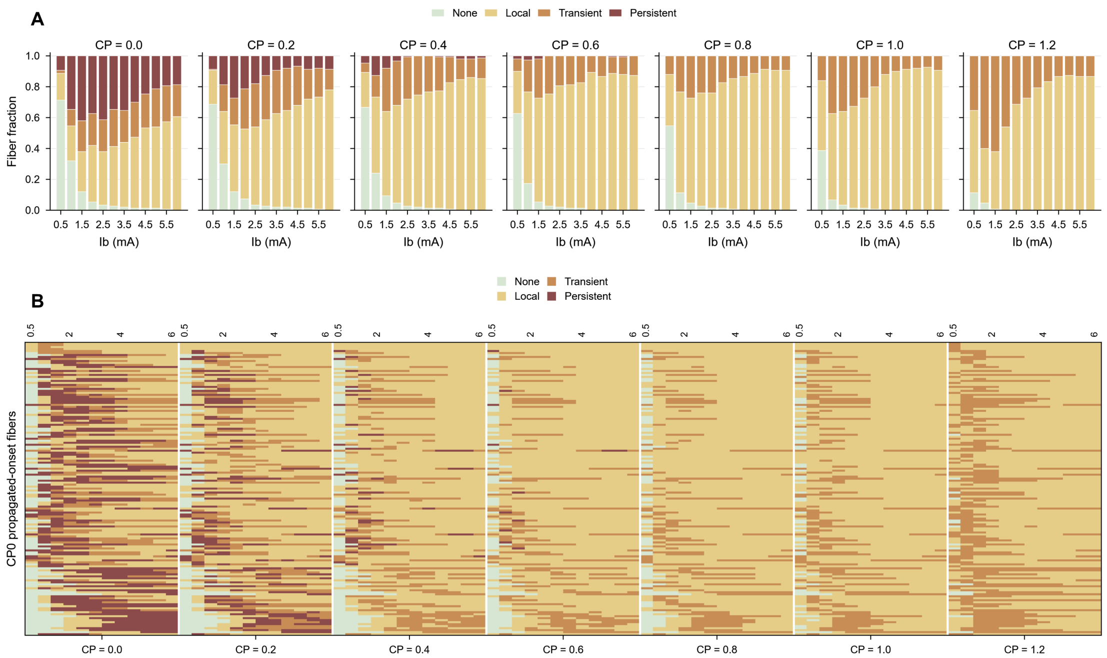


Fig. S14. Cooling shifts population-level KHFS onset responses from propagated to nonpropagating states (A) Fractions of fibers classified as none, local/nonpropagating, transient propagated onset, or persistent propagated onset across KHFS amplitudes and cooling-profile parameters (CP = 0 to 1.2). (B) Fiber-by-amplitude heat maps showing the corresponding onset class for each modeled fiber. Increasing cooling reduces persistent propagated onset over a broad range of KHFS amplitudes. Related to Fig. 3.

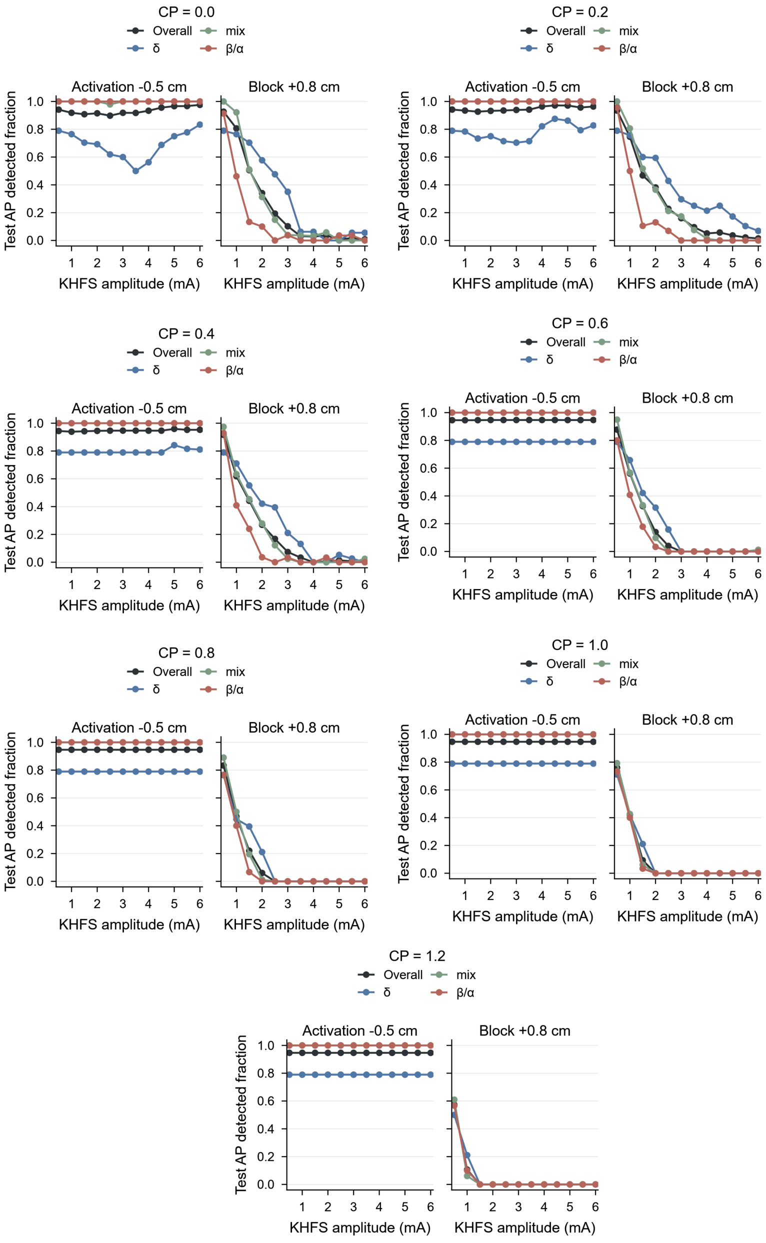


Fig. S15. Cooling- and fiber-dependent recruitment before and after the KHFS block site For cooling-profile parameters CP = 0 to 1.2, paired plots show the fraction of detected action potentials near the activation site (−0.5 cm) and downstream of the block site (+0.8 cm) as a function of KHFS amplitude. Curves report the overall population and Aδ, mixed-diameter, and Aα/Aβ fiber classes. Recruitment remains high near the activation site, whereas downstream detection falls with KHFS amplitude and is shifted by cooling strength and fiber class. Related to Fig. 3.

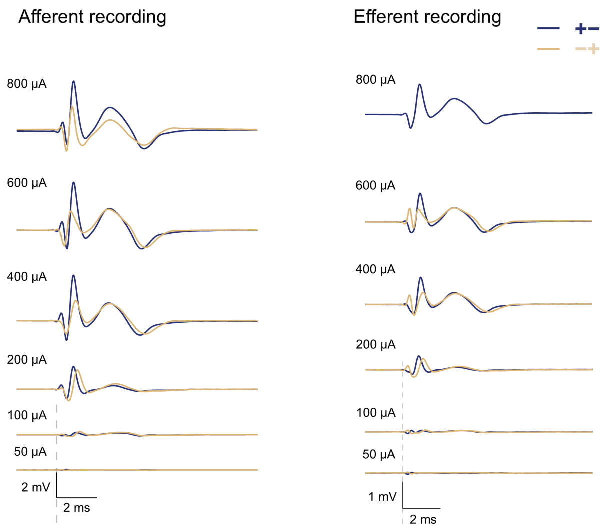


Fig. S16. CNAP waveforms across stimulus amplitudes and polarities in afferent and efferent recordings Representative afferent and efferent CNAPs evoked by 50- to 800-µA bipolar stimulation are shown for the two opposite stimulus polarities. Labeled N1–P1, N2–P2, and P3 components illustrate the amplitude-dependent emergence and polarity robustness of the fast and slow compound responses. Scale bars, 2 ms; 2 mV for afferent recordings and 1 mV for efferent recordings. Related to Fig. 4.

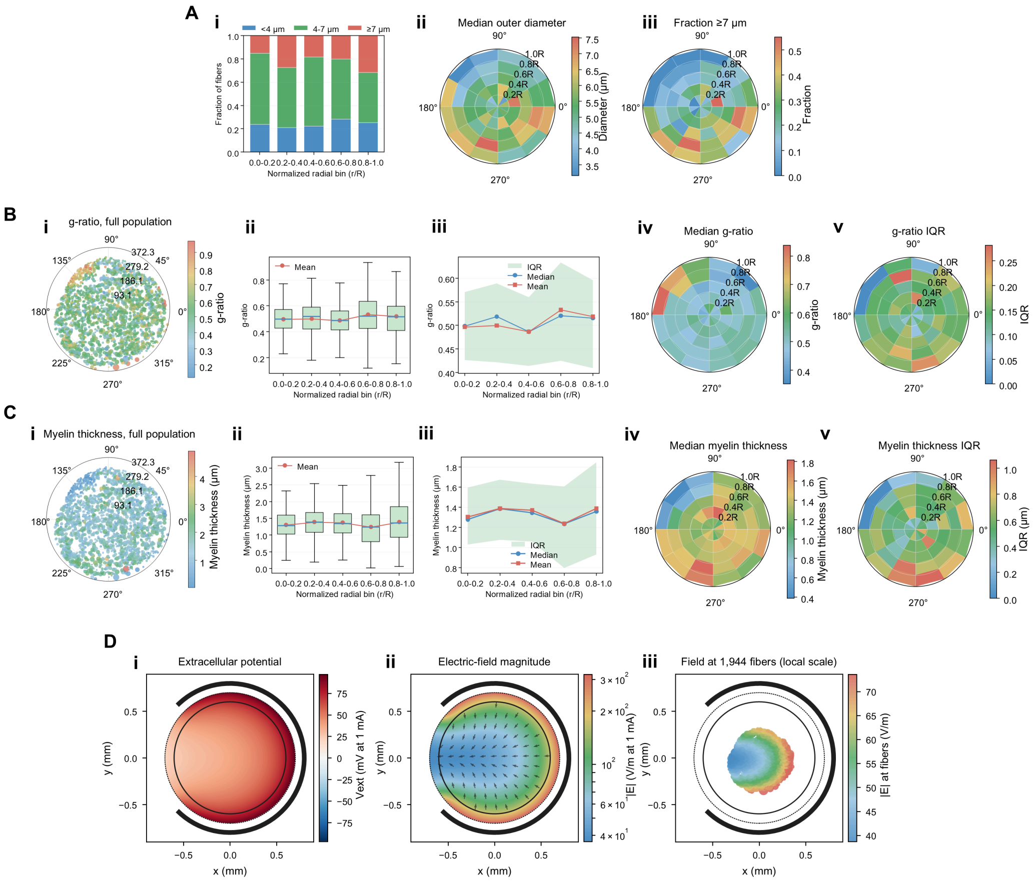

Fig. S17. Spatial distributions of fiber geometry and the ring-electrode field in the reconstructed sciatic nerve (A) Spatial and radial distributions of outer fiber diameter, including the fraction of fibers in the <4-μm, 4- to <7-μm, and ≥7-μm classes. (B) Spatial and radial distributions of g-ratio. (C) Spatial and radial distributions of myelin thickness. (D) Extracellular potential, electric-field magnitude, and fiber-position sampling beneath the ring electrode for the reconstructed 1,944-fiber population. Related to Fig. 4.

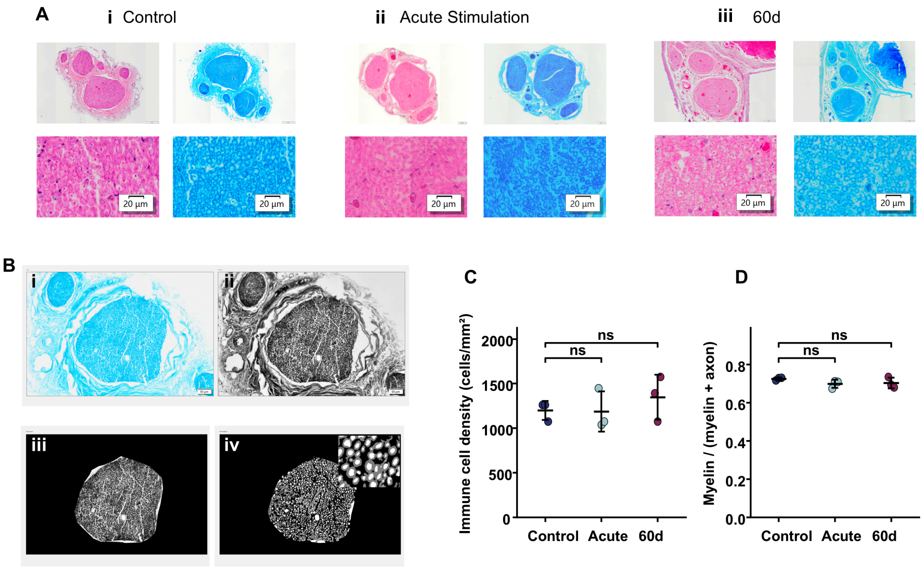


Fig. S18. Histological and morphometric assessment of sciatic nerves after acute cooling-assisted operation and chronic implantation (A) Hematoxylin and eosin (H&E; red) and Luxol fast blue

(blue) staining of sciatic nerves from untreated controls, animals examined after acute stimulation, and animals examined after 60 days of implantation. (i to iii) Representative low- and high-magnification views. (B) Representative image-processing workflow for myelin morphometry (i to iv). (C) Immune-cell density in H&E-stained sections showed a modest upward trend after acute stimulation and after 60 days of implantation. (D) Myelin area fraction, defined as myelin/(myelin + axon), remained similar among groups. Data are mean ± SD; n=3 animals per group. Overall group differences were evaluated using a two-sided Kruskal–Wallis test, followed by two-sided Mann–Whitney U tests with Holm correction for the displayed pairwise comparisons. All displayed comparisons were not significant after correction (adjusted $P > 0.05$); ns, not significant.

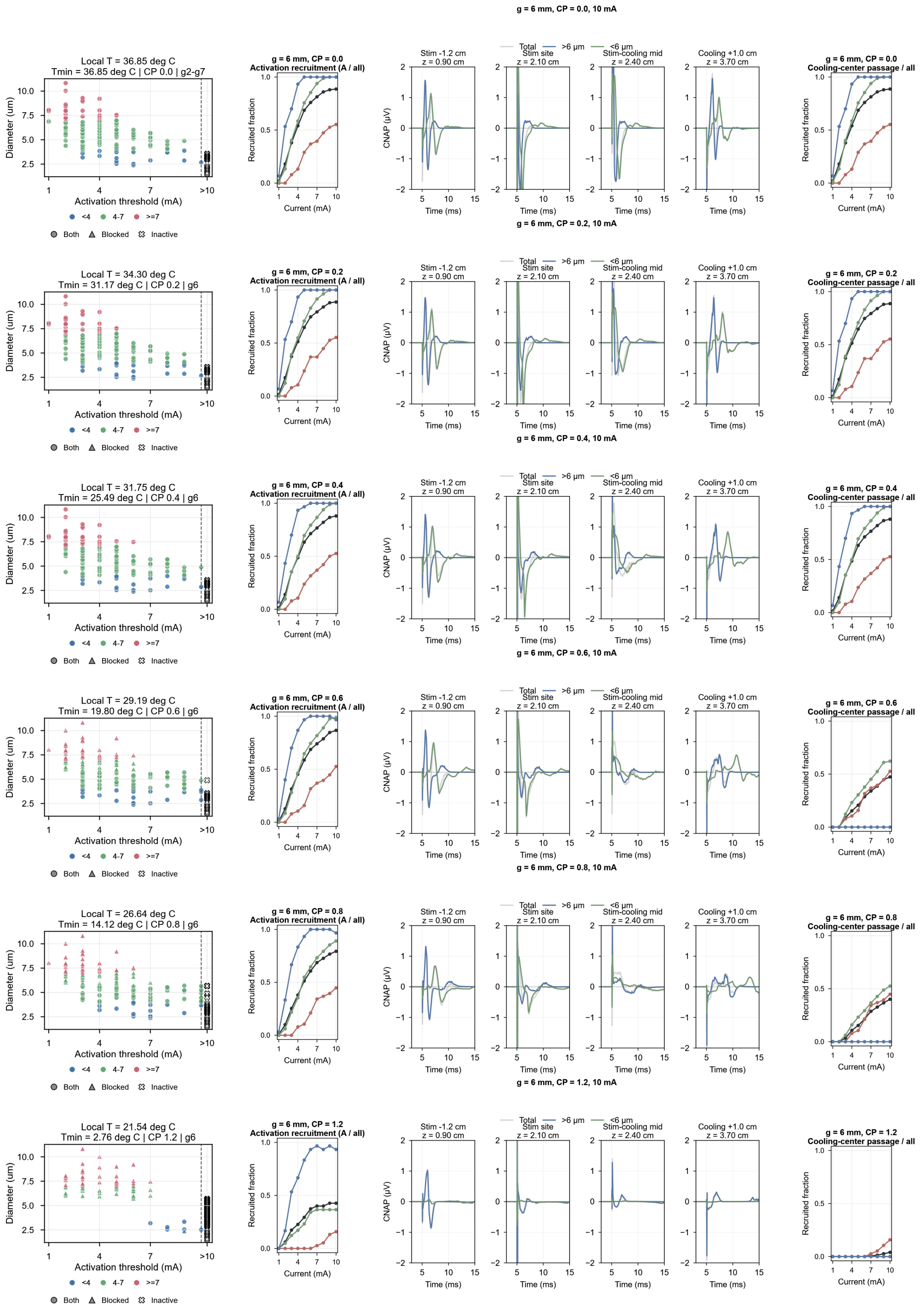

Fig. S19. Cooling-profile strength reshapes activation, propagation, and CNAP composition at a 6-mm block–drive spacing Rows show cooling-profile parameters (CP) of 0.0, 0.2, 0.4, 0.6, 0.8, and 1.2 at a block–drive spacing of 6 mm; the corresponding local and minimum temperatures are annotated. Within each row, plots show, from left to right, fiber diameter versus activation threshold and propagation outcome; activation recruitment versus stimulation current; simulated CNAPs at 10 mA at four positions along the stimulation-to-cooling axis, decomposed into total, >6-μm, and <6-μm contributions; and the fraction of the population passing the cooling center versus current. The four CNAP positions are 1.2 cm proximal to the stimulation site (z = 0.90 cm), the stimulation site (z = 2.10 cm), the stimulation–cooling midpoint (z = 2.40 cm), and 1.0 cm distal to the cooling center (z = 3.70 cm). "Both" denotes fibers for which local activation and passage through the cooled region were both detected, "Blocked" denotes fibers for which local activation was detected but passage was not, and "Inactive" denotes fibers for which recruitment was not detected within the tested current range.

### Supplementary Tables

Tables S1 to S10 are provided below in editable form. Table titles and legends precede each table; units are reported in the column headings or in a dedicated Unit column.

**Table S1. Key resources**

Materials, equipment, software, and experimental resources used in this study.

Resources

| Resource | Source / supplier | Identifier / version | Category |
|---|---|---|---|
| PDMS prepolymer and curing agent | Dow Sylgard 184 | Cat# 01064291Z | Chemicals, peptides, and recombinant proteins |
| Titanium foil (10 µm) | Sigma-Aldrich | Cat# GF25842933 | Chemicals, peptides, and recombinant proteins |
| Platinum foil (40 µm) | Goodfellow | Platinum foil, 0.04 mm, custom order | Chemicals, peptides, and recombinant proteins |
| Conductive silver epoxy | MG Chemicals | Cat# 8331S | Chemicals, peptides, and recombinant proteins |
| Diamond microparticles, mean size 2.5 µm | Pureon (Microdiamant) | Microdiamant MSY 2.5 | Chemicals, peptides, and recombinant proteins |
| Biocompatible silicone adhesive | NuSil | Cat# MED-2000 | Chemicals, peptides, and recombinant proteins |
| Water-soluble tape | Aquasol Corporation | Cat# ASWT-1 | Chemicals, peptides, and recombinant proteins |
| Paper tape | 3M | 3M Masking Tape 2308 | Chemicals, peptides, and recombinant proteins |
| Silicone tubing (0.7 mm ID / 1.3 mm OD) | VWR Collection | Cat# 228-1450 | Chemicals, peptides, and recombinant proteins |
| Transfer connector / adapter | IDEX Health & Science | Cat# P-850 / P-851 | Chemicals, peptides, and recombinant proteins |
| Ethanol | Sigma-Aldrich | Cat# 100983 | Chemicals, peptides, and recombinant proteins |
| Xylene or xylene substitute | Sigma-Aldrich | Cat# XX0060-4L | Chemicals, peptides, and recombinant proteins |
| Deionized / distilled water | Sigma-Aldrich | Cat# EM3234 | Chemicals, peptides, and recombinant proteins |
| Agar | Sigma-Aldrich | Cat# 05040 | Chemicals, peptides, and recombinant proteins |
| Phosphate-buffered saline (PBS) | Gibco | Cat# 10010031 | Chemicals, peptides, and recombinant proteins |
| Sterile normal saline | Baxter | Item# 2B7231 | Chemicals, peptides, and recombinant proteins |
| Erythromycin ophthalmic ointment | Bausch & Lomb | NDC 24208-910-55 | Chemicals, peptides, and recombinant proteins |
| Isoflurane | Baxter | NDC 10019-360-40 | Chemicals, peptides, and recombinant proteins |
| Paraffin embedding wax | Leica Biosystems | Cat# 39601006 | Chemicals, peptides, and recombinant proteins |
| Hematoxylin solution | Sigma-Aldrich | Cat# HHS32 | Chemicals, peptides, and recombinant proteins |

| Resource | Source / supplier | Identifier / version | Category |
|---|---|---|---|
| Eosin solution | Sigma-Aldrich | Cat# HT110116 | Chemicals, peptides, and recombinant proteins |
| Differentiation solution / acid alcohol | Sigma-Aldrich | Cat# A3179 | Chemicals, peptides, and recombinant proteins |
| Bluing reagent / ammonia water | Sigma-Aldrich | Cat# S5134 | Chemicals, peptides, and recombinant proteins |
| Luxol fast blue staining solution | Sigma-Aldrich | Cat# L0294 | Chemicals, peptides, and recombinant proteins |
| lithium carbonate solid | Sigma-Aldrich | Cat# 255823 | Chemicals, peptides, and recombinant proteins |
| Mounting medium / neutral balsam | Cat# 317616 | Cat# 317616 | Chemicals, peptides, and recombinant proteins |
| Hematoxylin and eosin staining kit / solutions | Phygene | Cat# PH0516 | Critical commercial assays |
| Luxol fast blue myelin staining kit | Servicebio | Cat# G1030 | Critical commercial assays |
| Sprague–Dawley rat (adult male, 250–300 g) | SHENZHEN HUATENG BIOMEDICAL TECHNOLOGY CO,LTD | SD | Experimental models: Organisms/strains |
| MATLAB | MathWorks | 2021; https://www.mathworks.com/products/matlab.html | Software and algorithms |
| COMSOL Multiphysics | COMSOL | https://www.comsol.com/comsol-multiphysics | Software and algorithms |
| NEURON | Yale / NEURON Project | 8.26; https://neuron.yale.edu/neuron/ | Software and algorithms |
| Python | Python Software Foundation | https://www.python.org/ | Software and algorithms |
| 3.3 kΩ chip thermistor, 0402 package | Murata Manufacturing Co., Ltd. | Cat# NCP15XW332J03RC | Other |
| K-type fine-wire thermocouple | Cape Vincent | TYPEC K 2*0.1mm | Other |
| Thermocouple recorder | SMRF-T | CT06A | Other |
| Laboratory-built thermistor detection unit | This paper | N/A | Other |
| Electrophysiology recording system | jiansu brain medical technology Co.ltd,nanjing china | Neurologo amplifier | Other |
| Electrical stimulator | PlexStim | Electrical Stimulator 2.0 | Other |
| Arduino development board | Arduino | Arduino Uno | Other |
| Absorbable surgical suture, polyglycolic acid (PGA), USP 4-0 (1.5 metric), 90 cm, 1/2 circle, round bodied, 19 mm needle | Yankee | N/A | Other |
| Paraffin embedding cassette | our laboratory 3d print | N/A | Other |
| Embedding mold | our laboratory 3d print | N/A | Other |
| Disposable microtome blade | LIICA | Cat# 14035838382CN | Other |
| Microscope slide (positively charged if used) | CITOTEST | REF.188108 | Other |
| Coverslip | CITOTEST | REF.10212432C | Other |
| LPKF S4 | Electrode fabrication | Shared fabrication platform; described directly in Methods | Shared instrument (Methods) |
| Phrozen Mini 8K | Microchannel fabrication | Shared 3D-printing platform; described directly in Methods | Shared instrument (Methods) |

| Resource | Source / supplier | Identifier / version | Category |
| --- | --- | --- | --- |
| Leica paraffin microtome | Histology, H&E staining, Luxol fast blue staining, and imaging | Routine shared histology instrument | Shared instrument (Methods) |
| Olympus FV3000 | Histology, H&E staining, Luxol fast blue staining, and imaging | Routine shared microscopy platform | Shared instrument (Methods) |
| VS200-BU slide scanner | Histology, H&E staining, Luxol fast blue staining, and imaging | Routine shared slide-scanning platform | Shared instrument (Methods) |
| DScope U3P100 | Electrochemical characterization | Routine shared oscilloscope | Shared instrument (Methods) |
| Osmo Action 3 | Assessment of cooling effects on onset response | Routine shared video-recording device | Shared instrument (Methods) |

**Table S2. Device, nerve-geometry, and stimulation parameters**

Parameters used to define the device-level and digital-twin geometry.

Main nerve and stimulation parameters

| Parameter | Value |
|---|---|
| Modeled nerve length | 5.1 cm |
| Intraneural diameter | 1.2 mm |
| Intraneural radius | 600 µm |
| Epineurium thickness | 100 µm |
| Extracellular conductivity | 1.0 S/m |
| Stimulation geometry | bipolar point-source pair |
| Stimulation center | 2.0 cm |
| Contact radial position | 800 µm |
| Inter-contact spacing | 1.0 mm |
| Pulse waveform | symmetric biphasic |
| Phase width | 0.05 ms |
| Interphase gap | 0 ms |
| Stimulus onset | 10 ms |
| Mixed-population current sweep | 0.5–6.0 mA in 0.5-mA increments |
| Representative mixed-population point | 6.0 mA; Fig. 2N |
| Neural finite-element panel condition | 4 mA; figs. S4 and S13 |
| In vivo low-frequency CNAP default | 400 µA, 50 µs per phase |
| In vivo CNAP amplitude series | 50–800 µA; fig. S16 |
| KHFS frequency conditions | 10 and 20 kHz, panel-specific; 20 kHz tests frequency generalization |

Block–drive geometry conditions

| Context | Block–drive spacing | Use |
|---|---|---|
| Final in vivo device | 9 mm | Experimental NeuroSwitch cuff |
| Representative population model | 8 mm | 150-fiber model; stimulation and cooling centers at 2.0 and 2.8 cm |
| Fig. 5P/S19 scan | 6 mm | Position-resolved spacing-scan condition |

**Table S3. Thermal properties and cooling-field parameters**

Material thermal properties and the representative cooling-field condition used in simulations.

Material thermal properties

| Material / subdomain | Thermal conductivity λ (W $m^{-1}$ $K^{-1}$) | Density ρ (kg $m^{-3}$) | Specific heat capacity $C_p$ (J $kg^{-1}$ $K^{-1}$) | Notes | Ref |
|---|---|---|---|---|---|
| Muscle | 0.49 | 1090 | 3421 | Solid; isotropic. | (*69*) |
| Epidural tissue | 0.45 | 1040 | 3500 | Solid; isotropic. Approximated using generic connective-tissue thermal properties from IT’IS database. | (*69*) |
| Perineurium | 0.30 | 1100 | 3400 | Solid; isotropic. Perineurium-specific thermophysical data are scarce; approximated using generic connective-tissue thermal properties from IT’IS database. | (*69*) |
| Tracts | 0.55 | 1040 | 3600 | Solid; isotropic. Tracts treated as CNS white-matter-like tissue; values rounded from literature/IT’IS database. | (*69*) |
| Nerve fiber | 0.21 | 911 | 2348 | Solid; isotropic. Current values match adipose/epidural fat; replace with peripheral nerve thermophysical properties if this row represents fascicle/axoplasm. | (*69*) |
| Myelin | 0.45 | 911 | 3500 | Solid; isotropic. Myelin row appears to mix adipose-like density with water-like conductivity/heat capacity; verify intended proxy (lipid-rich myelin vs white matter). | (*69*) |
| PDMS (polydimethylsiloxane, 0% diamond) | 0.180 | 970 | 1620 | Solid; isotropic. Baseline composite: 0 wt% diamond. | |
| Diamond-doped PDMS (10%) | 0.205 | 1046 | 1510 | Solid; isotropic. Diamond fraction: 10 wt% (case-dependent). Diamond doping ratio is defined as the mass fraction of diamond filler in the PDMS matrix (wt%). | |
| Diamond-doped PDMS (20%) | 0.239 | 1134 | 1370 | Solid; isotropic. Diamond fraction: 20 wt% (case-dependent). Diamond doping ratio is defined as the mass fraction of diamond filler in the PDMS matrix (wt%). | |

| Material / subdomain | Thermal conductivity λ (W $m^{-1}$ $K^{-1}$) | Density ρ (kg $m^{-3}$) | Specific heat capacity $C_p$ (J $kg^{-1}$ $K^{-1}$) | Notes | Ref |
|---|---|---|---|---|---|
| Diamond-doped PDMS (30%) | 0.287 | 1239 | 1330 | Solid; isotropic. Diamond fraction: 30 wt% (case-dependent). Diamond doping ratio is defined as the mass fraction of diamond filler in the PDMS matrix (wt%). | |
| Diamond-doped PDMS (40%) | 0.349 | 1366 | 1384 | Solid; isotropic. Diamond fraction: 40 wt% (case-dependent). Diamond doping ratio is defined as the mass fraction of diamond filler in the PDMS matrix (wt%). | |
| Diamond-doped PDMS (50%) | 0.448 | 1521 | 1169 | Solid; isotropic. Diamond fraction: 50 wt% (case-dependent). Diamond doping ratio is defined as the mass fraction of diamond filler in the PDMS matrix (wt%). | |
| Ethanol, $C_2H_5OH$ (liquid, ~25 °C) | 0.169 | 785.1 | 2440 | Fluid. Constant room-temperature properties (≈25 °C). (Viscosity handled in the flow module.) | |
| Platinum, Pt (electrode conductor, ~25 °C) | 71.6 | 21450 | 133 | Solid; isotropic. Replaces previous Pt–Ir entry. | |

Representative cooling-field parameters

| Parameter | Value |
|---|---|
| Baseline temperature | 37 °C |
| Cooling center | 2.8 cm |
| CP | 1.0 |
| Width-scale factor | 2.0 |
| Representative steady-state mixed-population minimum temperature | 8.44 °C |
| Distal 35 °C recovery station | 4.706 cm |

### Table S4. Electrical properties

Electrical properties assigned to tissue, device, and electrode domains.

Material and domain electrical properties

| Material / subdomain | Electrical conductivity σ (S $m^{-1}$) | Relative permittivity $\varepsilon_r$ (–) | Notes | Ref |
|---|---|---|---|---|
| Ethanol, $C_2H_5OH$ (liquid) | 1e-6 | 24.3 | Isotropic (low-conductivity dielectric). | |
| PDMS (polydimethylsiloxane) | 9.86e-6 | 2.75 | Isotropic. | |
| PDMS (diamond-doped, 10–50%) | 9.86e-6 | 2.75 | Electrical properties assumed identical to baseline PDMS unless otherwise specified. | |
| Epidural tissue | 0.018 | 9.4e6 | Isotropic (as defined in the model). | (*70, 71*) |
| Muscle | 0.4 | 5e5 | Isotropic (as defined in the model). | (*69, 71*) |
| Perineurium | 2.7e-4 | 2000 | Isotropic. | (*57*) |
| Platinum, Pt (electrode conductor) | 9.43e6 | 1 | Isotropic; replaces previous Pt–Ir entry (σ updated to Pt). | (*57*) |
| Tracts (anisotropic) | 1.03, 0.08, 0.08 | 80, 30000, 30000 | Anisotropic (principal components) as defined in model. | (*70, 71*) |
| Nerve fiber (anisotropic σ) | 1.03, 0.08, 0.08 | 80 | Anisotropic σ; isotropic $\varepsilon_r$. | (*70, 71*) |
| Myelin | 1.3e-7 | 15, 1, 1 | $\varepsilon_r$ provided as directional components in model. | (*70, 72*) |
| Ground (ideal conductor, boundary) | 1e9 | 1 | Used to approximate electrical ground. | |

**Table S5. Myelinated-axon geometry and fiber sampling**

Geometric parameters and sampling rules for the reconstructed mixed-fiber population.

Model scope: the fixed node, MYSA, myelin, and 1000-µm internode geometry below belongs to the EFM neuron model used for fiber-specific Aβ and Aδ simulations. The sampling block describes the separate reconstructed population.

Myelinated-axon geometry

| **Parameter (English name)** | **Symbol** | **Value** | **Unit** | **Notes** | **Reference** |
|---|---|---|---|---|---|
| Node length | L_node | 1.0 | µm | Node of Ranvier | (*42*) |
| MYSA length | L_MYSA | 3.0 | µm | Paranode compartment | (*42*) |
| Myelinated segment length (internode) | L_myelin | 1000.0 | µm | Myelinated segment between nodes | (*42*) |
| Node diameter | D_node | 2.6 | µm | Node caliber set using PNS nodal constriction (30–50% of internodal axon diameter) with MRG-based geometry scaling. | (*42*); (*73, 74*) |
| MYSA diameter (proximal) | D_MYSA_prox | 5.4 | µm | Proximal MYSA diameter approximates internodal axon diameter; taper follows PNS nodal constriction. | (*42*); (*73, 74*) |
| MYSA diameter (distal) | D_MYSA_dist | 2.8 | µm | Distal MYSA diameter approaches nodal caliber; taper follows PNS nodal constriction. | (*42*); (*73, 74*) |
| Myelin diameter | D_myelin | 7.8 | µm | Diameter of the myelinated segment | (*42*) |
| Submyelin gap thickness (periaxonal space) | t_gap | 0.01 | µm | Periaxonal gap set to 10 nm, within reported ~10–15 nm range in compact myelin ultrastructure. | (*41*) |
| MYSA diameter taper (linear) | D_MYSA(x) | 5.4 → 2.8 | µm | Linear taper along MYSA: D_MYSA(x)=D_MYSA_prox+(D_MYSA_dist-D_MYSA_prox)*x/L_MYSA, x∈[0,L_MYSA]. | (*42*) |

Fiber sampling parameters

| **Parameter** | **Value** |
|---|---|
| Eligible source fibers | 1,944 |
| Simulated representative fibers | 150 |
| Sampling strategy | spatial-diameter stratified |
| Radial bins | 5 |
| Angular bins | 8 |
| Diameter quantile bins | 4 |
| Nonempty stratification cells | 154 |
| Sampling seed | 20260418 |
| Maximum included diameter | 16 µm |
| Myelin filter | positive myelin thickness required |

**Table S6. Fiber classes and model-family composition**

Fiber classes and model-family assignments used in the reconstructed population.

Model scope: compartment model only.

Fiber classes in the reconstructed population

| Fiber class | Diameter rule | Count |
|---|---|---|
| Aδ | <4 μm | 38 |
| Mixed-diameter | 4 to <7 μm | 82 |
| Aα/β | ≥7 μm | 30 |

Model-family composition

| Model family and class | Count |
|---|---|
| Peña-style Aδ (*57*) | 38 |
| Peña-style mixed-diameter (*57*) | 52 |
| MRG-style mixed-diameter | 30 |
| MRG-style Aα/β | 30 |

### Table S7. Shared temperature-scaling and potassium-current parameters

Common temperature-scaling constants and iKf/iKs parameters.

Model scope: compartment model only.

Shared temperature-scaling parameters

| Parameter | Description | Value | Unit | Applies to | Notes |
|---|---|---|---|---|---|
| E_K | Baseline K reversal potential | -90 | mV | K2P, iKf_iKs | used to compute E_KT via TkEk \| Value shown as -0.09 V in model; used to form E_KT and deltaV. |
| F | Faraday constant | 96485 | C/mol | K2P, iNa16_iNa11 | used if E_NaT uses Nernst form \| Used in V_T_mV = R*T/F. |
| R | Gas constant | 8.314 | J/(mol*K) | K2P, iNa16_iNa11 | used if E_NaT uses Nernst form \| Used in V_T_mV = R*T/F. |
| T_ref | Reference temperature for scaling terms | 310 | K | K2P, iKf_iKs, iNa16_iNa11 | reference point for TkVm and TkEk \| reference point \| From model parameter table; used in TkVm and TkEk. |
| V_rest | Resting voltage (used in TkVm normalization) | -80 | mV | K2P, iKf_iKs, iNa16_iNa11 | enters TkVm scaling \| enters TkVm \| Value shown as -0.08 V in model. |

iKf/iKs parameters

| Parameter | Meaning | Value | Unit | Applies to / temperature role | Reference |
|---|---|---|---|---|---|
| gKf_node | Fast K current maximum conductance (node region) | 0 S $cm^{-2}$ | S $cm^{-2}$ | iKf: sets regional current amplitude | (*42*) |
| gKf_MYSA | Fast K current maximum conductance (MYSA region) | 0.15074 S $cm^{-2}$ | S $cm^{-2}$ | iKf: sets regional current amplitude | (*42*) |
| gKf_STIN_FLUT | Fast K current maximum conductance (STIN/FLUT regions) | 0.002568 S $cm^{-2}$ | S $cm^{-2}$ | iKf: sets regional current amplitude | (*42*) |
| gKs_node | Slow K current maximum conductance (node region) | 0 S $cm^{-2}$ | S $cm^{-2}$ | iKs: sets regional current amplitude | (*75*) |
| gKs_MYSA | Slow K current maximum conductance (MYSA region) | 0.002581 S $cm^{-2}$ | S $cm^{-2}$ | iKs: sets regional current amplitude | (*42*) |
| gKs_STIN_FLUT | Slow K current maximum conductance (STIN/FLUT regions) | 0.002581 S $cm^{-2}$ | S $cm^{-2}$ | iKs: sets regional current amplitude | (*42*) |
| k_TkVm | Vm temperature coefficient in TkVm | 0.45 | mV $K^{-1}$ | both: sets temperature dependence of TkVm | (*33*) |
| k_TkEk | E_K temperature coefficient in TkEk | 0.31 | mV $K^{-1}$ | both: sets temperature dependence of TkEk | (*33*) |
| Q10_K | Q10 base for Kf/Ks kinetics | 3 | — | kf, ks: kinetic acceleration base | (*42*) |

| Parameter | Meaning | Value | Unit | Applies to / temperature role | Reference |
|---|---|---|---|---|---|
| Tref_kf | Reference temperature for Q10_kf | 307.15 | K | kf: reference point in Q10_kf(T) | (*42*) |
| Tref_ks | Reference temperature for Q10_ks | 309.15 | K | ks: reference point in Q10_ks(T) | (*42*) |
| q10_slope | Exponent slope in Q10 expressions | 0.1 | $K^{-1}$ | kf, ks: sets temperature sensitivity per 10 K | (*42*) |
| anA | alpha_Kf scale coefficient | 0.0462e6[1/(V*s)] | 1/(V·s) | kf: kinetic shape parameter | (*42*) |
| anB | alpha_Kf voltage-shift coefficient | -83.2e-3[V] | V | kf: kinetic shape parameter | (*42*) |
| anC | alpha_Kf slope coefficient | 1.1e-3[V] | V | kf: kinetic shape parameter | (*42*) |
| bnA | beta_Kf scale coefficient | 0.0824e6[1/(V*s)] | 1/(V·s) | kf: kinetic shape parameter | (*42*) |
| bnB | beta_Kf voltage-shift coefficient | -66e-3[V] | V | kf: kinetic shape parameter | (*42*) |
| bnC | beta_Kf slope coefficient | 10.5e-3[V] | V | kf: kinetic shape parameter | (*42*) |
| asA | alpha_Ks scale coefficient | $0.3 \times 1000\ s^{-1}$ | $s^{-1}$ | ks: kinetic shape parameter | (*42*) |
| asB | alpha_Ks voltage-shift coefficient | -0.027 [V] | V | ks: kinetic shape parameter | (*42*) |
| asC | alpha_Ks slope coefficient | -0.005[V] | V | ks: kinetic shape parameter | (*42*) |
| bsA | beta_Ks scale coefficient | $0.03 \times 1000\ s^{-1}$ | $s^{-1}$ | ks: kinetic shape parameter | (*42*) |
| bsB | beta_Ks voltage-shift coefficient | 0.010[V] | V | ks: kinetic shape parameter | (*42*) |
| bsC | beta_Ks slope coefficient | -0.001[V] | V | ks: kinetic shape parameter | (*42*) |
| vtraub | Voltage offset used in K channel kinetics | -80[mV] | mV | both: sets effective voltage reference in kinetics | (*42*) |
| kf(0) | Initial value of kf gating variable | 0 | — | kf: initialization | (*42*) |
| ks(0) | Initial value of ks gating variable | 0 | — | ks: initialization | (*42*) |

**Table S8. Sodium-current and vtrap parameters**

$i_{\mathrm{Na16}}$, $i_{\mathrm{Na11}}$, and temperature-responsive $i_{\mathrm{Na18}}$ conductance, temperature, and voltage-trap kinetic parameters. The $\mathrm{Na_V}1.8$ parameter block includes the shared temperature-mapping coefficients, $\mathrm{Na_V}1.8$ gate-specific $Q_{10}$ values, empirical cold-resistance scaling anchors, and low-temperature activation-shift coefficient used in Eqs. 60 to 67.

The iNa16/iNa11 values in this table belong to the EFM neuron model, not to the compartmental active-node model summarized in Table S9.

Model scope: EFM neuron model.

iNa16/iNa11 parameters

| Parameter | Meaning | Value | Unit | Applies to | Temperature role | Model family | Compartment | Software module |
|---|---|---|---|---|---|---|---|---|
| gNa16 | Max conductance (fast Na component in iNa16) | 16000 | mS cm⁻² (= 16 S cm⁻²) | iNa16 | scales current amplitude | EFM neuron model | Whole modeled neuron/fiber domain; not compartment-specific | COMSOL Multiphysics |
| gNa11 | Max conductance (slow/persistent Na component in iNa11) | 0.5 | S cm⁻² | iNa11 | scales current amplitude | EFM neuron model | Whole modeled neuron/fiber domain; not compartment-specific | COMSOL Multiphysics |
| E_Na | Baseline Na reversal potential | 60 | mV | both | used to compute E_NaT | EFM neuron model | Whole modeled neuron/fiber domain; not compartment-specific | COMSOL Multiphysics |
| k_TkVm | Vm temperature coefficient | 0.45 | mV K⁻¹ | both | used in TkVm | EFM neuron model | Whole modeled neuron/fiber domain; not compartment-specific | COMSOL Multiphysics |
| k_TkENa | E_Na temperature coefficient | 0.2 | mV K⁻¹ | both | used in TkENa | EFM neuron model | Whole modeled neuron/fiber domain; not compartment-specific | COMSOL Multiphysics |
| k_Vshift_m | Activation gate voltage-shift coefficient | 0.000353 | V/K | m, mp | used in VmNa_m | EFM neuron model | Whole modeled neuron/fiber domain; not compartment-specific | COMSOL Multiphysics |
| k_Vshift_h | Inactivation gate voltage-shift coefficient | 0.735 | mV K⁻¹ | h | used in VmNa_h | EFM neuron model | Whole modeled neuron/fiber domain; not compartment-specific | COMSOL Multiphysics |
| Q10_m_mp | Q10 base for m/mp kinetics | 2.2 | — | m, mp | acceleration factor | EFM neuron model | Whole modeled neuron/fiber domain; not compartment-specific | COMSOL Multiphysics |
| Q10_h | Q10 base for h kinetics | 2.9 | — | h | acceleration factor | EFM neuron model | Whole modeled neuron/fiber domain; not compartment-specific | COMSOL Multiphysics |
| amA, amB, amC | vtrap6 parameters | See the vtrap parameter block below in Table S8. | (as defined) | m | kinetic shape | EFM neuron model | Whole modeled neuron/fiber domain; not compartment-specific | COMSOL Multiphysics |
| bmA, bmB, bmC | vtrap7 parameters | See the vtrap parameter block below in Table S8. | (as defined) | m | kinetic shape | EFM neuron model | Whole modeled neuron/fiber domain; not compartment-specific | COMSOL Multiphysics |

| Parameter | Meaning | Value | Unit | Applies to | Temperature role | Model family | Compartment | Software module |
|---|---|---|---|---|---|---|---|---|
| ahA, ahB, ahC | vtrap8 parameters | See the vtrap parameter block below in Table S8. | (as defined) | h | kinetic shape | EFM neuron model | Whole modeled neuron/fiber domain; not compartment-specific | COMSOL Multiphysics |
| bhA, bhB, bhC | vtrap9 parameters | See the vtrap parameter block below in Table S8. | (as defined) | h | kinetic shape | EFM neuron model | Whole modeled neuron/fiber domain; not compartment-specific | COMSOL Multiphysics |
| ampA, ampB, ampC | vtrap1 parameters | See the vtrap parameter block below in Table S8. | (as defined) | mp | kinetic shape | EFM neuron model | Whole modeled neuron/fiber domain; not compartment-specific | COMSOL Multiphysics |
| bmpA, bmpB, bmpC | vtrap2 parameters | See the vtrap parameter block below in Table S8. | (as defined) | mp | kinetic shape | EFM neuron model | Whole modeled neuron/fiber domain; not compartment-specific | COMSOL Multiphysics |
| m0 | Initial m | m_inf(V_rest*TkVm) | — | iNa16 | via TkVm | EFM neuron model | Whole modeled neuron/fiber domain; not compartment-specific | COMSOL Multiphysics |
| h0 | Initial h | h_inf(V_rest*TkVm) | — | iNa16 | via TkVm | EFM neuron model | Whole modeled neuron/fiber domain; not compartment-specific | COMSOL Multiphysics |
| mp0 | Initial mp | mp_inf(V_rest*TkVm) | — | iNa11 | via TkVm | EFM neuron model | Whole modeled neuron/fiber domain; not compartment-specific | COMSOL Multiphysics |

iNa16/iNa11 vtrap parameters

| Parameter | vtrap function | Value | Unit | Applies to | Reference |
|---|---|---|---|---|---|
| amA | vtrap6 | 1860000.0 | 1/(V*s) | m | (*42*) |
| amC | vtrap6 | 0.0103 | V | m | (*42*) |
| bmB | vtrap7 | 0.0257 | V | m | (*42*) |
| amB | vtrap6 | 0.0214 | V | m | (*42*) |
| bmA | vtrap7 | 86000.0 | 1/(V*s) | m | (*42*) |
| bmC | vtrap7 | 0.00916 | V | m | (*42*) |
| ahB | vtrap8 | 0.114 | V | h | (*42*) |
| bhA | vtrap9 | 2300.0 | $s^{-1}$ | h | (*42*) |
| ahA | vtrap8 | 62000.0 | 1/(V*s) | h | (*42*) |
| bhC | vtrap9 | 0.0134 | V | h | (*42*) |
| ahC | vtrap8 | 0.011 | V | h | (*42*) |
| bhB | vtrap9 | 0.0318 | V | h | (*42*) |
| ampA | vtrap1 | 10000.0 | 1/(m*V) | mp | (*42*) |
| bmpC | vtrap2 | 0.01 | V | mp | (*42*) |
| ampC | vtrap1 | 0.0102 | V | mp | (*42*) |
| bmpA | vtrap2 | 250.0 | 1/(m*V) | mp | (*42*) |
| ampB | vtrap1 | 0.027 | V | mp | (*42*) |

| Parameter | vtrap function | Value | Unit | Applies to | Reference |
|---|---|---|---|---|---|
| bmpB | vtrap2 | 0.034 | V | mp | (*42*) |

<u>Na$_V$1.8 improved temperature-responsive parameters</u>

| Parameter | Meaning | Value | Unit | Applies to |
|---|---|---|---|---|
| $T_{\mathrm{ref,gate}}$ | Reference temperature for gate Q10 scaling | 295.15 | K | iNa18 |
| $T_{\mathrm{ref,amp}}$ | Reference temperature for conductance normalization | 303.15 | K | iNa18 |
| $T_{\mathrm{ref,vshift}}$ | Reference temperature for cold activation shift | 303.15 | K | iNa18 |
| $q_{10,m}$ | Activation-gate Q10 | 2.2 | dimensionless | iNa18 |
| $q_{10,h}$ | Fast-inactivation-gate Q10 | 2.8 | dimensionless | iNa18 |
| $q_{10,s}$ | Slow-gate s Q10 | 1 | dimensionless | iNa18 |
| $q_{10,u}$ | Slow-gate u Q10 | 1 | dimensionless | iNa18 |
| $k_{\mathrm{TkVm}}$ | Effective membrane-voltage temperature coefficient | 0.45 | mV K$^{-1}$ | iNa18 |
| $k_{\mathrm{TkENa}}$ | Effective Na reversal-potential temperature coefficient | 0.2 | mV K$^{-1}$ | iNa18 |
| $k_{\mathrm{Vshift},m}^{\mathrm{cold}}$ | Low-temperature activation-shift coefficient | 0.15 | mV K$^{-1}$ | iNa18 |
| $Q_{10,g,\mathrm{low}}$ | Low-temperature amplitude extrapolation factor | 1.25 | dimensionless | iNa18 |
| $Q_{10,g,\mathrm{high}}$ | High-temperature amplitude extrapolation factor | 1.2 | dimensionless | iNa18 |
| $g_{\mathrm{scale,Na18}}$ bounds | Normalized amplitude-scaling bounds | 0.15 to 2.50 | dimensionless | iNa18 |
| $V_{\mathrm{drv}}$ | Saturating driving-force scale | 200 | mV | iNa18 |
| $g_{\mathrm{raw,Na18}}$ anchor at 283.15 K | Temperature of empirical relative-current anchor (relative amplitude 0.62) | 283.15 | K | iNa18 |
| $g_{\mathrm{raw,Na18}}$ anchor at 293.15 K | Temperature of empirical relative-current anchor (relative amplitude 0.82) | 293.15 | K | iNa18 |
| $g_{\mathrm{raw,Na18}}$ anchor at 303.15 K | Temperature of empirical relative-current anchor (relative amplitude 1.00) | 303.15 | K | iNa18 |
| $g_{\mathrm{Na18}}$ | Maximum Na$_V$1.8 conductance density | Set by fiber model | S cm$^{-2}$ | iNa18 |

All temperatures in Table S8 are absolute temperatures in K; voltage quantities are in mV; $Q_{10}$ factors, relative-amplitude factors, and gating variables are dimensionless. The empirical relative-amplitude anchors used by $g_{\mathrm{raw,Na18}}$ are 0.62 at 283.15 K, 0.82 at 293.15 K, and 1.00 at 303.15 K. These values define the piecewise scaling in Eq. 65 and correspond to 10 °C, 20 °C, and 30 °C, respectively.

**Table S9. K2P and active-node channel parameters**

TRAAK/TREK1 K2P parameters and active-node channel-density values used in the model.

Model scope: compartment model only.

K2P (TRAAK/TREK1) parameters

| Symbol | Description | Value | Unit | Module / notes | Reference |
|---|---|---|---|---|---|
| E_L | Leak/K2P reversal potential | -0.09 | V | Shared: Value shown as -0.09 V in model; used to form E_LT in K2P current. | (*42*) |
| I_scale_K2P | Scaling factor for K2P current density | 1 | 1 | Shared: Multiplier in K2P_node expression. | (*33*) |
| V_drv | Voltage scale in drv(x) | 0.2 | V | Shared: drv(x)=sign(x)*V_drv*(1-exp(-abs(x)/V_drv)). | (*33*) |
| g_TRAAK_node | Max conductance density for TRAAK component | 3 S cm⁻² | S cm⁻² | TRAAK: pre-low-temperature-response-calibration compartment-model value (30000 S m⁻²). | (*49*) |
| g_TREK1_node | Max conductance density for TREK1 component | 5 S cm⁻² | S cm⁻² | TREK1: pre-low-temperature-response-calibration compartment-model value (50000 S m⁻²). | (*49*) |
| tau_TRAAK0 | Baseline time constant in tau_TRAAK(T) | 0.004 | s | TRAAK: Model value: 0.004 s. | (*50*) |
| z_eff | Effective gating charge | 2.2 | 1 | TRAAK/TREK1: Used in O_inf and Oinf_act/deact. | (*50*) |
| Vd | Mid-voltage for kon/koff sigmoid | 0.04 | V | TRAAK: Model value: 0.04 V. | (*50*) |
| kd | Slope factor for kon/koff voltage dependence | 0.01 | V | TRAAK: Model value: 0.01 V. | (*50*) |
| kd0 | Max on-rate parameter in kon(Vm) | 10 | s⁻¹ | TRAAK: Model value: 10 s⁻¹. | (*50*) |
| kr0 | Max off-rate parameter in koff(Vm) | 10 | s⁻¹ | TRAAK: Model value: 10 s⁻¹. | (*50*) |
| dmu12_act | Half activation Δμ parameter in Oinf_act | 0.055 [V] | V | TREK1: Activation half-shift. | (*49*) |
| dmu12_deact | Half deactivation Δμ parameter in Oinf_deact | 0.035 [V] | V | TREK1: Deactivation half-shift. | (*49*) |
| tau_on | Base activation (depolarization) time constant | 0.005 [s] | s | TREK1: Temperature-scaled to tau_onT. | (*49*) |
| tau_off | Base deactivation (repolarization) time constant | 0.003 [s] | s | TREK1: Temperature-scaled to tau_offT. | (*49*) |
| dVt | Voltage-rate scale in speed term | 100 [V/s] | V/s | TREK1: Used in speed = d(Vm,t)/dVt. | (*49*) |

| Symbol | Description | Value | Unit | Module / notes | Reference |
|---|---|---|---|---|---|
| eps_dir | Heaviside smoothing parameter (numerical regularization) | 1e-3 | 1 | TREK1: Defined in parameter table; not explicitly used in shown w=tanh(speed) expression but retained for reproducibility. | (*33*) |
| O_AK(t0) | Initial value of O_AK | 0.05 | 1 | TRAAK: Initial time derivative set to 0. | (*33*) |
| D_AK(t0) | Initial value of D_AK | 0 | 1 | TRAAK: Initial time derivative set to 0. | (*33*) |
| O_TREK2(t0) | Initial value of O_TREK2 | 0.05 | 1 | TREK1: Initial time derivative set to 0. | (*33*) |

Active-node channel-density parameter sets

| Parameter | Value | Model family | Compartment | Software module | Low-temperature-response calibration state |
|---|---|---|---|---|---|
| gNa11 | 0.005 S $cm^{-2}$ | Generic compartment model | Active node | NEURON compartmental implementation | Before calibration |
| gNa16 | 0.375 S $cm^{-2}$ | Generic compartment model | Active node | NEURON compartmental implementation | Before calibration |
| g_TRAAK_node | 3.0 S $cm^{-2}$ | Generic compartment model | Active node | NEURON compartmental implementation | Before calibration |
| g_TREK1_node | 5.0 S $cm^{-2}$ | Generic compartment model | Active node | NEURON compartmental implementation | Before calibration |
| MRG-specific gNa11 | 0.0075 S $cm^{-2}$ | MRG-style compartment model | Active node | NEURON compartmental implementation | After calibration |
| MRG-specific gNa16 | 0.5625 S $cm^{-2}$ | MRG-style compartment model | Active node | NEURON compartmental implementation | After calibration |
| MRG-specific gTRAAK | 8.0 S $cm^{-2}$ | MRG-style compartment model | Active node | NEURON compartmental implementation | After calibration |
| MRG-specific gTREK1 | 24.0 S $cm^{-2}$ | MRG-style compartment model | Active node | NEURON compartmental implementation | After calibration |

The conductance values in Table S8 and the two active-node parameter sets in Table S9 are not interchangeable. Table S8 reports the EFM neuron model implementation. In Table S9, the generic compartment-model set records the values before low-temperature-response calibration, whereas the MRG-specific set records the values after calibration.

### Table S10. Bioheat and perfusion parameters

Blood perfusion, metabolic heat generation, and arterial blood-temperature parameters.

Bioheat/perfusion parameters

| Tissue/Material (–) | Blood density $\rho_b$ (kg m⁻³) | Blood specific heat capacity $C_b$ (J kg⁻¹ K⁻¹) | Blood perfusion rate $\omega_b$ (s⁻¹) | Metabolic heat generation $Q_{met}$ (W m⁻³) | Arterial blood temperature $T_b$ (K) | References ($\rho_b$, $C_b$, $\omega_b$, $Q_{met}$, $T_b$) |
|---|---|---|---|---|---|---|
| Muscle | 1060 | 3600 | 0.0015 | 3500 | 310.15 | (*69*, *76*) |
| epidural | 1060 | 3600 | 0.003 | 1500 | 310.15 | (*69*, *76*) |
| perineurium | 1060 | 3600 | 0.0 | 500 | 310.15 | (*69*, *76*) |
| nerve_fiber | 1060 | 3600 | 0.001 | 4000 | 310.15 | (*69*, *76*) |

**Table S11. Definitions of principal physical field variables**

Definitions of the five principal field variables used in the thermal-fluid and electrical-field equations. Numerical parameter values remain in Tables S2 to S10.

| Symbol | Definition | SI unit | Type | Applicable equations/module |
|---|---|---|---|---|
| $\mathbf{u}$ | Fluid velocity vector | m s⁻¹ | Field variable | Eqs. 1–3; thermal-fluid module |
| $p$ | Fluid pressure | Pa | Field variable | Eq. 2; thermal-fluid module |
| $T$ | Absolute temperature | K | Field/state variable | Eqs. 3–7 and temperature-dependent neural equations; thermal and neural modules |
| $V$ | Electric potential | V | Field variable | Eqs. 9–11; electrical-field module |
| $\mathbf{E}$ | Electric-field vector | V m⁻¹ | Derived field variable | Eqs. 8–10; electrical-field module |

### Materials availability

This study did not generate unique biological reagents. Custom-designed cuff devices, associated assembly components, and related non-commercial experimental materials described in this study are available from the lead contact upon reasonable request, subject to institutional policies and material-transfer requirements where applicable.

### Data and code availability

Data, analysis code, and simulation resources are being curated in a private GitHub repository at https://github.com/yangshu717/multimodal_NeuroSwitch. Access will be provided to editors and reviewers upon request. The repository will be made public upon publication and archived as a versioned release with a permanent DOI. NeuroSwitch devices and fabrication files are available from the lead contact upon reasonable request.

**References**


1. S. S. Casagrande, A. Z. Beccera, K. F. Rust, C. C. Cowie, Opioid prescription and diabetes among Medicare beneficiaries. *Diabetes Res Clin Pract* **196**, 110240 (2023).
2. T. H. Nost *et al.*, Prevalence of substance use disorder diagnoses in patients with chronic pain receiving reimbursed opioids: An epidemiological study of four Norwegian health registries. *Scand J Pain* **24**, 20240059 (2024).
3. N. D. Volkow, D. L. Longo, A. T. McLellan, Opioid Abuse in Chronic Pain — Misconceptions and Mitigation Strategies. *New England Journal of Medicine* **374**, 1253–1263 (2016).
4. Y. J. Kim *et al.*, Wireless and bioresorbable triboelectric nerve block system for postoperative pain control. *Nature Biomedical Engineering*. 2026 (https://doi.org/10.1038/s41551-025-01579-2).
5. G. Lee *et al.*, A bioresorbable peripheral nerve stimulator for electronic pain block. *Sci Adv* **8**, eabp9169 (2022).
6. J. T. Reeder *et al.*, Soft, bioresorbable coolers for reversible conduction block of peripheral nerves. *Science* **377**, 109–115 (2022).
7. C.-E. Wong *et al.*, Sciatic nerve stimulation alleviates acute neuropathic pain via modulation of neuroinflammation and descending pain inhibition in a rodent model. *Journal of Neuroinflammation* **19**, 153 (2022).
8. C. E. Wong *et al.*, Sciatic nerve stimulation alleviates neuropathic pain and associated neuroinflammation in the dorsal root ganglia in a rodent model. *J Transl Med* **22**, 770 (2024).
9. K. L. Kilgore, N. Bhadra, Reversible nerve conduction block using kilohertz frequency alternating current. *Neuromodulation* **17**, 242–254; discussion 254–255 (2014).
10. S. Liu *et al.*, Neural basis of transcutaneous electrical nerve stimulation for neuropathic pain relief. *Neuron* **113**, 3616–3631.e6 (2025).
11. D. D. Price, Dorsal horn neuronal responses and quantitative sensory testing help explain normal and abnormal pain. *Pain* **154**, 1161–1162 (2013).
12. X. Navarro *et al.*, A critical review of interfaces with the peripheral nervous system for the control of neuroprostheses and hybrid bionic systems. *Journal of the Peripheral Nervous System* **10**, 229–258 (2005).
13. N. Jayaprakash *et al.*, Organ- and function-specific anatomical organization of vagal fibers supports fascicular vagus nerve stimulation. *Brain Stimul* **16**, 484–506 (2023).
14. U. Latif *et al.*, Consensus Guidelines for the Use of Peripheral Nerve Stimulation in the Treatment of Chronic Pain and Neurological Diseases: A Neuron Project from the American Society of Pain and Neuroscience. *J Pain Res* **18**, 5949–5990 (2025).
15. D. M. Ackermann, Jr., N. Bhadra, E. L. Foldes, X. F. Wang, K. L. Kilgore, Effect of nerve

cuff electrode geometry on onset response firing in high-frequency nerve conduction block. *IEEE Trans Neural Syst Rehabil Eng* **18**, 658–665 (2010).
16. D. M. Ackermann, N. Bhadra, M. Gerges, P. J. Thomas, Dynamics and sensitivity analysis of high-frequency conduction block. *J Neural Eng* **8**, 065007 (2011).
17. C. A. Miller, P. J. Abbas, K. V. Nourski, N. Hu, B. K. Robinson, Electrode configuration influences action potential initiation site and ensemble stochastic response properties. *Hearing Research* **175**, 200–214 (2003).
18. K. P. Cheng *et al.*, Application of kilohertz-frequency block to mitigate off-target motor effects of vagus nerve stimulation in swine. *Nat Commun* **17**, 1066 (2025).
19. C. Tai, J. Wang, M. B. Chancellor, J. R. Roppolo, W. C. de Groat, Influence of temperature on pudendal nerve block induced by high frequency biphasic electrical current. *J Urol* **180**, 1173–8 (2008).
20. U. Ahmed *et al.*, Anodal block permits directional vagus nerve stimulation. *Sci Rep* **10**, 9221 (2020).
21. C. van den Honert, J. T. Mortimer, Generation of unidirectionally propagated action potentials in a peripheral nerve by brief stimuli. *Science* **206**, 1311–2 (1979).
22. F. Ciotti *et al.*, Towards enhanced functionality of vagus neuroprostheses through in silico optimized stimulation. *Nat Commun* **15**, 6119 (2024).
23. D. M. Ackermann, Jr., E. L. Foldes, N. Bhadra, K. L. Kilgore, Effect of bipolar cuff electrode design on block thresholds in high-frequency electrical neural conduction block. *IEEE Trans Neural Syst Rehabil Eng* **17**, 469–77 (2009).
24. Y. A. Patel, R. J. Butera, Differential fiber-specific block of nerve conduction in mammalian peripheral nerves using kilohertz electrical stimulation. *J Neurophysiol* **113**, 3923–9 (2015).
25. N. Bhadra, K. L. Kilgore, High-frequency electrical conduction block of mammalian peripheral motor nerve. *Muscle Nerve* **32**, 782–90 (2005).
26. M. Franke *et al.*, Combined KHFAC + DC nerve block without onset or reduced nerve conductivity after block. *J Neural Eng* **11**, 056012 (2014).
27. F. Lembeck, P. Holzer, Substance P as neurogenic mediator of antidromic vasodilation and neurogenic plasma extravasation. *Naunyn Schmiedebergs Arch Pharmacol* **310**, 175–83 (1979).
28. I. M. Chiu, C. A. von Hehn, C. J. Woolf, Neurogenic inflammation and the peripheral nervous system in host defense and immunopathology. *Nature Neuroscience* **15**, 1063–1067 (2012).
29. S. D. Brain, T. J. Williams, J. R. Tippins, H. R. Morris, I. MacIntyre, Calcitonin gene-related peptide is a potent vasodilator. *Nature* **313**, 54–6 (1985).
30. W. Janig, S. J. Lisney, Small diameter myelinated afferents produce vasodilatation but not plasma extravasation in rat skin. *J Physiol* **415**, 477–86 (1989).
31. R. A. Rietmeijer, B. Sorum, B. Li, S. G. Brohawn, Physical basis for distinct basal and mechanically gated activity of the human K(+) channel TRAAK. *Neuron* **109**, 2902–2913.e4 (2021).
32. S. G. Brohawn *et al.*, The mechanosensitive ion channel TRAAK is localized to the mammalian node of Ranvier. *Elife* **8**, e50403 (2019).
33. H. Kanda, S. Tonomura, J. G. Gu, Effects of Cooling Temperatures via Thermal K2P Channels on Regeneration of High-Frequency Action Potentials at Nodes of Ranvier of Rat Abeta-Afferent Nerves. *eNeuro* **8**, ENEURO.0308–21.2021 (2021).

34. K. Zimmermann *et al.*, Sensory neuron sodium channel Nav1.8 is essential for pain at low temperatures. *Nature* **447**, 855–8 (2007).
35. J. R. Schwarz, The effect of temperature on Na currents in rat myelinated nerve fibres. *Pflugers Arch* **406**, 397–404 (1986).
36. S. G. Lomber, B. R. Payne, J. A. Horel, The cryoloop: an adaptable reversible cooling deactivation method for behavioral or electrophysiological assessment of neural function. *J Neurosci Methods* **86**, 179–94 (1999).
37. P. C. Petersen, G. Buzsaki, Cooling of Medial Septum Reveals Theta Phase Lag Coordination of Hippocampal Cell Assemblies. *Neuron* **107**, 731–744.e3 (2020).
38. P. Borgdorff, P. G. Versteeg, An implantable nerve cooler for the exercising dog. *Eur J Appl Physiol Occup Physiol* **53**, 175–9 (1984).
39. T. Morgan *et al.*, Thermal block of mammalian unmyelinated C fibers by local cooling to 15-25 degrees C after a brief heating at 45 degrees C. *J Neurophysiol* **123**, 2173–2179 (2020).
40. D. M. Ackermann, E. L. Foldes, N. Bhadra, K. L. Kilgore, Nerve conduction block using combined thermoelectric cooling and high frequency electrical stimulation. *J Neurosci Methods* **193**, 72–6 (2010).
41. C. C. H. Cohen *et al.*, Saltatory Conduction along Myelinated Axons Involves a Periaxonal Nanocircuit. *Cell* **180**, 311–322.e15 (2020).
42. C. C. McIntyre, A. G. Richardson, W. M. Grill, Modeling the excitability of mammalian nerve fibers: influence of afterpotentials on the recovery cycle. *J Neurophysiol* **87**, 995–1006 (2002).
43. S. F. Lempka, C. C. McIntyre, K. L. Kilgore, A. G. Machado, Computational analysis of kilohertz frequency spinal cord stimulation for chronic pain management. *Anesthesiology* **122**, 1362–76 (2015).
44. E. D. Musselman, N. A. Pelot, W. M. Grill, Validated computational models predict vagus nerve stimulation thresholds in preclinical animals and humans. *J Neural Eng* **20**, 036032 (2023).
45. M. A. Hussain, W. M. Grill, N. A. Pelot, Highly efficient modeling and optimization of neural fiber responses to electrical stimulation. *Nat Commun* **15**, 7597 (2024).
46. D. P. Marshall, E. S. Farah, E. D. Musselman, N. A. Pelot, W. M. Grill, PyFibers: An open-source NEURON-Python package to simulate responses of model nerve fibers to electrical stimulation. *PLoS Comput Biol* **21**, e1013764 (2025).
47. C. Boehler, S. Carli, L. Fadiga, T. Stieglitz, M. Asplund, Tutorial: guidelines for standardized performance tests for electrodes intended for neural interfaces and bioelectronics. *Nat Protoc* **15**, 3557–3578 (2020).
48. K. G. Beam, P. L. Donaldson, A quantitative study of potassium channel kinetics in rat skeletal muscle from 1 to 37 degrees C. *J Gen Physiol* **81**, 485–512 (1983).
49. H. Kanda *et al.*, TREK-1 and TRAAK Are Principal K(+) Channels at the Nodes of Ranvier for Rapid Action Potential Conduction on Mammalian Myelinated Afferent Nerves. *Neuron* **104**, 960–971.e7 (2019).
50. M. Schewe *et al.*, A Non-canonical Voltage-Sensing Mechanism Controls Gating in K2P K(+) Channels. *Cell* **164**, 937–49 (2016).
51. M. Volgushev, T. R. Vidyasagar, M. Chistiakova, T. Yousef, U. T. Eysel, Membrane properties and spike generation in rat visual cortical cells during reversible cooling. *J Physiol* **522 Pt 1**, 59–76 (2000).

52. B. Bromm, Spike frequency of the nodal membrane generated by high-frequency alternating current. *Pflugers Arch* **353**, 1–19 (1975).
53. E. D. Musselman, J. E. Cariello, W. M. Grill, N. A. Pelot, ASCENT (Automated Simulations to Characterize Electrical Nerve Thresholds): A pipeline for sample-specific computational modeling of electrical stimulation of peripheral nerves. *PLOS Computational Biology* **17**, e1009285 (2021).
54. M. L. Hines, N. T. Carnevale, The NEURON simulation environment. *Neural Comput* **9**, 1179–209 (1997).
55. C. A. Bossetti, M. J. Birdno, W. M. Grill, Analysis of the quasi-static approximation for calculating potentials generated by neural stimulation. *J Neural Eng* **5**, 44–53 (2008).
56. A. L. Hodgkin, A. F. Huxley, A quantitative description of membrane current and its application to conduction and excitation in nerve. *The Journal of Physiology* **117**, 500–544 (1952).
57. E. Peña, N. A. Pelot, W. M. Grill, Computational models of compound nerve action potentials: Efficient filter-based methods to quantify effects of tissue conductivities, conduction distance, and nerve fiber parameters. *PLOS Computational Biology* **20**, e1011833 (2024).
58. R. V. Shannon, A model of safe levels for electrical stimulation. *IEEE Transactions on Biomedical Engineering* **39**, 424–426 (1992).
59. S. F. Cogan, Neural stimulation and recording electrodes. *Annu Rev Biomed Eng* **10**, 275–309 (2008).
60. G. Yi, W. M. Grill, Kilohertz waveforms optimized to produce closed-state Na+ channel inactivation eliminate onset response in nerve conduction block. *PLOS Computational Biology* **16**, e1007766 (2020).
61. E. Peña, N. A. Pelot, W. M. Grill, Non-monotonic kilohertz frequency neural block thresholds arise from amplitude- and frequency-dependent charge imbalance. *Scientific Reports* **11**, 5077 (2021).
62. E. Peña, N. A. Pelot, W. M. Grill, Spatiotemporal parameters for energy efficient kilohertz-frequency nerve block with low onset response. *Journal of NeuroEngineering and Rehabilitation* **20**, 72 (2023).
63. M. C. Kiernan, Effects of temperature on the excitability properties of human motor axons. *Brain* **124**, 816–825 (2001).
64. L. Rueda-Ruzafa, S. Herrera-Pérez, A. Campos-Ríos, J. A. Lamas, Are TREK Channels Temperature Sensors? *Frontiers in Cellular Neuroscience* **15**, 744702 (2021).
65. F. Maingret, M. Fosset, F. Lesage, M. Lazdunski, E. Honoré, TRAAK Is a Mammalian Neuronal Mechano-gated K+ Channel. *Journal of Biological Chemistry* **274**, 1381–1387 (1999).
66. F. Maingret *et al.*, TREK-1 is a heat-activated background K+ channel. *The EMBO Journal* **19**, 2483–2491 (2000).
67. J. Noël *et al.*, The mechano-activated K+ channels TRAAK and TREK-1 control both warm and cold perception. *The EMBO Journal* **28**, 1308–1318 (2009).
68. S. G. Brohawn, E. B. Campbell, R. MacKinnon, Physical mechanism for gating and mechanosensitivity of the human TRAAK K+ channel. *Nature* **516**, 126–130 (2014).
69. C. Baumgartner *et al.* (IT'IS Foundation, 2025).
70. N. A. Pelot, C. E. Behrend, W. M. Grill, On the parameters used in finite element modeling of compound peripheral nerves. *J Neural Eng* **16**, 016007 (2019).

71. S. Gabriel, R. W. Lau, C. Gabriel, The dielectric properties of biological tissues: III. Parametric models for the dielectric spectrum of tissues. *Phys Med Biol* **41**, 2271–93 (1996).
72. H. Ye, J. Ng, Shielding effects of myelin sheath on axolemma depolarization under transverse electric field stimulation. *PeerJ* **6**, e6020 (2018).
73. C. Johnson, W. R. Holmes, A. Brown, P. Jung, Minimizing the caliber of myelinated axons by means of nodal constrictions. *J Neurophysiol* **114**, 1874–84 (2015).
74. C. H. Berthold, M. Rydmark, Electrophysiology and morphology of myelinated nerve fibers. VI. Anatomy of the paranode-node-paranode region in the cat. *Experientia* **39**, 964–76 (1983).
75. S. Y. Chiu, J. M. Ritchie, R. B. Rogart, D. Stagg, A quantitative description of membrane currents in rabbit myelinated nerve. *J Physiol* **292**, 149–66 (1979).
76. H. H. Pennes, Analysis of tissue and arterial blood temperatures in the resting human forearm. *J Appl Physiol* **1**, 93–122 (1948).
